# Breakdown and repair of the aging brain metabolic system

**DOI:** 10.1101/2023.08.30.555341

**Authors:** Polina Shichkova, Jay S. Coggan, Lida Kanari, Elvis Boci, Cyrille Favreau, Stefano Maximiliano Antonel, Daniel Keller, Henry Markram

## Abstract

The study presented explores the complex relationship between the aging brain, energy metabolism, blood flow and neuronal activity by introducing a comprehensive, data-driven molecular model of the neuro-glial vascular system, including all key enzymes, transporters, metabolites, and blood flow vital for neuronal electrical activity with 16’800 interaction pathways. We find significant alterations in metabolite concentrations and differential effects on ATP supply in neurons and astrocytes and within subcellular compartments within aged brains, and identify reduced Na^+^/K^+^-ATPase as the leading cause of impaired neuronal action potentials. The model predicts that the metabolic pathways cluster more closely in the aged brain, suggesting a loss of robustness and adaptability. Additionally, the aged metabolic system displays reduced flexibility, undermining its capacity to efficiently respond to stimuli and recover from damage. Through transcription factor analysis, the estrogen-related receptor alpha (ESRRA) emerged as a central target connected to these aging-related changes. An unguided optimization search pinpointed potential interventions capable of restoring the brain’s metabolic flexibility and restoring action potential generation. These strategies include increasing the NADH cytosol-mitochondria shuttle, NAD+ pool, ketone β-hydroxybutyrate, lactate and Na^+^/K^+^-ATPase and reducing blood glucose levels. The model is open-sourced to help guide further research in brain metabolism.

## Main

### Mechanisms of brain aging

The risk of numerous disorders, including neurodegenerative diseases, increases dramatically with age (Niccoli and Partridge, 2012; Hou et al., 2019). At the core of brain aging lies energy metabolism (López-Otín et al., 2013, 2023; Mattson and Arumugam, 2018). Neuronal activity is energetically demanding, requiring substantial amounts of ATP, as reflected in the brain’s disproportionate oxygen and glucose consumption compared to the rest of the body (Kety, 1957; Mink et al., 1981; Sokoloff, 1996; Rolfe and Brown, 1997). Metabolic support and neuronal activity are closely linked (Mann et al., 2021), suggesting that age-related loss of metabolic support impacts the generation of electrical activity in the brain. But the vast number of biochemical reactions forming the metabolic system constitute a highly complex system that makes it exceedingly difficult to isolate how changes in the metabolic system impact neuronal activity.

Various dynamic models of brain metabolism have been developed over the decades. Early models (Aubert et al., 2001; Cloutier et al., 2009) focused on core components of the metabolic system and generalized many processes such as mitochondrial respiration. Recent models have incorporated more detailed descriptions for selected subsystems, such as the pentose phosphate pathway (Winter et al., 2018), mitochondrial metabolism (Berndt et al., 2015; Theurey et al., 2019) or neuronal electrophysiology (Jolivet et al., 2015). While well-validated and suited for the target questions possible with these models, a considerably more biologically detailed model is required to address more complex questions such as how age-related changes in metabolism impact action potential generation and responses to stimuli.

The model we developed brings the previous models together with greater detail and adds previously omitted subsystems such as glutathione metabolism and regulation of glycogenolysis. The model also extends previous models by coupling the metabolic system to the intricate cellular processes underlying action potential generation such as the Na^+^/K^+^-ATPase pump, the glutamate-glutamine cycle, and ATP production by mitochondria and cytosol. The model allows, for the first time, a view on how electrical activity impacts the metabolic system and vice versa. Finally, the model also integrates blood flow and dynamic exchanges between the vasculature and the neurons and glia to yield a comprehensive view of the biochemical network operating across the Neuro-Glia-Vascular (NGV) system. The model therefore also allows addressing questions related to nutrient supply to the brain.

Since there is a vast amount of data related to brain metabolism in the literature and databases, we adopted a strict data-driven strategy to constrain the construction of the biochemical network of the NGV system. This strategy allowed us to use data relevant to the young and aged brain’s metabolic system and reconstruct and simulate their respective metabolic systems. The model integrates all key metabolites, transporters and enzymes with all key cellular and extracellular processes underlying neuronal firing and interactions with the blood (Fig. 1). Subcompartments such as mitochondrial matrix and intermembrane space, cytosol in neurons and astrocytes, blood flow bringing nutrients and oxygen in capillaries, endothelium, and the extracellular space (interstitium and basal lamina) are captured using their data-driven volumes (see the model at https://bbp.epfl.ch/ngv-portal-dev/), allowing modeling of their cross-compartment processes such as transport and exchange. The model does not capture metabolic waste management, such as lactate removal. Concentrations of molecules are specified in molar units (mM) and fluxes of reactions and transport processes are given in molar concentrations per second (mM/second).

**Fig. 1.**
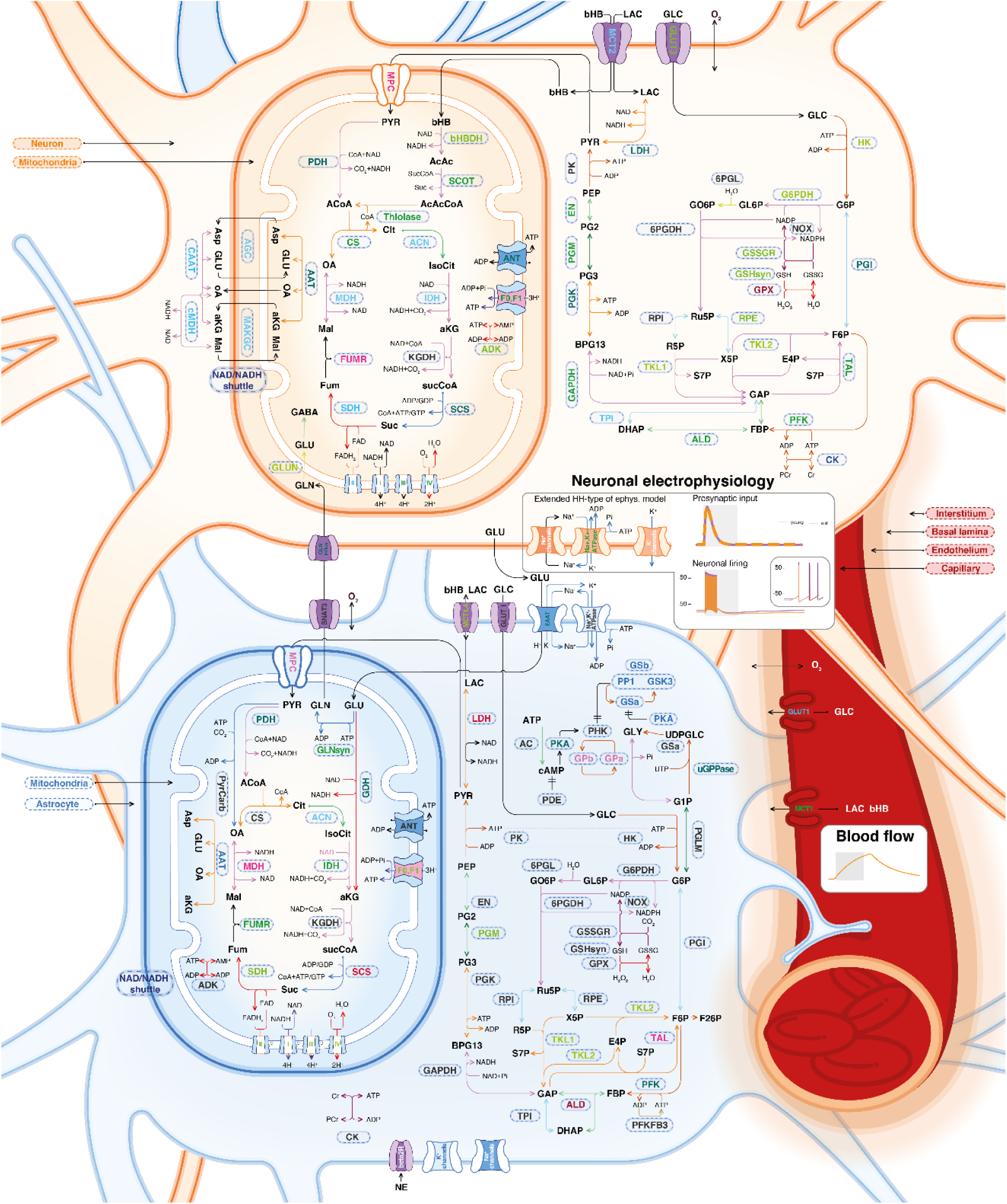
Model overview. The model consists of three connected sub-systems: metabolism, neuronal electrophysiology and the blood flow. Compartments of the model include the neuronal and astrocytic cytosol, mitochondrial matrix and intermembrane space, interstitium, basal lamina, endothelium, capillary, artery (only with fixed arterial concentrations of nutrients and oxygen), and endoplasmic reticulum (only with Ca^2+^ fixed pool). Enzymes and transporters shown in the figure correspond to the rate equations in the model which govern the dynamics of metabolite concentration changes. Neuronal electrophysiology is modeled in a slightly extended Hodgkin-Huxley type of model. Blood flow activation is described by a simple function dependent on the stimulus onset and duration according to the literature models.

We validated the model extensively against a corpus of data reported in the literature on how enzyme and transporter activities and metabolite concentrations change in response to stimulation that were not used to construct the model model (Supplementary Fig. 1, Supplementary Table 1). For example, the model shows that Na^+^/K^+^ pump ATP use in the astrocyte is comparable with that of the neuron (Fig. 3g) consistent with recent evidence (Barros, 2022). In line with previous studies (Bélanger et al., 2011), mitochondrial ATP production as a share of total ATP production is higher in neurons than in astrocytes, at 84% versus 70% (Supplementary Fig. 2), closely matching experimental estimates of 75% in astrocytes (Bouzier-Sore et al., 2006; Barros, 2022). These data emerged when the model was simulated and their consistency with a range of reported experimental data suggests that the model accurately captures the most essential elements of the brain’s metabolic system.

**Fig. 2.**
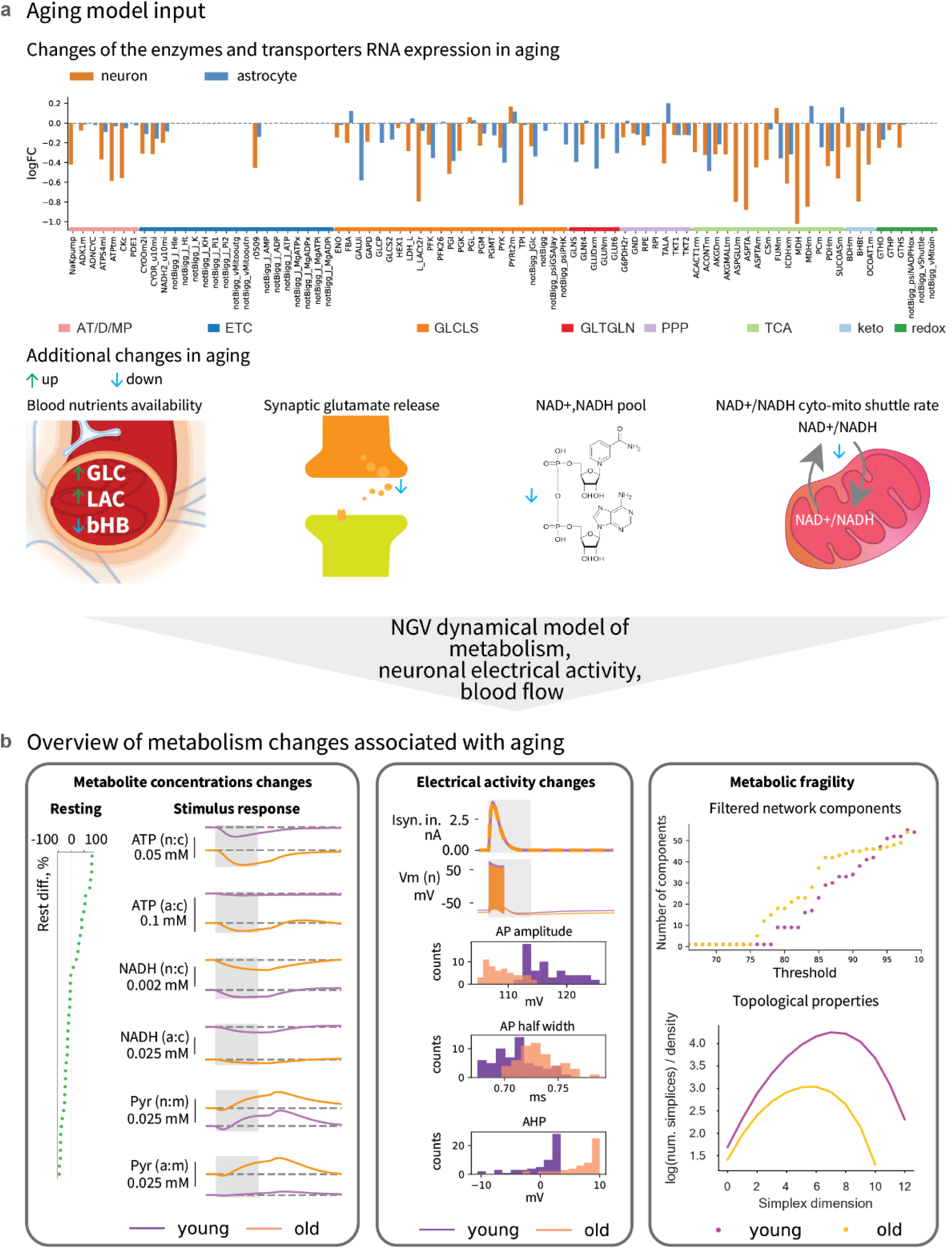
Aging model input (a) and results overview (b). **a,** Aging input is modeled with RNA expression fold changes of enzymes and transporters, scaling of arterial glucose, lactate and b-hydroxybutyrate, as well as the total NAD (reduced and oxidized) pool, synaptic effects of glutamate concentration changes upon release events, and the reducing equivalents (NADH-related) shuttle between cytosol and mitochondria. **b,** The key results include aging effects on metabolite levels, electrical activity of the neurons, and changes in adaptivity of the system in response to kinetic parameter perturbations (mimicking molecular damage and other conditions affecting enzyme and transporter functions).

**Fig. 3.**
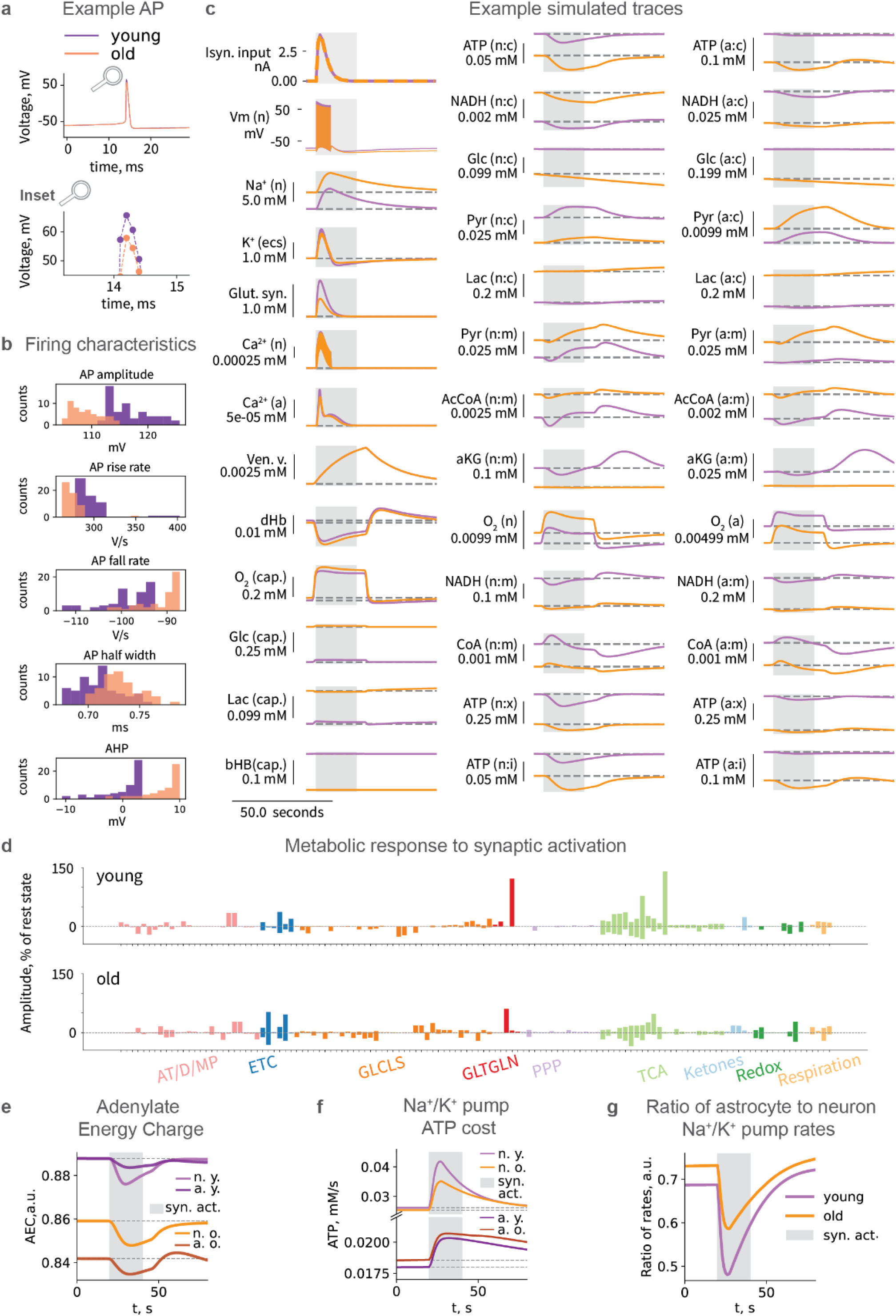
Simulation results. **a,** Example AP in voltage traces in simulations of young and aged neurons with insets providing a closer view. **b,** Characteristics of neuronal firing in young and old ages upon synaptic activation. **c,** Dynamics of metabolism in response to synaptic activation at different ages (only a selection of the most important variables is shown). Compartment names abbreviations: n - neuron, a - astrocyte, c - cytosol, m - mitochondria, x - mitochondrial matrix, i - mitochondrial IMS, cap. - capillary. **d,** Amplitude of concentration changes in response to synaptic activation in young (top) and old (bottom), individual metabolites labels are available in **Supplementary Fig. 18. e,** AEC: Adenylate Energy Charge in young and old neurons and astrocytes (AEC = (ATP + 0.5ADP)/(ATP + ADP + AMP)). **f,** Main energy consumption: Na^+^/K^+^-ATPase rate of ATP use. **g,** Ratio of astrocyte to neuron Na^+^/K^+^ pump rate.

Alterations in enzyme expression have recently been shown to actively contribute to tissue aging and therefore potential drug targets to counter aging (Palla et al., 2021). To model aging of NGV metabolism, we therefore used RNA expression changes (RNA FCs) from a comprehensive study on mouse cell-type changes (Schaum et al., 2020; Zhang et al., 2021a) to scale enzyme and transporter concentrations. These concentrations determine the output from their corresponding reaction/transport rate equations. After applying the RNAseq data (Schaum et al., 2020; Zhang et al., 2021a) to their respective metabolic pathways, we obtained a map of the decrease in expression of most enzymes with aging in both neurons and astrocytes. The map revealed, for example, that succinate dehydrogenase (SDH) is differentially affected by aging in neurons and astrocytes. SDH is a mitochondrial energy nexus and serves as complex II of the mitochondrial electron transport chain (ETC). SDH connects the tricarboxylic acid cycle (TCA) to the ETC. This result indicates that pre- and post-SDH enzymes of TCA (fumarase and succinate CoA ligase) display opposite changes in aged neurons and astrocytes. SDH itself decreases more in aged neurons than in aged astrocytes. In neurons, aging reduces both succinate CoA ligase and SDH, while increasing fumarase. Unlike in neurons, succinate CoA ligase levels rise in astrocytes during aging. SDH decreases slightly while fumarase levels decline more.

In addition to changes in enzyme and transporter expression, we used published values to adjust arterial glucose, lactate, β-hydroxybutyrate levels, the total NAD (reduced and oxidized) pool, as well as glutamate concentration changes caused by synaptic transmission (Dong and Brewer, 2019; Cox et al., 2022). The young brain’s metabolic system is in an equilibrium at rest - i.e. when no stimulus is applied. To be able to compare the young and aged models we could ensure that the aged system was also at steady state by reducing the NADH shuttle capacity between the cytosol and mitochondria. All aging data applied to the model are summarized in Fig. 2, with further details available in the Methods section. When we simulated the dynamics of this complex system, driven by either synaptic input or current injection that generated action potentials, we observed numerous age-specific differences that are consistent with prior reports (Supplementary Table 1) further validating the model and providing a spectrum of new insights into how the NGV metabolic system may age.

### Aging affects metabolite levels at rest and during stimuli

The simulated aging phenotype exhibits a distinct resting state profile of metabolite concentrations when compared to that of the young brain (Supplementary Fig. 3a), the change in concentrations in response to stimuli also differs (Fig. 3c, d, Supplementary Fig. 3b, 4, 5d, 6a), but the changes in responses to stimuli of varying amplitudes are not uniform across different metabolites (Supplementary Fig. 6, 7). We performed Uniform Manifold Approximation and Projection for Dimension Reduction (UMAP) dimensionality reduction on relative differences in concentration traces between the two ages, and observed numerous interdependencies between pathways. The pentose phosphate pathway (PPP) and TCA tend to form pathway-related clusters (Supplementary Fig. 8). Moreover, pairwise Kendall correlation between metabolic concentration temporal profiles is also affected by aging (Supplementary Fig. 9). This effect may be caused by widely described metabolic dysregulation in aging (Mattson and Arumugam, 2018). Reaction and transport fluxes are impacted as well (Supplementary Fig. 10-12). Aging effects on metabolite concentrations at rest and in response to stimuli are therefore metabolite-specific and largely uncorrelated, indicative of a fragmentation of the metabolic network in aging.

### Lactate transport directionality changes in the aging metabolic system

In the aged metabolic system, neuronal lactate import is lower, while astrocyte lactate export is slightly higher. This effect can be explained by mitochondrial hypometabolism, which results in increased pyruvate levels and correspondingly higher levels of lactate. To examine the dependence of lactate transport directionality upon glucose levels in aged and young metabolic systems (Supplementary Fig. 13), we simulated the effects of varying resting blood glucose levels in a range of 1.6 to 14.6 mM with increments of 1 mM.

In the young system, we observed the expected astrocyte-to-neuron lactate shuttle (ANLS) at all tested blood glucose levels; as blood glucose levels increase, lactate export from astrocytes rises while lactate import to neurons decreases. This directionality is consistent with higher glucose availability reducing the need for neurons to import lactate and more lactate is available in and exported from astrocytes. In the aged metabolic system, the lactate shuttle has the same directionality as in the young system for moderate blood glucose levels (6.45 to 10.6 mM), consistent with a recent publication (Acevedo et al., 2023), but both neurons and astrocytes export lactate when glucose levels are low-to-normal (1.6 to 5.6 mM) and both neurons and astrocytes import lactate when glucose levels are high (11.6 to 14.6 mM). A possible explanation for this dysregulation in the aged metabolic system could involve NAD^+^/NADH and ATP/ADP ratios due to their regulatory role over the entire metabolic network, but this counterintuitive prediction requires experimental verification.

### Aging-associated changes in metabolism alter electrophysiological characteristics

We show for the first time how the aging metabolic system leads to changes in the generation of action potential by both synaptic input (Fig. 3) and current injection (Supplementary Fig. 5). Age-related differences in neuronal firing characteristics evoked by current injection are particularly important because it excludes the metabolic demand caused by glutamate release. We found similar changes in metabolic profiles following synaptic input and current injection (Supplementary Fig. 14), suggesting that metabolic changes mostly impact the action potential generation ability of neurons. A more detailed molecular coupling between the metabolic system and the entire glutamate cycle would however need to be included in the model to strengthen this prediction.

We found that changes in action potential shape and size are caused by a reduction in Na^+^/K^+^-ATPase expression in the aged brain, supporting a recent theory of non-canonical control of neuronal energy status (Baeza-Lehnert et al., 2019). To better understand whether other aspects of the metabolic system, such as reduced supply of ATP, also contribute to these changes, we increased the Na^+^/K^+^-ATPase expression levels in the aged brain model to the same as in the young brain while leaving all other aspects of the aging metabolic system in their aged state. There were no significant differences in action potentials at low frequencies (4-8 Hz) and only slight changes at much high-frequencies (78-79 Hz) suggesting that the decreased expression of the Na^+^/K^+^-ATPase pump is the main factor impairing the ability of neurons to generate action potentials. It is however still possible that other aspects of the NGV metabolic network become more important after sustained neuronal activity such as those during intense cognitive demand.

### Lower supply and demand for energy in the aged brain

Energy deficiency is a prominent hypothesis in brain aging (Bonvento and Bolaños, 2021), but it is not clear if the supply is limiting and/or demand is reduced and whether astrocytes and neurons are impacted in the same way. Adenylate energy charge (AEC), a widely used proxy for cellular energy availability (Atkinson, 1968), is higher in the young compared to the old state (Fig. 3e). However, this value does not separate supply from demand. To separate the two factors, we first computed the total ATP cost of firing action potentials. We find that the young brain model consumes approximately 2 billion ATP molecules per second per NGV unit (where one unit is one neuron, one astrocyte and their associated extracellular matrix and capillaries) at 8 Hz firing, and the aged brain model consumes around 1.8 billions molecules per second, which aligns well with literature estimates (Howarth et al., 2012; Yi and Grill, 2019; Zhu et al., 2019). We find that ATP production is lower in aged cytosol of both neuron and astrocytes, and in aged neuronal mitochondria as shown in Supplementary Fig. 2, but if one considers the reduced ATP-consumption (Fig. 3f) because of the lower levels of Na^+^/K^+^-ATPase (Niven, 2016; Meyer et al., 2022), then ATP supply is not limiting for the Na^+^/K^+^-ATPase that is present. While reduced ATP supply does not seem to be limiting action potential generation in the acute state, it remains possible that a persistently lower supply of ATP may still be the cause of a loss of Na^+^/K^+^-ATPase expression and hence impaired action potential generation over a longer period.

We also find that neurons and astrocytes are differentially impacted with age. Normally, astrocytic Na^+^/K^+^-ATPases consume slightly less than ⅔ of ATP of the neuronal Na^+^/K^+^-ATPases (Fig. 3g). In astrocytes, the ATP supply is only reduced in the cytosol and not in the mitochondria and the catalytic subunit of the Na^+^/K^+^-ATPases expression is unchanged with aging. While ATP consumption of the Na^+^/K^+^-ATPase pump in neurons decreases with aging (Fig. 3f), it slightly increases in astrocytes resulting in an increase in the ratio of astrocyte to neuron Na^+^/K^+^-ATPases ATP consumption from around 0.69 in the young brain to around 0.72 with aging. Since astrocytes do not need to fire action potentials, this finding suggests that there is an increase in the demand placed on astrocytes to support the neurons to clear extracellular K^+^ in order to help neurons generate their action potentials.

### Aging brain metabolism is more fragile and susceptible to damage

Protein dysfunction is associated with several aging hallmarks, including loss of proteostasis, oxidative damage, and impaired DNA repair (Mattson and Arumugam, 2018; Schaum et al., 2020). Moreover, reduced fidelity of protein translation leads to a phenotype resembling an early stage of Alzheimer’s disease (Brilkova et al., 2022). To mimic molecular damage and simulate the effect on enzyme and transporter functions, we introduced one perturbation at a time for each protein’s kinetic parameter (Michaelis constant, inhibition and activation constants, catalytic rate constant, i.e. a parameter in the enzyme rate equation), adjusting its value by 20% (increasing or decreasing in separate simulations). We then calculated the changes in the response of all metabolites to obtain a measure of how sensitive their concentrations are to the individual perturbations. We ran 2,264 simulations with perturbed parameters to measure metabolite sensitivities at rest and during a stimulus for the young and aged systems (see formula in Fig. 4b). The difference between the resting and stimulated state’s sensitivities, normalized by the resting state sensitivities, yielded a rest-normalized sensitivity. Since a larger value for a metabolite implies that a stimulus produces a larger change in its concentration as compared to rest when another parameter in the system is perturbed, we interpret such a change as the ability to adapt to damage of the system, and therefore call this metric, “metabolic adaptability” (Fig. 4d).

**Fig. 4.**
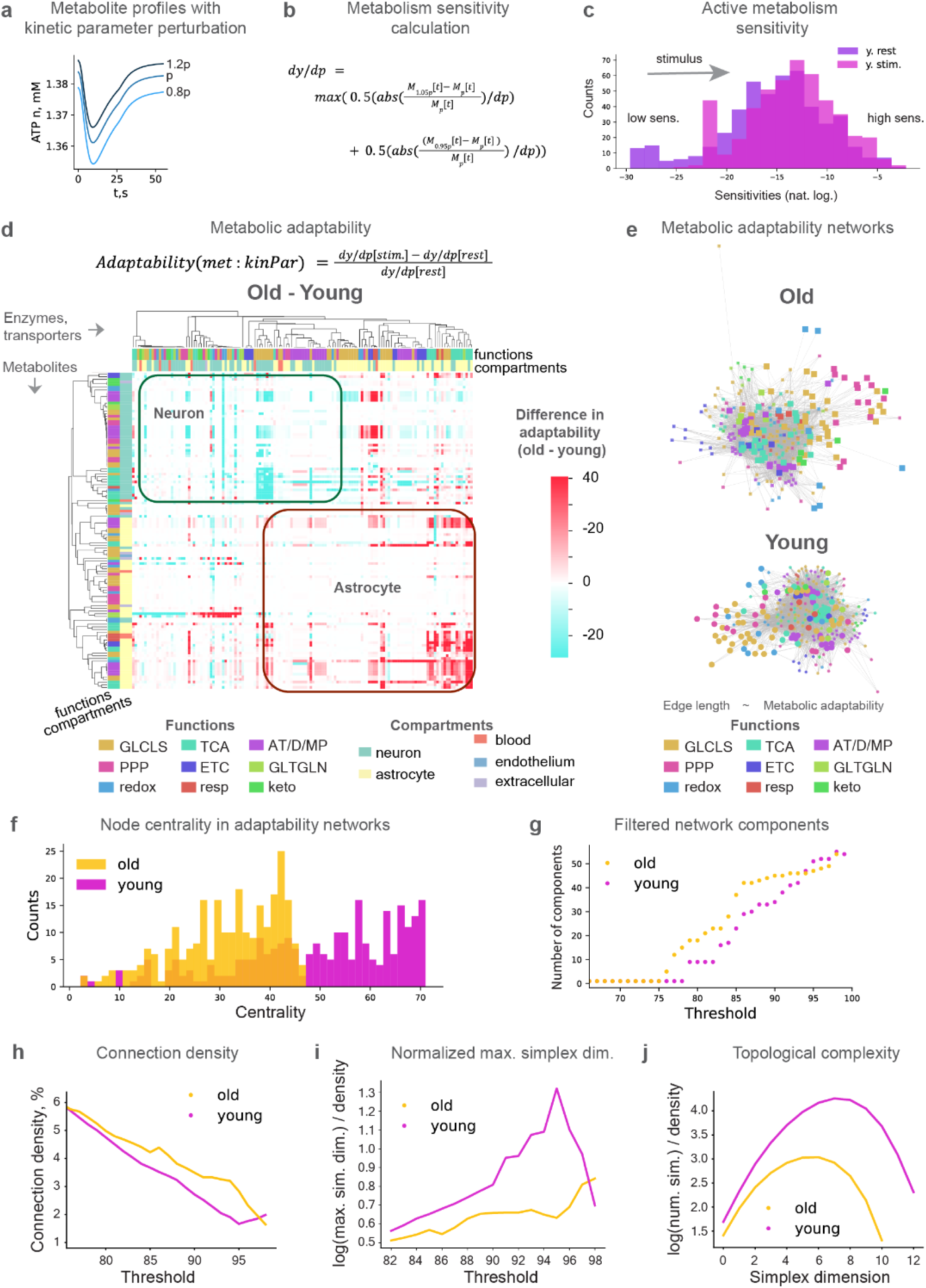
Metabolic response to kinetic perturbations changes with age. **a,** Example metabolite level profiles in response to kinetic parameter perturbation. **b,** Calculation of metabolic sensitivity to kinetic parameter perturbations. **c,** Active metabolism sensitivity. **d,** Metabolic adaptability to kinetic parameter perturbations. **e,** Metabolic adaptability networks in young and aged (same function-color relation as in **d**). **f,** Centrality of the nodes in the networks of metabolic adaptability aggregated by enzymes. **g,** Number of connected components in filtered networks of metabolic adaptability aggregated by enzymes. Ions, membrane potential, gating variables, mitochondrial membrane potential, and metabolites with fixed concentrations are removed from the analysis for all figures in this panel. **h,** Connection density of filtered networks. **i,** Maximum simplex dimension (log-transformed) normalized by connection density. **j,** Number of simplices (log-transformed) normalized by connection density (at 88% filtering threshold).

This metric allowed us to compare how the whole metabolic system of neurons and in astrocytes in the young and aged brain (Fig. 4d). Most of the metabolites in neurons decrease their adaptability with age, while adaptability of the astrocyte mostly increases. This observation is in line with the literature on astrocyte reactivity, which measures a set of phenotypic characteristics, including those of metabolism, inflammatory cytokines secretion and cytoskeleton rearrangements (Weber and Barros, 2015). However, in contrast to the “selfish” astrocyte hypothesis (Weber and Barros, 2015), we suggest that the increase in astrocytic adaptability is rather a “self-sacrifice” in an attempt to support the declining neurons.

We visualized the adaptability of the entire NGV metabolic network in the two age groups by positioning the nodes of both metabolites and enzymes using the Fruchterman-Reingold force-directed algorithm (Hagberg et al., 2008). The length of each of the 16800 edges were weighted by the inverse of metabolic adaptability (Fig. 4e; see Supplementary Information) to more intuitively reflect “metabolic fragility”. These networks displayed clustering of nodes largely by function and also revealed more evenly distributed clusters in young than in old systems, indicative of a robust network. To quantify the network differences between young and aged systems, we calculated the centrality of nodes, which is the reciprocal of the sum of shortest path distances between each node and all other nodes. The aged network showed on average longer distances than the young network (Fig. 4f), suggesting that the aged brain’s metabolic system is more fragile than in the young brain.

To quantify metabolic fragility in the network of the aged metabolic system we progressively removed edges below a given percentile and calculated the number of connected components in young and old networks (Fig. 4g). This analysis revealed that the aged network is fragmented into clusters or “islands”. Both networks are fully connected at thresholds below 76% and fully disconnected at 100%, but between 76% and 93% thresholds, we observed a higher number of connected islands in the aged network. We computed directed simplices, a type of all-to-all connected clique, using algebraic topology (Reimann et al., 2017; Sizemore et al., 2018)), to quantify the topological complexity of the network. This analysis revealed that the dimensions (number of nodes) of simplices and the number of simplices (see Methods) are higher in the young state (Fig. 4h-j), indicating that the young metabolic network is more topological complex, more distributed and more robust than in the aged system.

### Potential drug targets to repair the aging metabolic system

The scale of the challenge of finding new drugs for therapeutic interventions is revealed by the more than 16800 possible enzyme/transporter-metabolite interaction pathways that we identified in the NGV’s metabolic network, in addition to the complexity of the metabolic response when any one pathway is perturbed. The measure of metabolic adaptability can guide identification of targets in the context of this complex dynamical system. Here, interaction pathways with the highest differences in metabolic adaptability (Supplementary Fig. 16) are potential targets to repair the aged metabolic system (Fig. 5a) with high priority targets being those that improve adaptability for the highest number of pathways. The ideal drug to repair the metabolic system is one that acts like transcription factors (TFs), regulating multiple enzymes and transporters to modulate an even larger number of metabolic pathways. We therefore applied the ChEA3 optimization algorithm (Keenan et al., 2019), which isolates the TFs with the largest overlap between a prioritized set of genes for those enzymes and transporters that show the biggest improvement in metabolic adaptability for the largest number of interaction pathways (Fig. 5b). We identified ten highest priority potential targets.

**Fig. 5.**
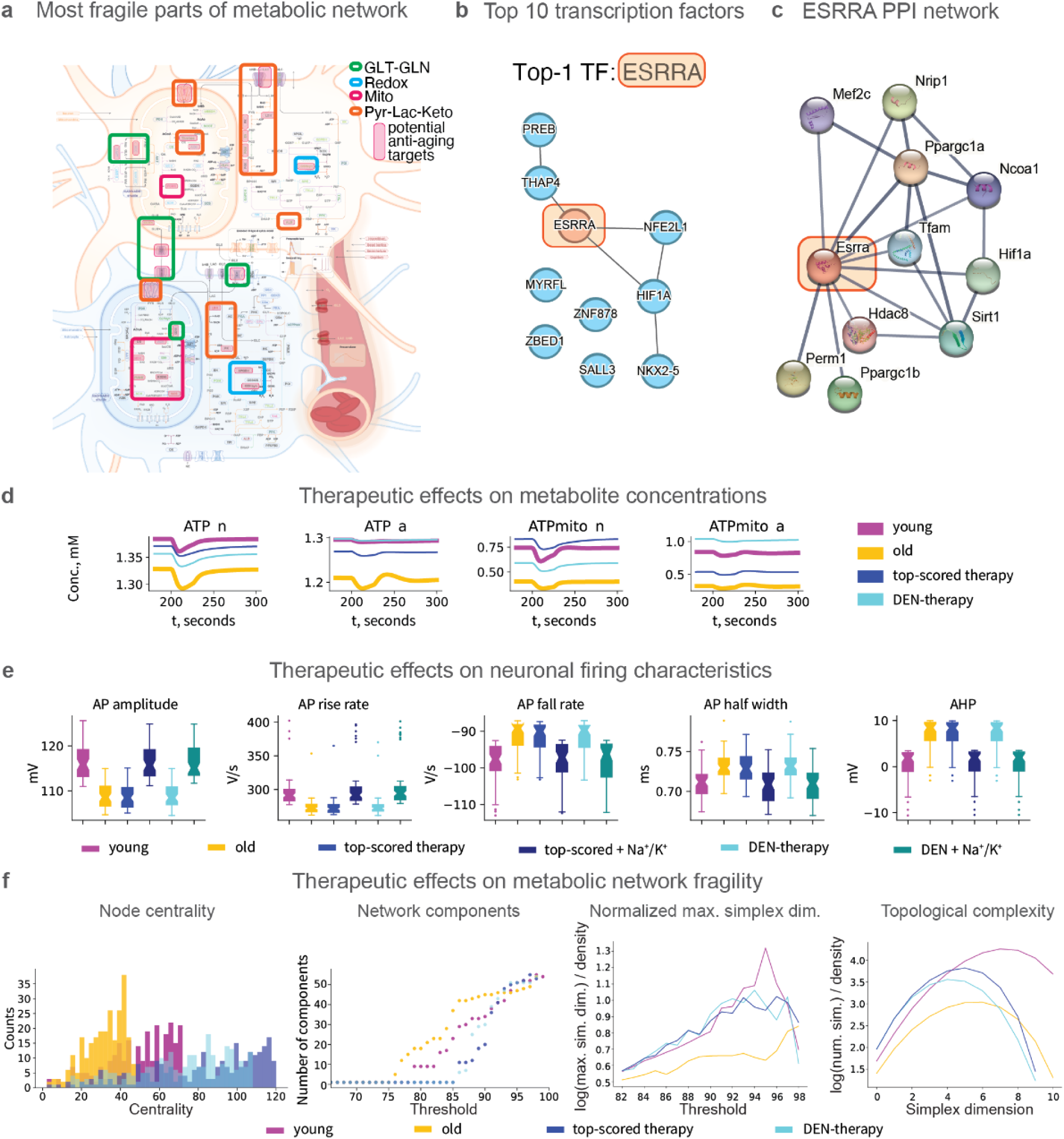
Reversing aging via targeted metabolism interventions. **a,** Sensitivity analysis-based potential targets are outlined by pink boxes and grouped by function in thick line boxes in the modeled system. **b,** Transcription factor enrichment results obtained from ChEA3 analysis (the top 10 TFs are shown). **c,** Results of the STRING-database search for ESRRA (the top TF from ChEA3 analysis). **d,** Time series traces of selected variables in young, aged, and treated aged states. **e,** Characteristics of neuronal firing in young, aged, and treated aged with selected therapies. In addition to selected top-performing and top-translatable therapies, we restored Na^+^/K^+^ pump expression to the young state. Application of the Na^+^/K^+^ pump expression restoration and each of the treatments restored characteristics of neuronal firing. Center line represents the median. **f,** Therapeutic effects on metabolic network fragility.

The TF with the best score was ESRRA (estrogen-related receptor alpha). This TF regulates expression of multiple metabolism-related genes, including those of mitochondrial function, biogenesis and turnover, as well as lipid catabolism (Tripathi et al., 2020). It is also linked to autophagy and NF-kB inflammatory response via Sirt1 signaling (Cantó et al., 2009; Yuk et al., 2015; Kim et al., 2018; Suresh et al., 2018). Mitochondrial dysfunction and autophagy impairments are consistently among the hallmarks of aging (López-Otín et al., 2013, 2023; Mattson and Arumugam, 2018; Amorim et al., 2022). Notably, ESRRA expression is downregulated in aging according to various studies (Schaum et al., 2020; Tripathi et al., 2020). Altogether, ESRRA acts as a regulatory hub of multiple aging-associated pathways as outlined in Supplementary Fig. 19. The other transcription factors that we identified are also validated by literature reports on TFs implicated in aging and neurodegeneration (see Supplementary Information).

To identify the major proteins associated with the top-scoring ESRRA (Fig. 5c), we searched the STRING database (Szklarczyk et al., 2019) and found Hif1a, Sirt1, Hdac8, Ppargc1a, Ppargc1b, Mef2c, Nrip1, Ncoa1, Tfam, Perm1 as the most prominent proteins involved. Numerous literature reports implicate these proteins in aging and neurodegeneration. The repair targets identified using our molecular model of the NGV system therefore largely aligns with experimental data in the literature on therapeutics for healthy aging (Campisi et al., 2019). We additionally suggest a role for less studied TFs in aging brain energy metabolism and provide insights into the links between molecular mechanisms implicated in aging and neurodegeneration (see Supplementary Information).

### Potential strategic interventions to repair the aging metabolic system

As an alternative to specifically targeting the enzymes and transporters, we investigated whether key features of the aged brain phenotype, such as energy deficiency and altered neuronal firing, could be repaired through strategic interventions. We conducted constrained optimizations (see Methods) for 1) the interaction pathway targets identified by the differences in metabolic adaptability (same as the input for transcription factor enrichment analysis above), 2) the interaction pathways potentially regulated by top transcription factor from the enrichment results mentioned earlier (ESRRA), 3) parameters corresponding to arterial blood glucose and ketone levels (mimicking dietary factors), 4) parameters corresponding to arterial blood lactate levels (mimicking exercise factors), and 5) total NAD-pool parameter in neuron and astrocyte (NAD-related supplementation). Surprisingly, optimization using a combination of diet (lower blood glucose and higher blood beta-hydroxybutyrate), exercise (higher blood lactate), and NAD-related supplementation and modulation of the cytosol-mitochondria NAD-associated reducing equivalents shuttle (hereafter referred to as DEN therapy), resulted in increase ATP levels in both neurons and astrocytes towards values of the young metabolic system comparable to that of the top-scoring drug targets (Fig. 5d, Supplementary Table 3). Interestingly, even though the parameter bounds for the optimisation were allowed to search for increasing or decreasing values, the DEN-therapy optimization converged unguided to a lower blood glucose, higher blood beta-hydroxybutyrate, higher blood lactate, and NAD-modulation, which are consistent with commonly accepted benefits of calorie restriction, exercise and NAD supplementation.

The DEN therapy restored the aged metabolic system, but not the action potential generation of neurons. As previously presented, restoring action potential amplitude and shape in our model can only be achieved by increasing the levels of the Na+/K+ pump back to youthful levels. We therefore additionally reversed the Na+/K+ pump’s age-related downregulation for each intervention (top-scored and DEN-therapy). This approach indeed, restored neuronal firing characteristics similar to those of a young state for each intervention (Fig. 5e) as well as ATP levels of both neurons and astrocytes. Interestingly, insulin is a common factor that activates the Na^+^/K^+^-ATPase, increases its expression, and also lowers blood glucose, consistent with DEN-therapy. When we performed a sensitivity analysis and calculated adaptabilities for the DEN-therapy and top-scored therapy we found that not only could network fragility be repaired, but even improved more than the young state (Fig. 5f).

## Discussion

This study presents a molecular model of the neuro-glial vascular system that brings together the key cellular and subcellular systems, molecules, metabolic pathways and processes required to couple neuronal electrical behavior with brain energy metabolism and blood flow. The data-driven strategy developed allows applying experimental data, in principle from any condition, to produce a model of that condition. We applied experimental data from the young and aged brain metabolic systems to produce a model for their respective metabolic systems. We identified 16’800 enzyme/transporter - metabolite interaction pathways in the brain’s metabolic system. A sensitivity analysis for each pathway produced a comprehensive view of how each pathway impacts each other pathway to support action potential generation. We find that the impact of one pathway on all the others is remarkably evenly distributed, indicative of a highly robust system with multiple routes to respond to changing metabolic demands and a system that is resilient to damage of any one pathway. By normalizing to resting sensitivities of each pathway we could develop a systems measure for metabolic adaptability for each pathway to evaluate changes in a different condition such as in the aged brain. Our analysis suggests that the aged metabolic system breaks down into islands where enzyme/transporter-metabolite interaction pathways cluster more than in the young brain, leaving this complex molecular system less robust to damage and more restricted when responding to stimuli. We identified the transcription factor, ESRRA and several key proteins it regulates, as top potential drug targets and a prioritization of potential strategic interventions that can repair the aging metabolic system.

The data-driven model we developed captures how brain energy metabolism interacts with neuronal activity with a high degree of biological fidelity. Each enzyme and transporter is modeled using an experiment-derived rate equation featuring its concentration, key kinetic properties, and effects of inhibitors and activators (where applicable and relevant). This approach allows integration of proteomics and transcriptomics data for modeling various conditions and diseases in which molecular levels and properties are affected. Compared to the more generalized phenomenological metabolic models, the model features 183 processes, including 95 enzymatic reactions, 19 processes for the transport of molecules across the cell and mitochondrial membranes, and 69 other processes for ionic currents, blood flow dynamics and other related non-enzymatic processes. Changing molecular concentrations are simulated using 151 differential equations. Additionally, cytosolic ADP, creatine, NAD, NADP are computed from the conservation law and total pool of relevant molecules.

The strategy we followed to build such a complex model was to apply biologically reported parameters for the components of the model (see Methods) and avoid overriding biological values to fit reports on time series of metabolic responses found in the literature that are few and often contradictory. In order for the system of equations to have a solution, we optimized the parameters by only requiring steady state solutions at rest, rather than changing the parameters to fit the metabolic time series responses reported in the literature, which are limited and often contradictory. Alternative approaches that have been used by others include likelihood-based optimisation targeting the reference time series data. This approach was not suitable in our case because the data was not available for most metabolites for meaningful likelihood-based parameter estimation with recorded traces of metabolite levels in the neurons and astrocytes. Bayesian parameter estimation, also previously used, was computationally too costly for the scale and complexity of our model. To increase the biological dataset for parameterization, we merged data across *in vitro* and *in vivo* conditions and averaged across natural biological variability. In some cases, we had to optimize weakly constrained parameters or include only the most relevant components, pathways and processes (see Methods). The model, while containing an unprecedented level of detail, is also not yet at a whole genome level. Similarly, while the model captures the key cellular elements, compartments and sub-compartments it does not yet capture explicitly the details on all possible geometric constraints.

The model validated against numerous experimental datasets, but a key litmus test was simply whether computational convergence occurred for this complex system. Parameters were minimally optimized to allow convergence for a steady state at rest, but a self-constrained converged state emerged when the system was stimulated with current injection and synaptic input. On the other hand, when we introduced random modifications to enzyme and transporter concentrations and their kinetic parameters, some numeric solutions failed or diverged far from the steady state at rest. It is therefore even more remarkable that simulations converged without significant modifications introduced when we imported and applied the data from the aged brain. Furthermore, single parameter perturbations introduced instabilities in the simulations more often than when multiple RNA-seq derived changes were introduced, suggesting that the set of parameters are self-constraining.

Our results in both young and aged brain states align well with a wide range of published experimental reports. Aside from the time series profiles of specific metabolites, enzymatic activities and, aging-observations, it is particularly noteworthy that the estimates that emerged from the simulations for the ATP consumption (Howarth et al., 2012; Yi and Grill, 2019; Zhu et al., 2019) and how aging-associated changes in metabolism affect neuronal action potentials are consistent with experimental reports (Power et al., 2002; Disterhoft and Oh, 2007; Kumar and Foster, 2007; Smithers et al., 2017; Vitale et al., 2021).

Calculating sensitivities is common when studying dynamical systems. In addition to sensitivity analysis, for the system undergoing transition between rest and stimulated states, we introduced adaptability and fragility as biologically interpretable measures. These measures capture the effects of perturbing an enzyme or transporter on all the metabolite levels in response to stimuli. These perturbations mimic the effects of conditions such as phosphorylation levels, transcription and translation errors, as well as molecular damage to enzyme and transporter kinetic properties. Perturbation analysis predicted diminished adaptability to changing energy demands with different changes in neurons and astrocytes in the aged brain. We could construct a network of enzyme-transporter - metabolite interaction pathways where each pathway can be evaluated in terms of metabolic adaptability allowing quantification of the changes. We find a structural breakdown and decreased topological complexity of the NGVmetabolic systems in the aged network as compared to young.

To identify potential targets where the youthful state can be restored, we identified the most fragile interaction pathways. We performed transcription factor enrichment analyses for the most sensitive enzymes and transporters. Functions of these targets largely overlap with known mechanisms of aging. Through constrained optimization, we identified a combination therapy that restores key features of the aging brain phenotype. This therapy involves maintaining specific levels of blood glucose, lactate, and β-hydroxybutyrate achievable through diet and exercise, coupled with redox state maintenance via NAD-supplementation, modulation of the cytosol-mitochondria reducing equivalent shuttle (related to NADH), and Na^+^/K^+^-ATPase activation. For instance, reversing the aging phenotype can be achieved in part by regulating insulin signaling, which lowers blood glucose and activates Na^+^/K^+^-ATPase.

Complex interventions that act on multiple enzymatic targets at the same time, including some of the top potential targets of DEN-therapy, also managed to restore ATP levels in cells. However, their development and implementation would require significantly more extensive research before they could be considered for practical application in treating aging-related conditions. The outcomes of these complex therapies appear to be comparable to those achieved with the simpler DEN-therapy.

The promising combination therapy identified in this study, which includes diet, exercise, NAD supplementation, NAD shuttle and Na^+^/K^+^-ATPase modulation, agrees well with proposed anti-aging interventions such as caloric restriction, the ketogenic diet, and exercise (López-Otín et al., 2023). Physical exercise shows beneficial anti-aging and brain-health effects mediated by the brain-derived neurotrophic factor (BDNF), insulin-like growth factor 1 (IGF-1) and lactate (Horowitz et al., 2020; Stillman et al., 2020; Xue et al., 2022). The ketogenic diet and caloric restriction, for example, impact the levels of β-hydroxybutyrate and glucose in the blood (Meidenbauer et al., 2014). Supplements such as urolithin (Singh et al., 2022), metformin (Kulkarni et al., 2020), and nicotinamide mononucleotide (Yoshino et al., 2018) affect mitochondrial health, energy supply and provide anti-inflammatory action, consistent with the important role of metabolism in aging.

The multiple validations and insights consistent with current findings suggest that the model can guide experiments on brain aging and diseases, including those on disease-associated genetic variants, enzymatic deficiencies, and the effects of different practical strategies. Energy-metabolism related transcriptomics, proteomics and metabolomics data can also be applied to the model to study their effects on metabolic dynamics and neuronal firing. Furthermore, the model can simulate a variety of stimuli to neurons to guide studies on the energy constraints of brain activity. To accelerate these research areas, the model is open sourced for public use (link provided after acceptance).

## Data Availability

All the data used in this study are publicly available from referenced sources. The model is available on the web portal (https://bbp.epfl.ch/ngv-portal-dev/) [it will be available after peer-reviewed publication].

## Funding

This study was supported by funding to the Blue Brain Project, a research center of the École Polytechnique Fédérale de Lausanne, from the Swiss government’s ETH Board of the Swiss Federal Institutes of Technology.

## Acknowledgements

The authors thank Judit Planas Carbonell, Claudia Savoia, and Jean-Denis Courcol for organizing web portal development and visualization, and Matthias Wolf for software support. We thank Karin Holm for writing assistance and Ayima Okeeva for the model notebook evaluation.

## Author contributions

PS: developed the model, performed analyses, developed the figures, contributed to conceptualization and interpretation of results, metabolic network metrics and writing. JSC: contributed to the conceptualization, interpretation of results and writing. LK: contributed the topological analysis and interpretation of network results, and related figures. EB: contributed to figures. CF: developed the movie on the portal. SA: developed the web portal. DK: contributed to conceptualization, interpretation of results, and writing. HM: contributed to conceptualization, interpretation of results, metabolic network metrics, and writing.

## Declaration of interests

The authors declare no competing interests.

## Methods

### Baseline model building

We reconstructed and simulated a model of NGV metabolism coupled to a simple blood flow model and a Hodgkin-Huxley (HH) type of neuron model. The main concepts of electro-metabo-vascular coupling, as well as blood flow and neuronal electrophysiology model are based on the models available from the literature (Aubert et al., 2001; Jolivet et al., 2015; Calvetti et al., 2018; Winter et al., 2018). Our model specifically emphasizes the key brain energy metabolism pathways and processes involved in neuronal signal transduction. However, to gain a more comprehensive understanding of the various complementary molecular mechanisms and pathways involved in aging and disease, it is desirable to further expand the model to a whole-cell scale and incorporate more regulatory processes. At present, this task is hindered by data sparsity. As more data becomes available, the model can be iteratively refined and expanded.

Compared to the more generalized phenomenological metabolism models, our metabolism model features 183 processes (95 enzymatic reactions, 19 processes of transport of molecules across the cell and mitochondrial membranes, and 69 other processes related to ionic currents, blood flow dynamics and some miscellaneous non-enzymatic processes, e.g. Mg^2+^ binding to mitochondrial adenine nucleotides). Every reaction, transport or other process is represented by its rate equation, which is literature-derived. Changes of molecular concentrations are described by a system of 151 differential equations and additionally cytosolic ADP, creatine, NAD, NADP are calculated from the conservation law and total pool of relevant molecules.

The model is based on literature data for enzyme kinetics and molecular concentrations. We have meticulously collected all parameters and equations from literature sources, as referenced in Supplementary Table 2 and throughout the model code, and programmatically queried databases BRENDA (Chang et al., 2021), SabioRK (Wittig et al., 2018). However, observed discrepancies in the parameters reported by different sources define the need for an optimization procedure, to derive plausible biological middle-ground. The parameters with uncertainties observed in the literature were constrained by their lower and upper bounds taking into account the type of the parameter (Michaelis constant of reaction, inhibition/activation constant, maximal rate of reaction, equilibrium constant, Hill coefficient) and optimized as described in the Optimization part of the Methods.

To have the most realistic biological average for the initial values of all variables (concentrations, membrane potential, mitochondria membrane potential, venous volume, gating variables) according to the literature, we considered not only measured and modeled literature data on the absolute values themselves, but also additional constraints, such as known ratios of NADH to NAD^+^ in the neuron (Neves, 2011; Dienel, 2012; Berndt et al., 2015; Mongeon et al., 2016) and astrocyte (Mongeon et al., 2016). One of the most important variables in the model, ATP concentration, was reported as being 2 mM in many experimental and modeling studies (Erecińska and Silver, 1989; Cloutier et al., 2009; Jolivet et al., 2015; Calvetti et al., 2018; Winter et al., 2018). However, some more recent data report it at 1 to 1.5 mM scale (Baeza-Lehnert et al., 2019; Köhler et al., 2020). Assuming that more recent measurement technologies can provide more precise data, we set cytosolic ATP in the neuron to approximately 1.4 mM according to Baeza-Lehnert et al. (Baeza-Lehnert et al., 2019) and to approximately 1.3 mM in the astrocyte according to Kohler et al. (Köhler et al., 2020) where it was reported in a range of 0.7 to 1.3 mM (acutely isolated cortical slices) and 1.5 mM (primary cultures of cortical astrocytes).

Mammalian ATP to ADP ratios are reported in a very wide range of values from 1 to more than 100 (Tantama et al., 2013). Ratio of ATP to AMP is around 100 (Erecińska and Silver, 1989). Furthermore, metabolite ratios from (Erecińska and Silver, 1989) were used to adjust initial concentrations of phosphocreatine and phosphate to the ATP levels. Lactate concentrations in different compartments, which is central to the ANLS debate, was set according to the recent Mächler et al. paper (Mächler et al., 2016). We also tested the model with all alternative literature concentrations for the metabolites mentioned above.

Glucose supply from blood is of key importance to brain energy metabolism (Benton et al., 1996). For this reason we approached it particularly meticulously. In our model, glucose concentrations are assigned to detailed compartments, such as arterial, capillary, endothelial, basal lamina, interstitium, neuronal cytosol and astrocytic cytosol (Barros et al., 2017). According to the literature, hexokinase flux is split approximately equally between neuron and astrocyte (Barros et al., 2007, 2017; Jolivet et al., 2010), so that we adjusted Vmax of hexokinase so that its flux matched the literature data at rest.

Upon activation, the ratio of glucose influx to astrocyte versus neuron increases, consistently with the literature knowledge (Jolivet et al., 2010).

### Implementation and simulation

This metabolism model is implemented and simulated in Julia programming language (Bezanson et al., 2017). We used the DifferentialEquations.jl package (Rackauckas and Nie, 2017) to solve the differential equations system using order 2/3 L-stable Rosenbrock-W method (autodifferentiation disabled, both absolute and relative tolerances set to 1e-8). We chose to use the Julia language because of its high performance, its extensively developed mathematical methods ecosystem, and the readability of the code, which supports its future use. Most of the analysis and figures-making code is written in Python programming language.

The model is built in a modular way, so that every molecular process has a dedicated rate function, and combination of relevant rate functions defines the dynamics of variables. This supports convenient testing of various enzymatic mechanisms, parameters and initial values of concentrations, as well as easier model subsetting and expansion.

Once the manuscript is accepted for publication, we will provide the GitHub repository with the code for model simulation, optimization, validation and analysis. These scripts are aimed to facilitate the model’s reuse in future studies.

### Optimization

Time series data on the dynamics of specific metabolites in neurons and astrocytes is very sparse and sometimes contradictory. To avoid favoring one data source over another, we only performed optimizations for the steady state (minimizing derivatives). We built and optimized the model bottom-up in multiple iterations, gradually expanding it with more details. We started with the model of neuronal electrophysiology (Pospischil et al., 2008; Øyehaug et al., 2012; Jolivet et al., 2015; Krishnan et al., 2015; Calvetti et al., 2018). We included detailed astrocytic ion management based on the existing literature model (Witthoft et al., 2013). Then for the metabolism model, we started with capillary dynamics, oxygen and glucose transport, and hexokinase, because they are very well studied and CMR of glucose is widely measured, which sets a strong constraint on hexokinase rate. Then we proceeded to add reaction by reaction and evaluate rates in simulations, each time adding a new reaction, first if needed roughly manually refining underconstrained parameters. Then after several reactions were added, we ran optimization (with an objective to minimize derivatives) for a selected small set of parameters which are the least constraint by the literature. Then we modeled lactate transport and connected it to glycolysis. We separately optimized PPP for steady state (with an objective to minimize derivatives). For the mitochondria, we started from the electron transport chain, which is mitochondrial-membrane potential dependent and extremely sensitive to parameter variations. We mostly used the ETC model from Theurey and the colleagues (Theurey et al., 2019), and then we carefully selected a small number of parameters to optimize them (with an objective to minimize derivatives) to make the ETC model compatible with ATP and ADP concentrations from more recent experimental evidence. Then we added one-by-one TCA reactions to ETC, the same way as described above for other pathways. And we also added ketones metabolism, part of MAS, glutamate-glutamine cycle (after having both neuron and astrocyte together in the system).

The optimization procedure referenced above is single objective optimization performed using BlackBoxOptim.jl [https://github.com/robertfeldt/BlackBoxOptim.jl of Robert Feldt] with the default algorithm (adaptive differential evolution optimizer) iteratively selecting different sets of processes to reduce the parameter space.

To avoid non-physiological molecular concentrations (negative or too high values), we used Julia-callbacks and the “isoutofdomain” mechanism in solving the differential equations system during optimization. For these biological plausibility reasons, we utilized “isoutofdomain” to control the solution of the differential equations system to stay non-negative, so that the solver takes smaller time steps if the solution leaves the domain, unless the minimum step size is reached and integration is terminated. The same methods were applied for the anti-aging optimization, but the selection of neuronal firing related variables from the young state simulated time series data were used for the objective function.

Computational models are often optimized by fitting parameters to the data using a selected algorithm. Indeed, some time series data are available for various aspects of brain metabolism, including for concentrations of glucose, lactate, pyruvate, NADH and ATP, the BOLD signal, and cerebral metabolic rates of oxygen and glucose. However, to our knowledge these usually come from different experiments rather than simultaneous measurements of multiple metabolite concentrations and other characteristics. It has been demonstrated by numerous studies that one can fit system dynamics to selected data given a sufficient number of weakly constrained parameters and nonlinear rate equations (Dyson, 2004). An interesting case is when measurements with similar metadata from different studies produce significantly different dynamics of metabolite concentrations, such as in the example of extracellular brain glucose from Kiyatkin and Lenoir (Kiyatkin and Lenoir, 2012) as compared to Fillenz and Lowry (Fillenz and Lowry, 1998), which was further used in one of the early integrative NGV models (Cloutier et al., 2009). We therefore aimed to avoid the global optimization of fitting parameters to selected time series. Instead we iteratively refined the bottom-up model by estimating parameters that would achieve the desired values of metabolite concentrations at steady state (in which the concentration derivatives with respect to time are minimized). More details are available in the next section (workflow and the key aspects) and the entire pipeline is shown in Supplementary Fig. 17. However, this approach has a downside: it does not guarantee exact matching of the experimentally recorded dynamics of any selected experiment. Good match with the time series observed experimentally and in other models can only be obtained if the underlying model has a sufficient level of detail, uses relevant kinetic data for initial parameterization and employs applicable constraints (e.g., physiological range of metabolite concentrations, typical range of values for kinetic parameters of a given type). While many time series produced by our model are close to the literature reports, glucose concentration traces and cerebral metabolic rate of glucose consumption have only modest stimulus responses as compared to the literature. This can be explained by our decision to follow the most detailed (to our knowledge) approach to glucose transport in the brain available in the literature (Barros et al., 2007; Simpson et al., 2007). This approach takes into account compartmentalisation into arterial, capillary, basal lamina, interstitial space, astrocytes and neurons, with glucose transfer between these compartments described by rates that consider intracellular/extracellular concentration-dependent trans-acceleration and asymmetry of transporters.

### Workflow and the key aspects of the bottom-up model building and optimization

In order to build the model in a bottom-up data-driven way and avoid unreasonable preference for any particular data source, we developed a workflow, which resulted in the model performing surprisingly well for different setups. It produced quality simulation outcomes which are largely consistent with various literature. The only drawbacks are that the workflow is largely iterative, time-demanding, and requires manual intervention. Here are the steps and the key considerations.

**Step 1.** Collect as much reliable data as available. In our case of building a model which combines the metabolism, electrophysiology and the blood flow, the following data were needed: molar concentrations of molecules (metabolites, proteins, ions), enzymes and transporters kinetic parameters, electrophysiology and blood flow dynamics parameters, rate equations of all processes, mechanisms of reactions and the data on their inhibitors and activators with corresponding mechanisms of action, existing models of pathways and their combinations. In most cases, reaction rate is modeled in the literature with at least a few different equations. This is due to the use of different formalisms. For example, the same reaction can be described in a precise mechanistic way considering multiple transition states of complexes formed by enzyme with substrates, products, regulators, or it can be described in a more simplified form of modular rate law or Michaelis-Menten kinetics, when assumptions about the reaction mechanism are met. It is important to keep collected models of reactions and how they are used in the existing models of pathways and systems, because for practical applications the scale of the model needs to be balanced with how many parameters are used for each equation in the model. For example, detailed mechanistic rate equations can be parameterized well for small models when there is enough consistent reliable data, but for cases with high uncertainty in the data, it is often hard to optimize and not overfit such models.

**Step 2.** Next, we model individual reactions. In some cases (most of which are relatively old biochemistry studies), time series data on individual enzymes are available. These can be used to optimize the parameters of enzymatic rate equations, especially if they are underconstrained, coming from different species or tissues. This step also allows us to evaluate how fast individual reactions are, how significant are the effects of inhibitors and activators and whether to include them in the model or not, and how problematic each particular reaction is in terms of the steady state and response to changing inputs.

**Step 3.** Once the data is collected, we bring together reactions one-at-a-time according to the reconstructed pathways networks. This process is highly iterative and needs to be repeated multiple times starting from different data. We need to try multiple combinations to see in which cases the optimization needed to bring the combination close to steady state is minimal. It is also important to combine those small subsets of reactions with pseudo-reactions of substrates source flux and products sink flux, to have an estimate of how this unit will perform once it is plugged into a bigger system. Iterating on this step, one can grow the system up to the models of pathways in individual cells, and existing models of those pathways are very helpful for initial choice of the most promising combinations of reaction rates and parameters. It is also useful to keep approximately the same level of detail for the equations of all reactions in the pathway. For the refinement of the parameters when connecting reactions in a pathway, instead of just following commonly used list of reactions of the pathway in order the metabolites enter it, it is useful to start from different steps of the pathway, especially with the reactions which are either known key regulators of the overall pathway flux (bottlenecks) or close to connection points to other pathways, or those with the most complicated mechanisms. The key aspects to decide on the performance of selected parameters set in the model are the concentrations at the steady state (or pseudo-steady state if the formal one cannot be achieved in a reasonable time), their response to stimuli (at least qualitatively in which direction and approximately how fast do they change, when no data is available), reaction and transport fluxes. It is important to keep several best performing models for all subsystems/pathways, because once they are plugged into a bigger system, performance ranking can change.

**Step 4.** Once small units/pathways are built in at least a few variations, they can be connected into bigger systems. For the optimization of connecting reactions, it is important to start from different entry points, compare overall fluxes of the pathways, and consider volumetric scaling aspects. In some cases temporary use of pseudo-reactions for source and sink of some metabolites for the optimization significantly improves the performance.

**Step 5.** The large metabolic system can further be connected (using the same strategy as in Step 4) to the electrophysiology and blood flow models. Electrophysiology and blood flow models can be found in the literature in a number variations and need to be optimized separately if needed.

**Step 6.** The models of the neuron and the astrocyte can be connected in the same way as described above. Simulations and sensitivity analysis can further be used to select the parameters optimization of which has the highest effects and can efficiently improve the model according to available data. If no consistently reliable data is available, the objective function can be set to minimization of derivatives at rest state for the system to be at the steady state.

### Validation

First, we tested the response of the key metabolites (ATP, NADH, lactate, glucose) to the stimuli. All concentration related variables were ensured to stay in the range of biologically plausible values by the callbacks and the “isoutofdomain” parameter to a solver as described in the Optimization part of Methods. Next, we calculated the BOLD signal (Supplementary Figure 1d) and OGI (in range of 4.5-5 depending on stimulus, while literature data is in range of 4-5.5) using equations from Jolivet et al. 2015 to compare them with the literature (Jolivet et al., 2015; Winter et al., 2018; Jung et al., 2021). These two high-level phenomena are commonly used as benchmarks in NGV metabolism modeling papers (Jolivet et al., 2015; Calvetti et al., 2018; Winter et al., 2018). We also qualitatively compared dynamics of some key metabolites and reaction and transport fluxes to their expected response to stimuli. Then we estimated energy use from the components of the Na^+^/K^+^-ATPase rate equation (calculated from sum of neuron and astrocyte Na^+^/K^+^ pump ATP consumption flux in mM concentration per second with the volume of 17.8 um^3^ and the literature estimate of ionic gradients sharing 31% of total energy use) and compared it to the literature estimates (Howarth et al., 2012). We further validated aging-associated effects against the literature data shown in Supplementary Table 1.

### Implementing aging effects in the model

Aging is a multifactor phenomenon which affects metabolism at different levels: transcriptome, proteome, metabolome, and potentially even kinetic properties of enzymes and transporters due to accumulated genetic damage, lower protein synthesis fidelity and higher chances of protein misfolding. To implement the aging effect in our model in a fully data driven way, the data on neuron and astrocyte specific proteomics, metabolomics and kinetics of enzymes are needed. However, most of such data is not yet publicly available.

We modeled the aging effects as following:

1. enzymes and transporters expression fold changes from TMS dataset (Schaum et al., 2020; Zhang et al., 2021a) applied as scaling factors to levels of corresponding enzymes and transporters
2. scaled initial concentrations of blood glucose, lactate, beta-hydroxybutyrate according to the literature data on difference in their levels in aging (approximation, because effect size depends on the literature source)
3. total NAD+ and NADH concentration pool scaling, because it decreases in aging according to qualitative literature (approximation)
4. synaptic glutamate release pool (approximation, but synaptic input is set the same for comparability of the results)
5. scaling of reducing equivalents shuttles between cytosol and mitochondria: NADH shuttle is a generalized rate equation based on activity of multiple enzymes of malate-aspartate and glycerol-phosphate shuttles, for which we followed literature to model it (Jolivet et al., 2015).

For the above factors, which mention “approximative/approximation”, the direction of change is according to the literature, but the absolute number of scaling factors (not known/contradictory in the literature) is set with an objective for the model to be steady at rest.

We implemented the aging effects on enzyme and transporter levels in two parallel ways: 1) using cell-type specific transcriptomics data (Schaum et al., 2020; Zhang et al., 2021a) and 2) using integrated proteomics data from the metaanalysis we performed earlier (Shichkova et al., 2021). The first approach featured higher coverage depth for the astrocyte-specific data. So that, to reduce bias from inferring missing data in the second method, we decided to rely on RNA data for implementing aging effects into simulation, while we used the second data source as a part of validation.

### RNA fold changes for modeling aging effects

Anextensive single-cell transcriptomics mouse dataset has recently become available (Schaum et al., 2020; Zhang et al., 2021a), providing insights into the aging patterns of various cells including neurons and astrocytes. However, RNA needs to be translated into proteins. RNA data need to be used with caution when inferring age-dependent protein concentrations. Nonetheless, using RNA fold changes to scale enzyme and transporter levels results in metabolite concentration changes consistent with the literature (Supplementary Table 1).

We mapped reaction IDs to gene names using the gene-reaction-rules from a publicly available metabolism reconstruction Recon 3D (Brunk et al., 2018). Then for the cases of multiple genes per reaction (i.e. enzymes built of several protein subunits or different isoforms present at the same time) we calculated age-scaling in two ways: 1) by using geometric mean of all fold changes, and 2) taking fold changes which results in lowest levels of RNA in aging, i.e. using the assumption that each protein subunit or isoform can be rate limiting if it’s concentration is not sufficient to build fully-functional protein. We applied each of these methods twice: for all genes and only for those with significant changes (significance defined by the source data paper). Next, we manually went through the mapping of all genes to reactions and kept only those that are enzyme subunits/isoforms and not regulatory factors. We then refined it by subcellular location.

### Protein levels for modeling aging effects

Several studies measured brain protein levels in different ages, but they provided mostly brain tissue/regions data, rather than single neuron and astrocyte age-specific protein levels. The other studies provided neuron and astrocyte specific protein levels, but they were either using cultured cells, or young/adult rodents. For these reasons even a combination of proteomics data sets remains sparse in terms of cell-type and age specific protein quantification. Even though using protein levels directly to scale Vmax of the enzymes and transporters would allow consideration of posttranscriptional effects of protein synthesis and degradation, to reduce potential bias, we decided to rely only on the RNAseq data for age-associated changes in enzyme and transporter levels.

### Other necessary aging factors

Arterial glucose, lactate and b-hydroxybutyrate, as well as total NAD (reduced and oxidized) pool are fixed in the model, but multiple studies report that they change in aging. For this reason we scaled them according to the literature. The resulting model was far from steady state, which could be explained by some missing age-associated changes. We then scaled NADH exchange between mitochondria and cytosol, as it is also known to be affected by the aging process, and it resulted in a well-functioning model producing biologically meaningful observations. For a more realistic setup, we also scaled synaptic effects of glutamate concentration changes upon release events, but it had less effect and the age-associated changes in electric features extracted from simulations with only current injection are consistent with those driven synaptically.

### Adaptability calculation and search for potential anti-aging strategies

As described in the main text, the adaptability calculation is a modification of sensitivity analysis with perturbation of one parameter at a time by 20% of its initial value and subsequent calculation of the difference between the resting and stimulated state’s sensitivities, normalized by the resting state sensitivities (see formula in Fig. 4b). We then consider enzymes and transporters with the highest difference in adaptability between young and aged states as the most fragile and therefore potential anti-aging targets. Furthermore, to identify enriched transcription factors for these targets we applied the ChEA3 algorithm (Keenan et al., 2019). As described in the main text, we then performed constrained optimisation for 20 sets of parameters combining those of adaptability-based and the top transcription factor regulated enzymes and transporters, as well as parameters related to diet, exercise and NAD supplementation.

### Topological analysis

For the topological analysis of the adaptability networks, we employed methodology from algebraic topology. The distribution of directed simplices, introduced in Reimann et al. 2017 (Reimann et al., 2017), has been essential for the study of brain networks and has revealed significant links between the maximum simplex dimension and the robustness of networks. The distribution of directed simplices was computed with the open software https://github.com/JasonPSmith/flagser-count. Due to varied connectivity density, defined as the number of edges over the total number of possible edges, for different sensitivity thresholds, we divided the logarithm of the number of simplices by the connectivity density. This normalization allows us to compare networks of different connectivity densities and identify which parts of the networks are more susceptible to changes.

## Supplementary Information

**Supplementary Table 1.**
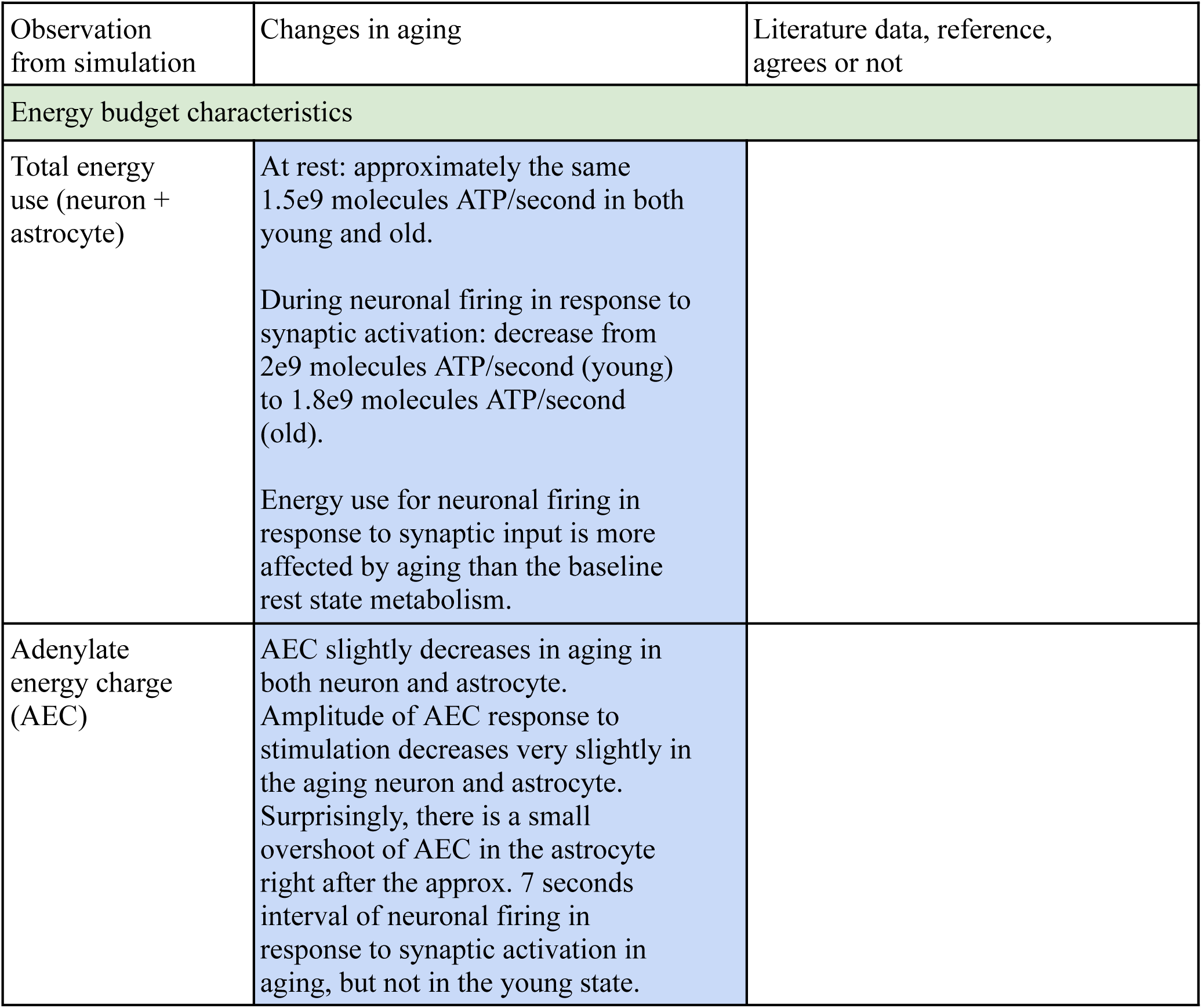
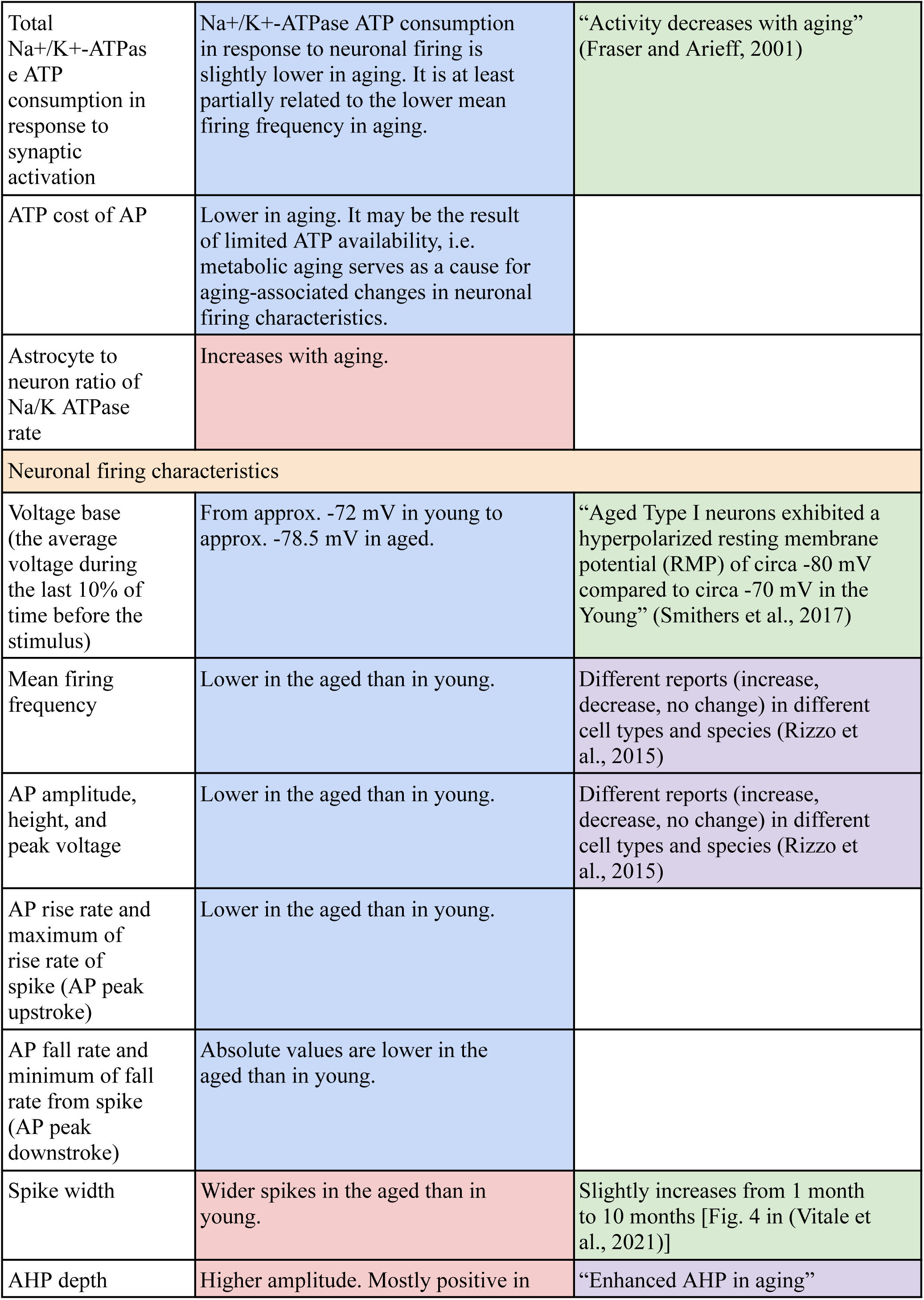
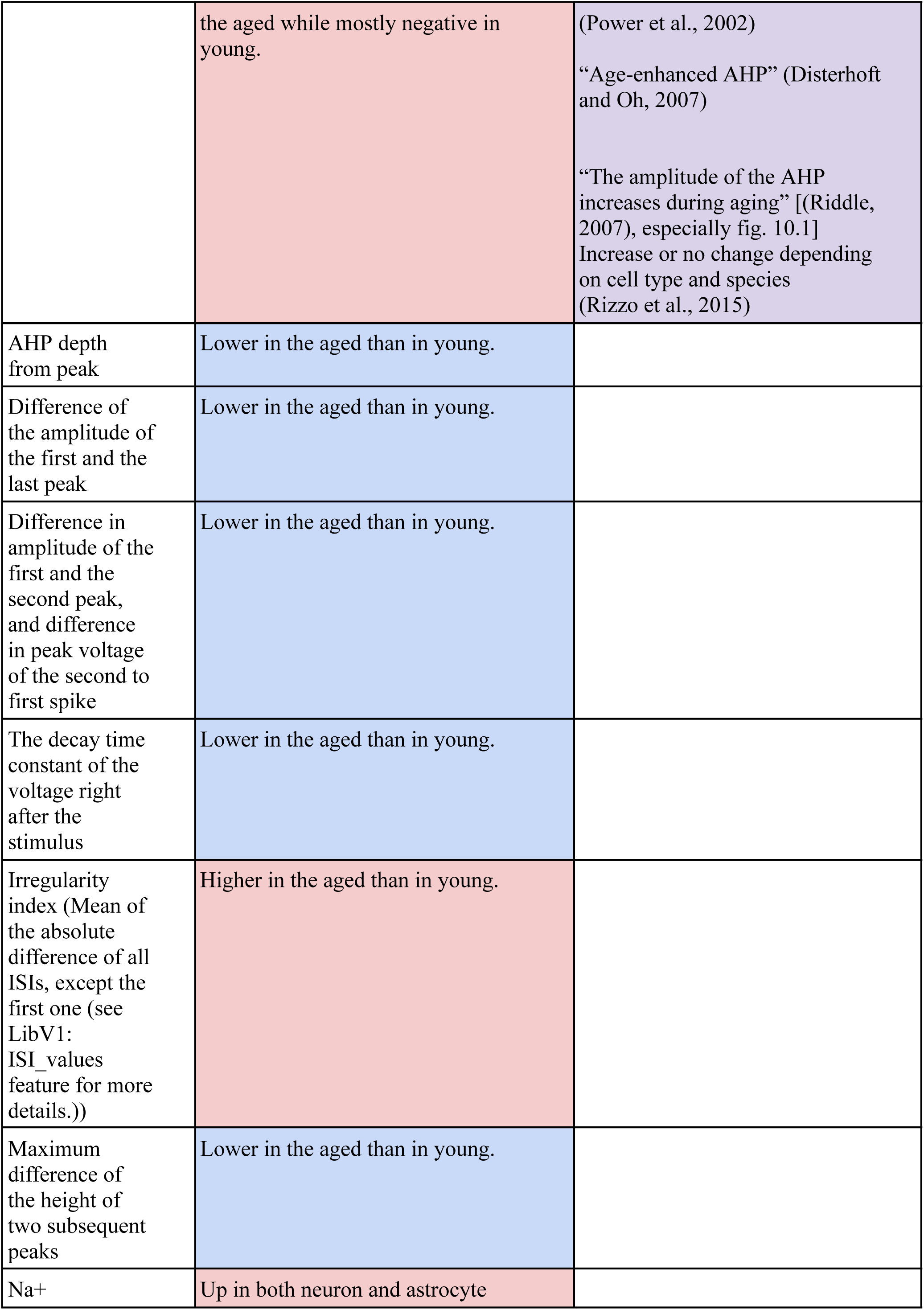
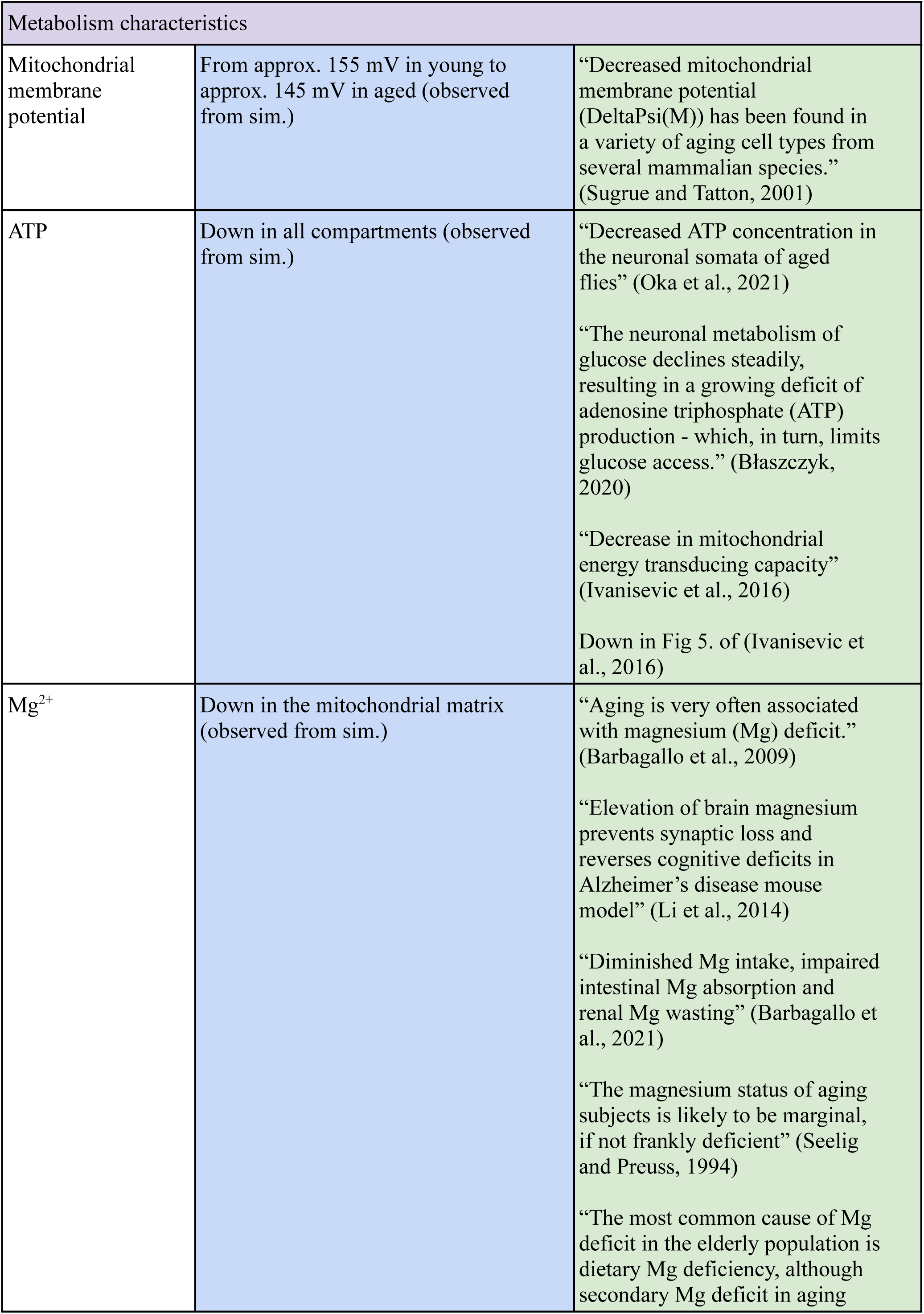
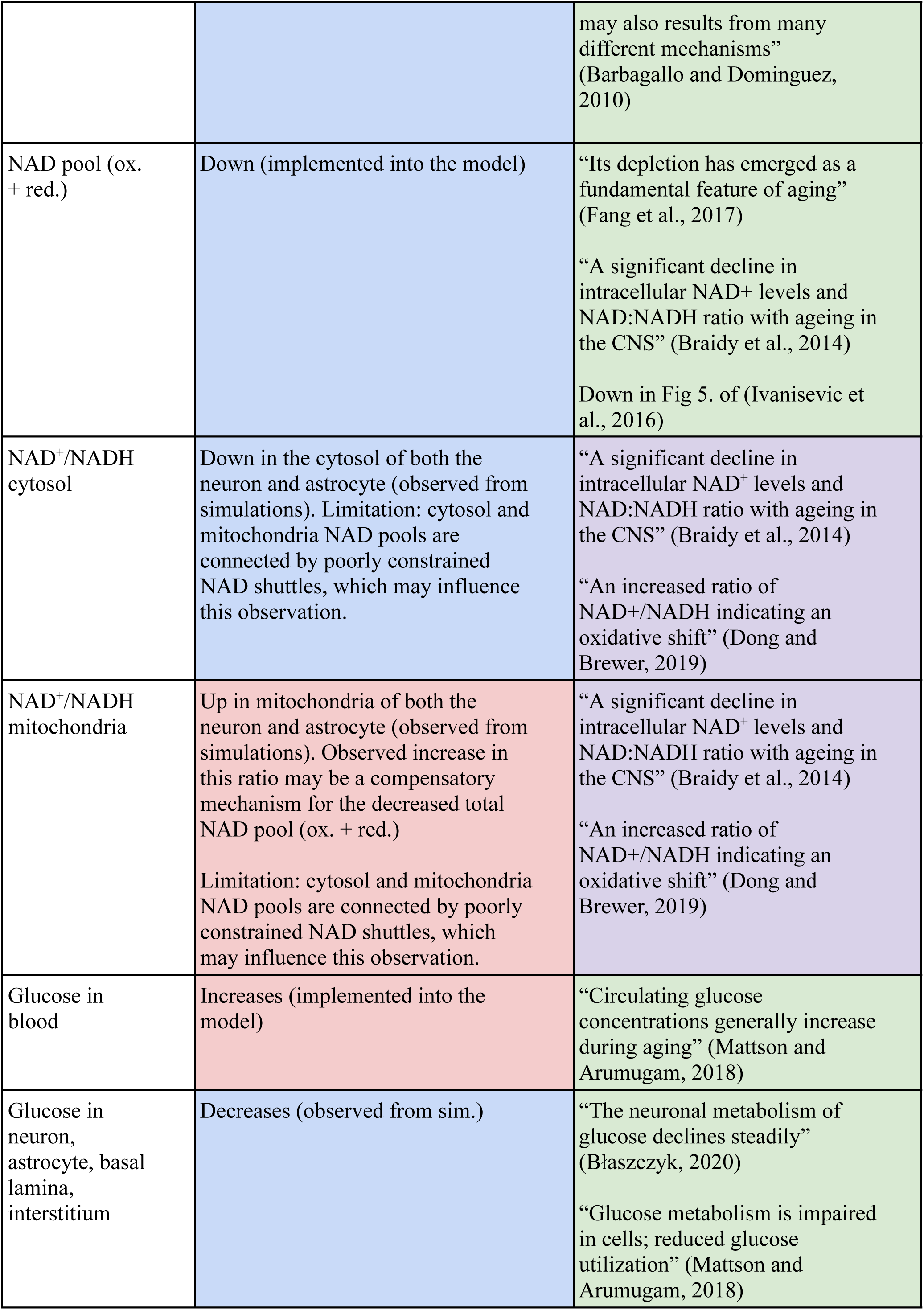
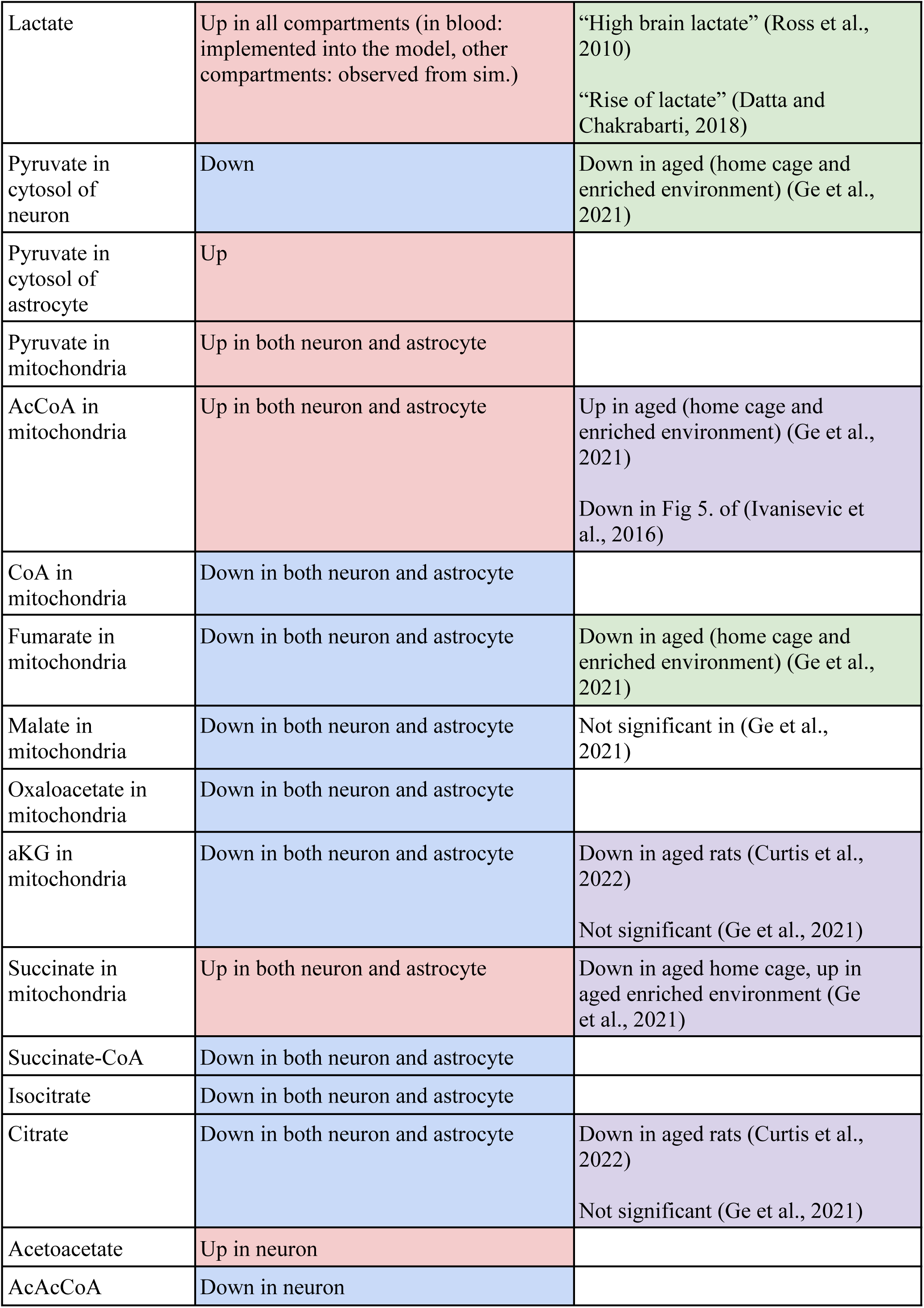
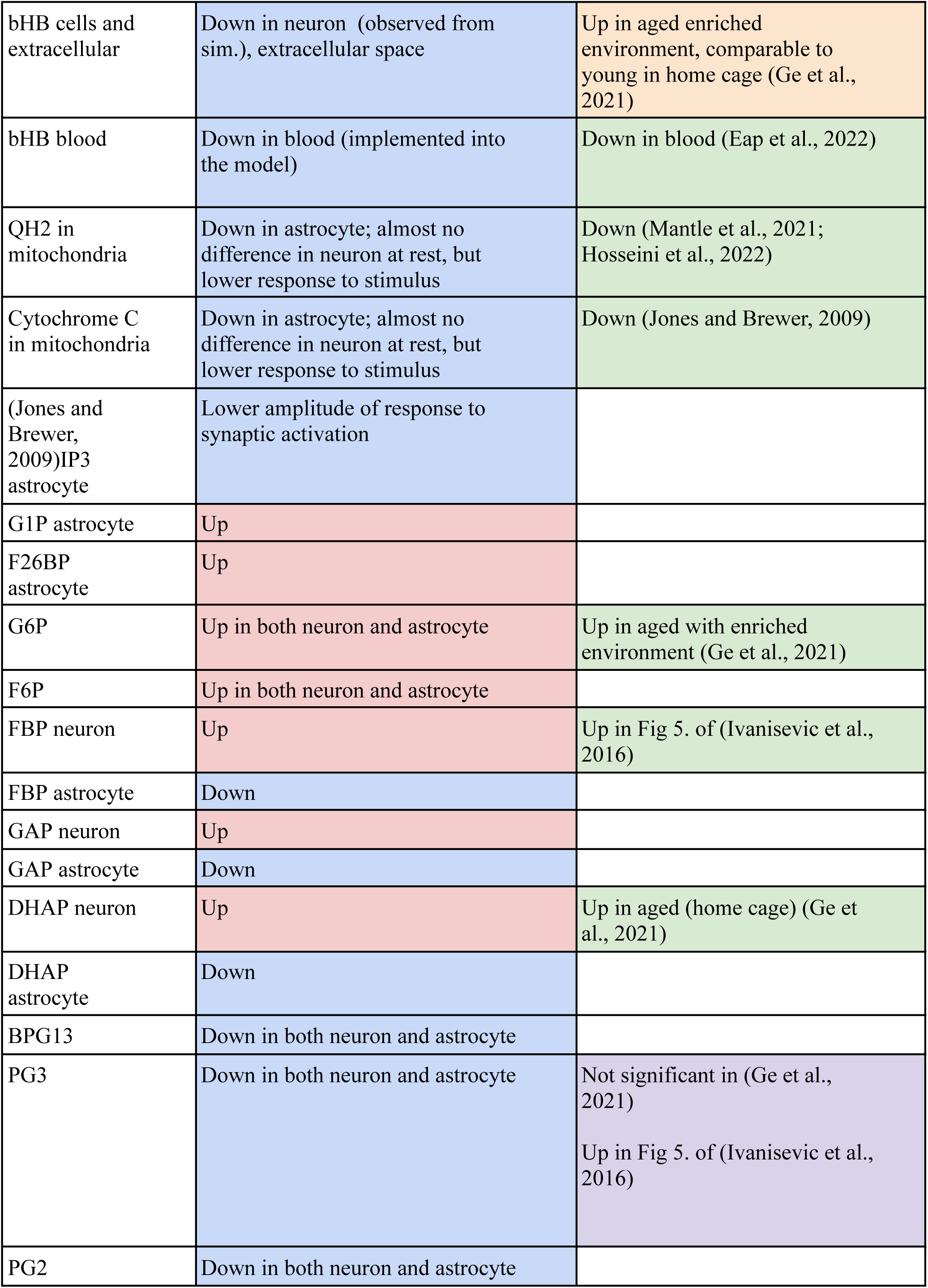
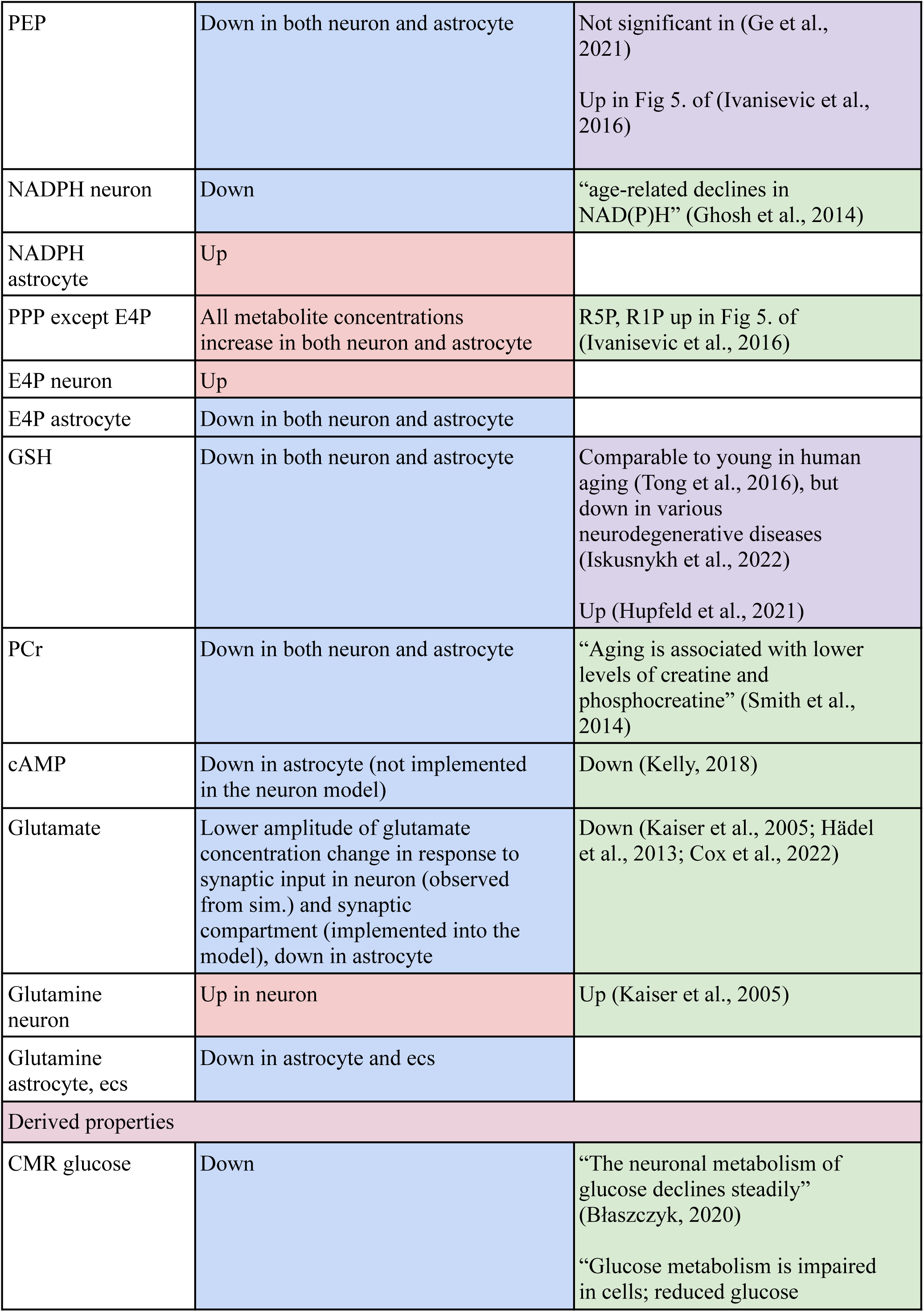
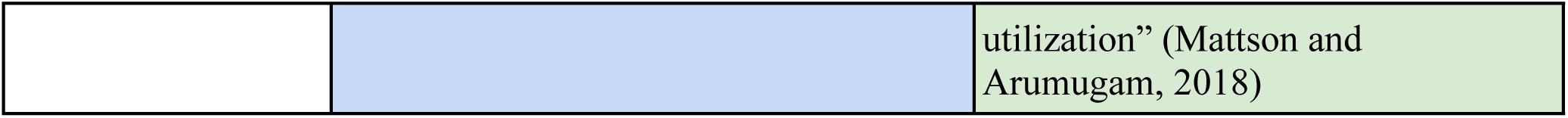
Observed aging effects and their comparison to the literature. Decrease in aging is highlighted by blue background, increase in aging is highlighted by red; consistency with literature is highlighted by green background, inconsistency is highlighted by orange background; literature data from different sources providing contradictory evidence is highlighted by purple background.

**Supplementary Table 2.**
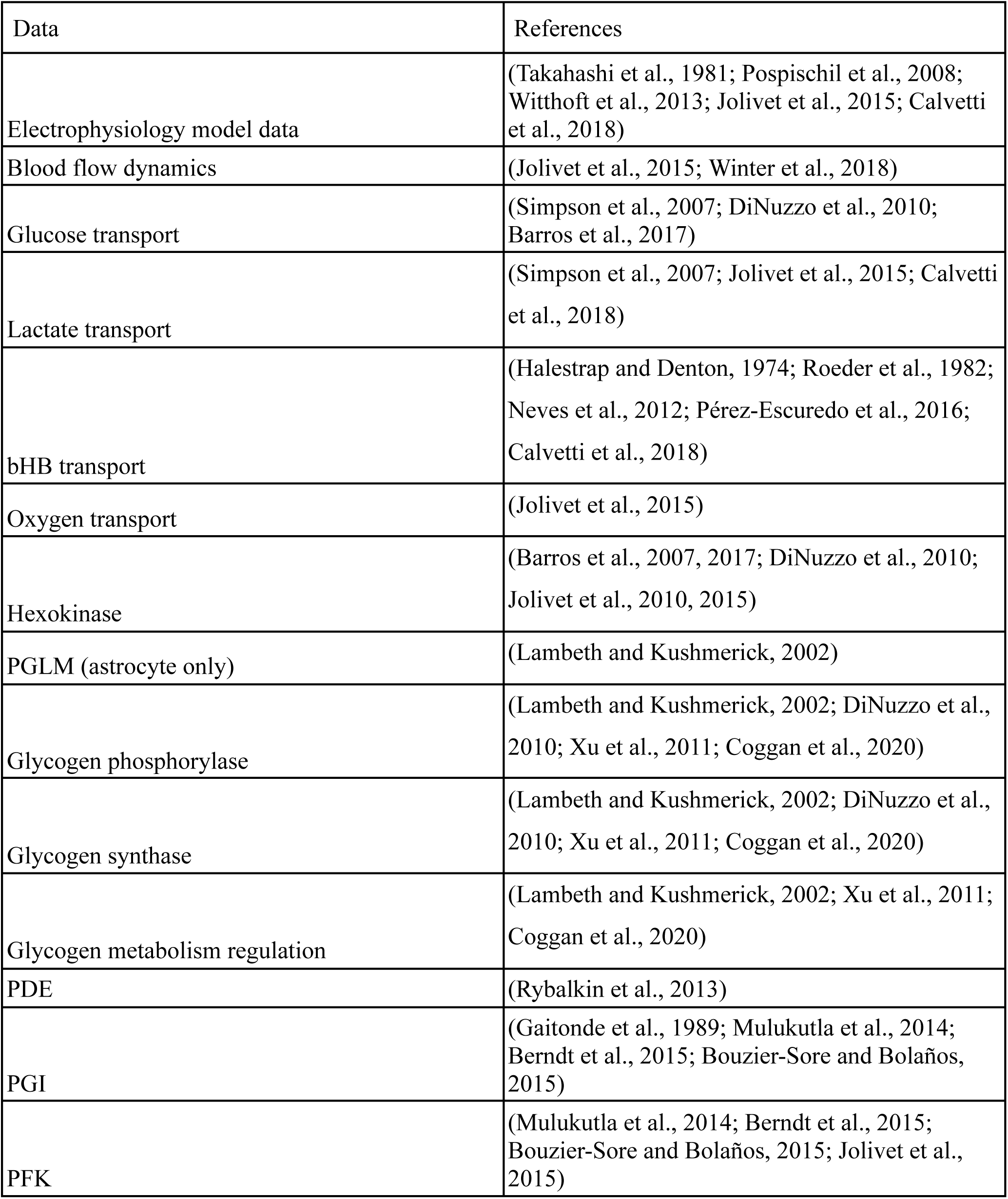
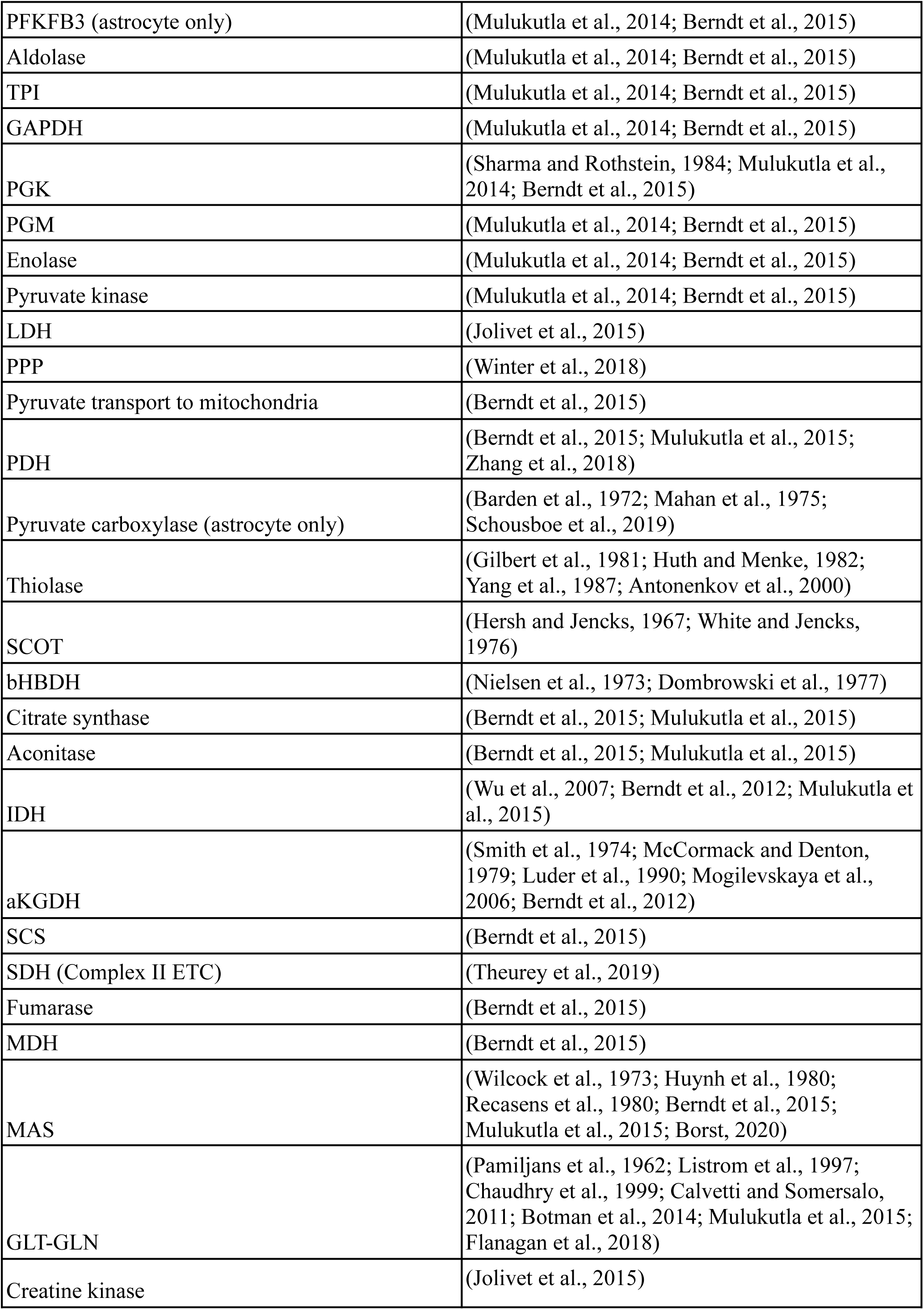
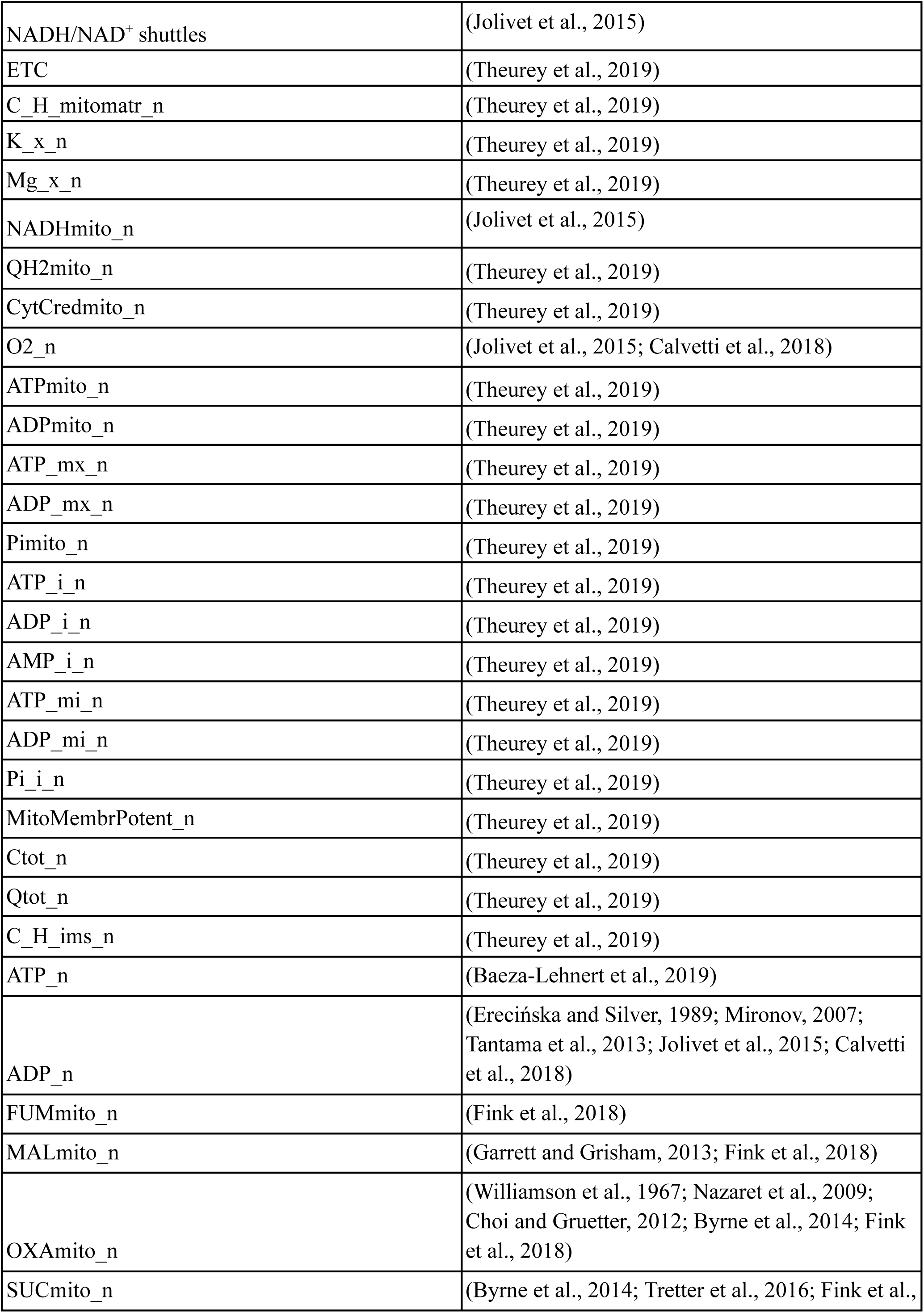
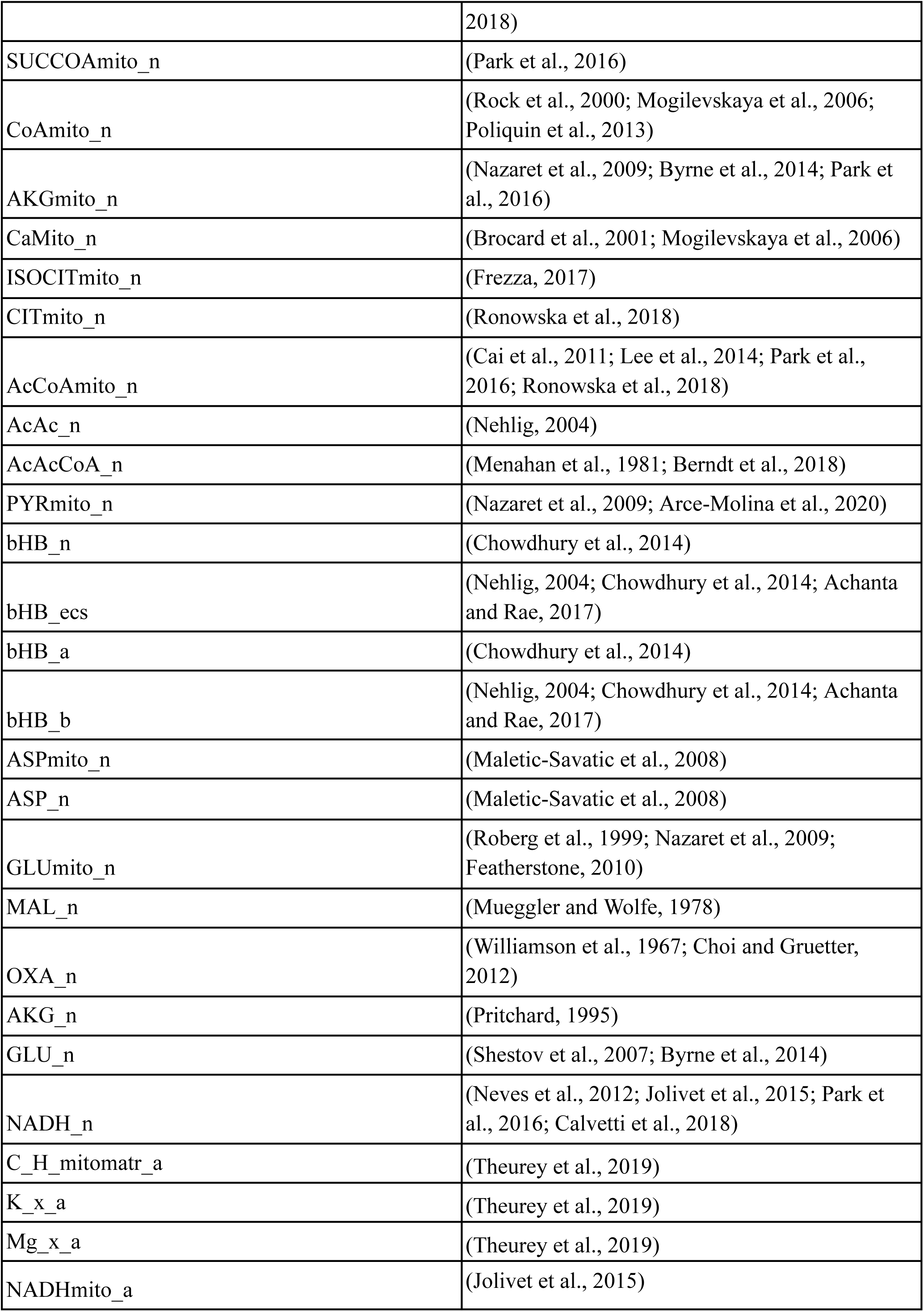
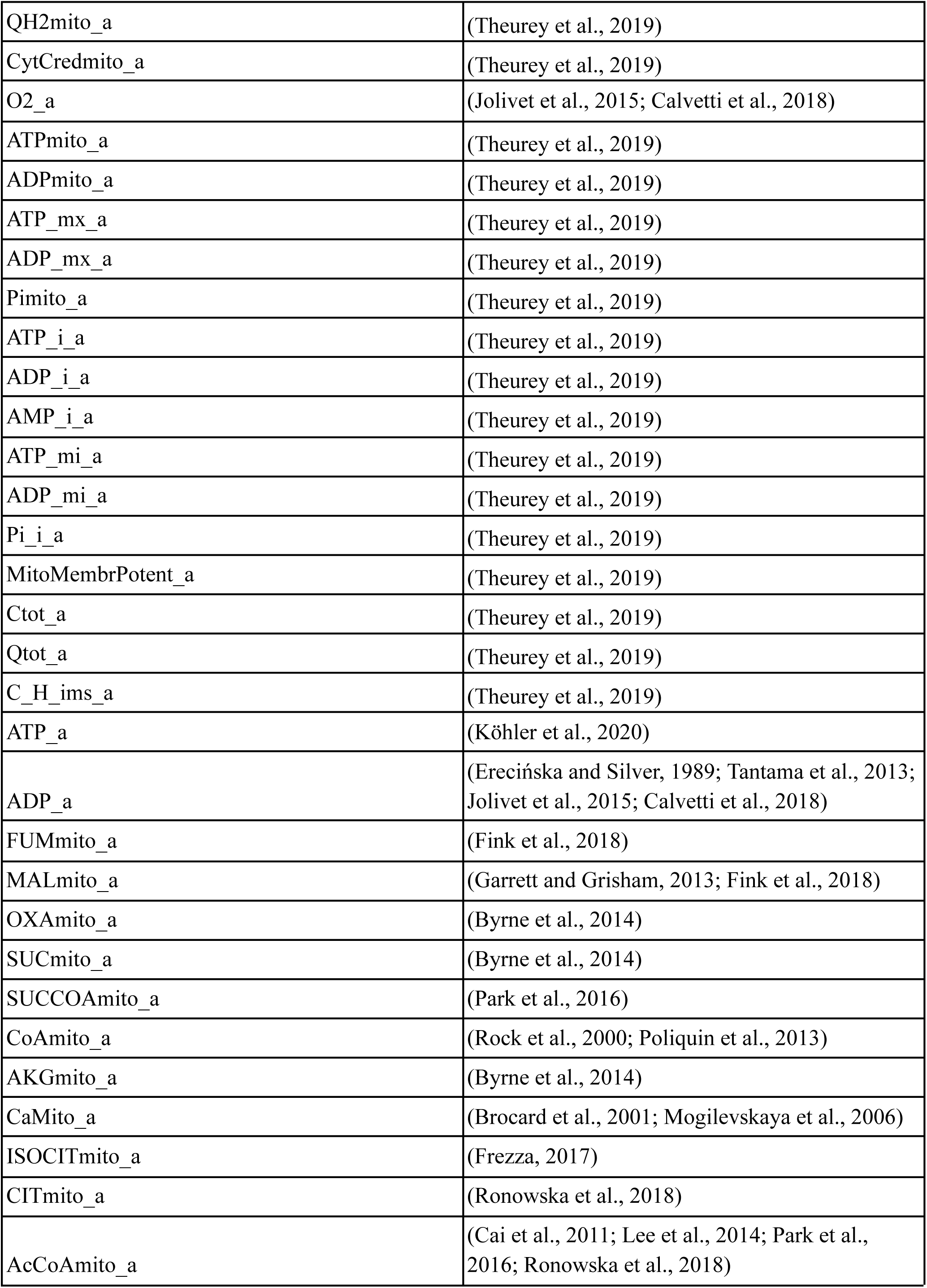
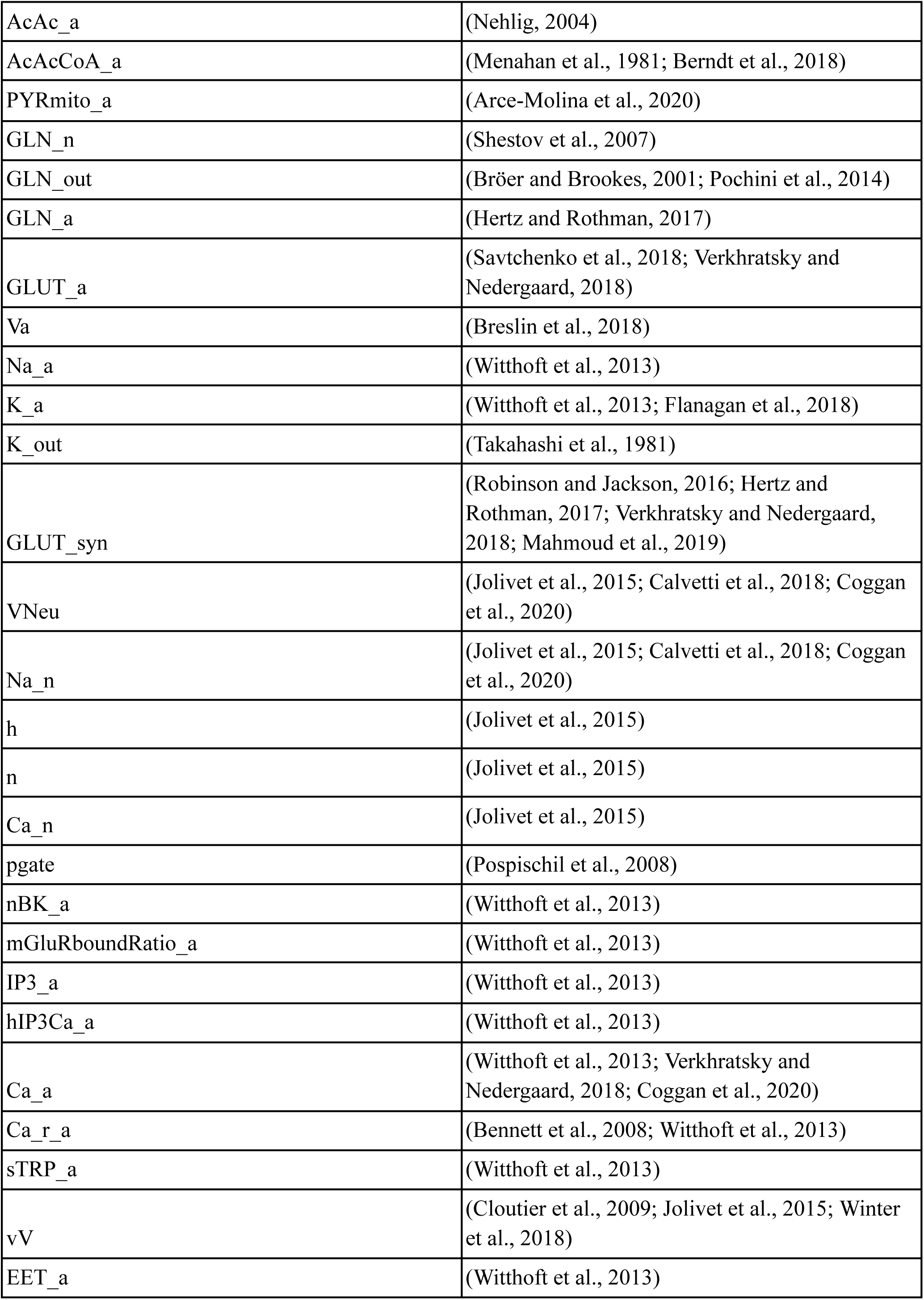
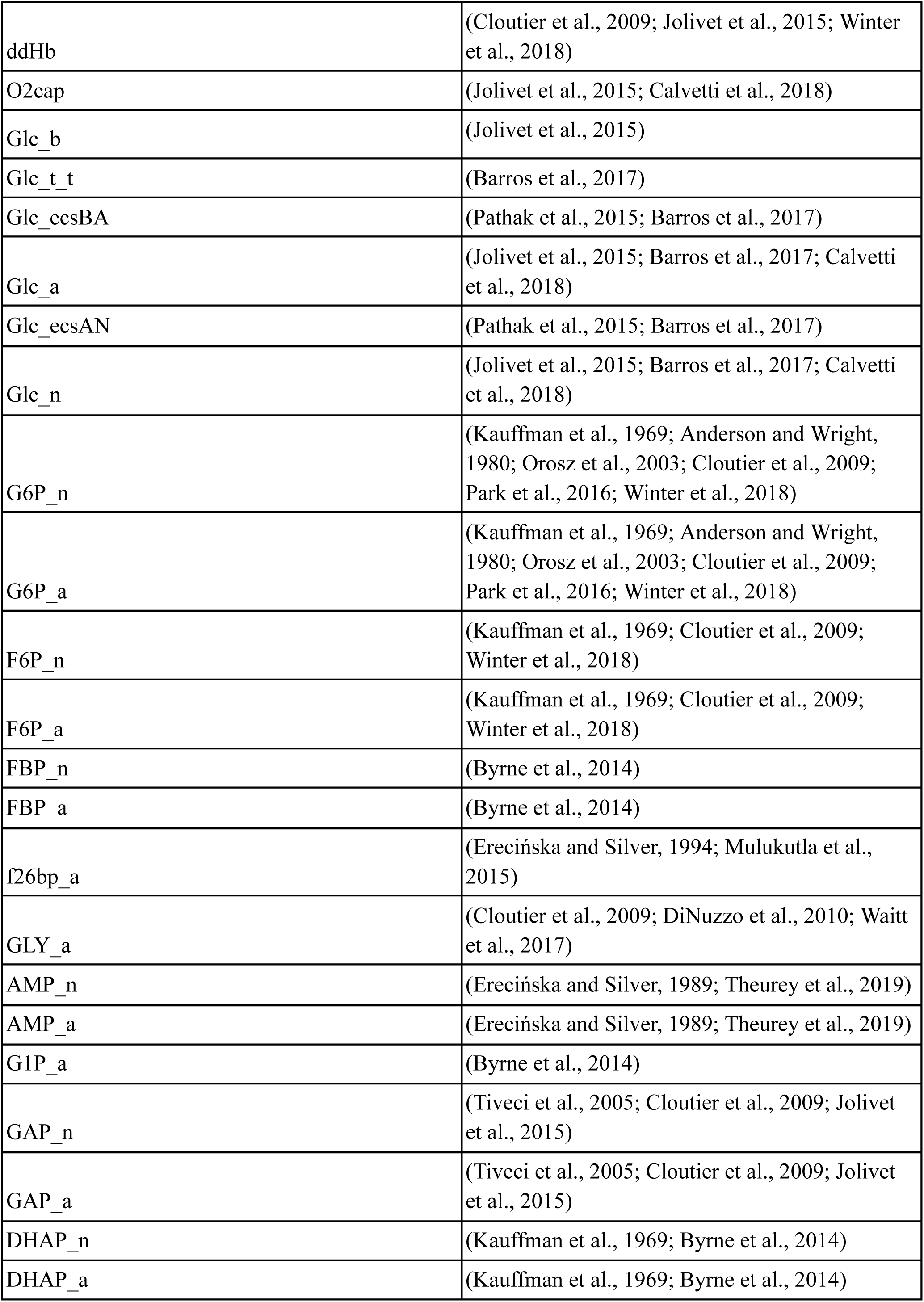
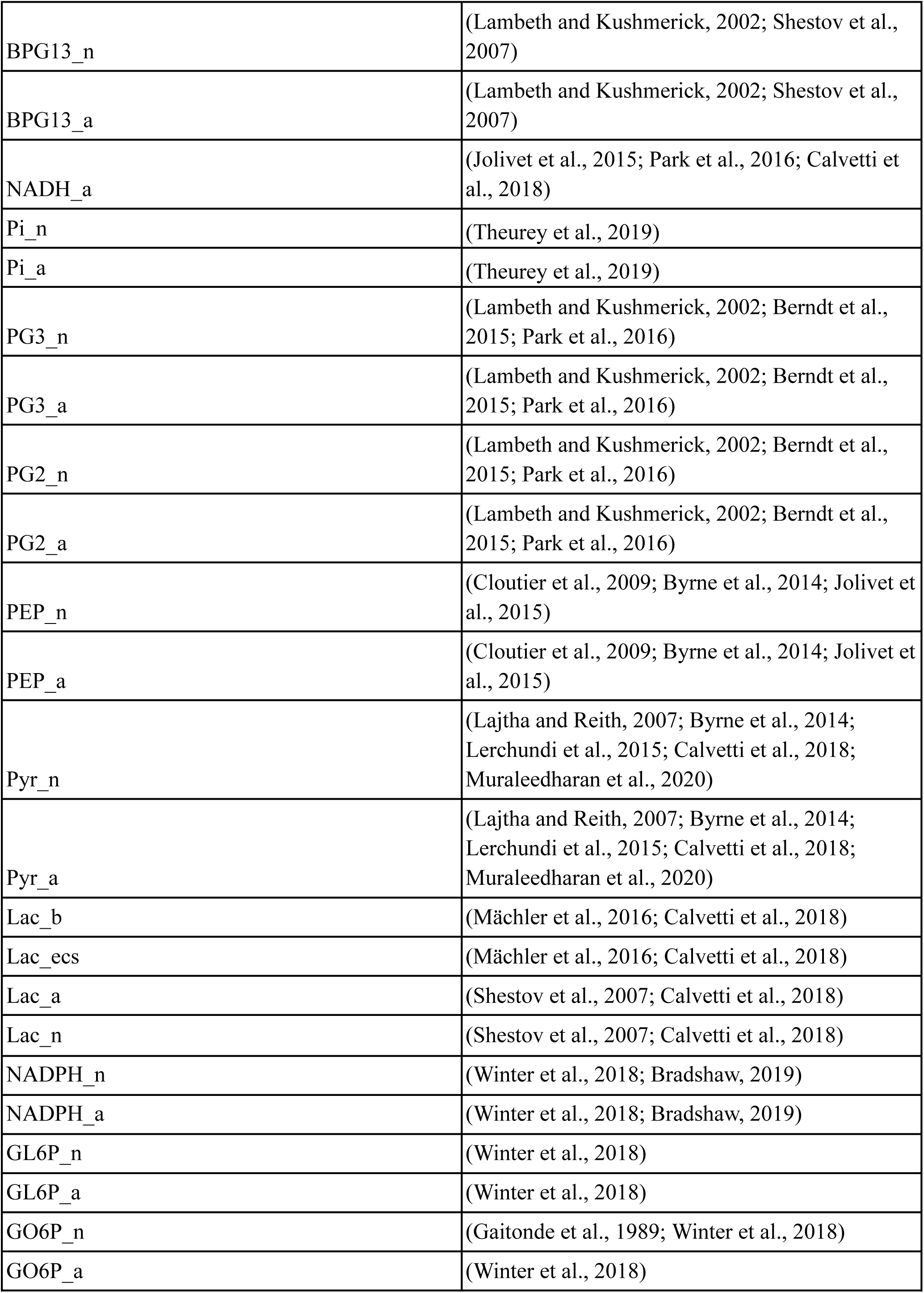
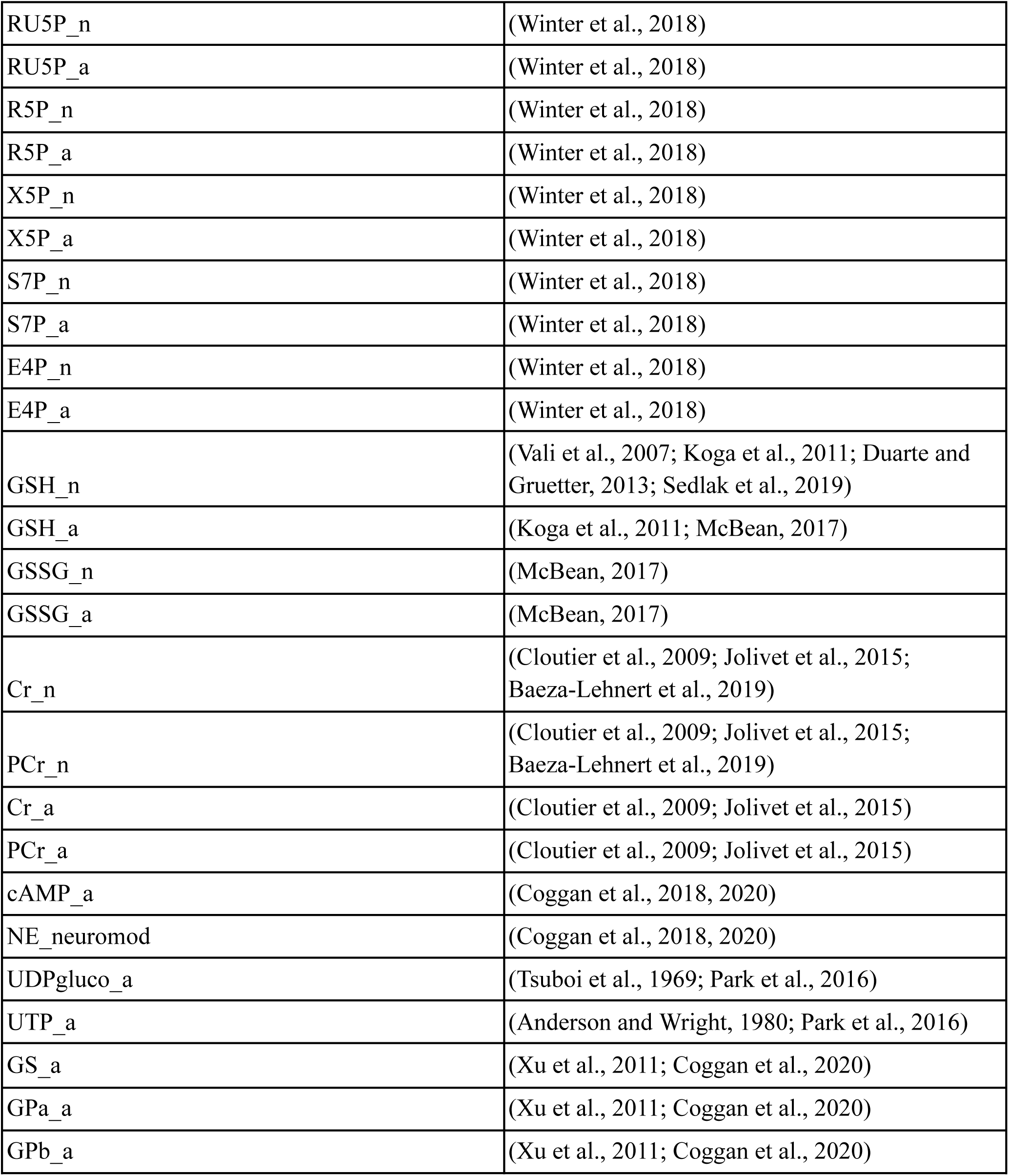
Data sources.

**Supplementary Table 3.** Anti-aging optimisation results.

|  |  |  |
| --- | --- | --- |
| Therapy (detailed list of parameters for each case is in antiage_opt/prep_opt.ipynb) | File ID | Fitness (lower is better) |
| 01 DEN-therapy<br>(diet represented by blood glucose and beta-hydroxybutyrate levels, exercise effects represented by blood lactate levels, NAD supplementation represented by total NAD, and NADH cytosol-mitochondria shuttle capacity modulation) | opt01_dietNadSuppShuttles_0508_1716_FITNESS | 324.6946224064965 |
| 02 Diet and exercise (diet represented by blood glucose and beta-hydroxybutyrate levels, exercise effects represented by blood lactate levels) | opt02_diet_0508_1736_FITNESS | 330.8649909859092 |
| 03 Diet and exercise + targeted selection of enzymes and transporters (diet represented by blood glucose and beta-hydroxybutyrate levels, exercise effects represented by blood lactate levels, selection of top-vulnerable enzyme and transporter targets based on sensitivity analysis) | opt03_sel2diet_0508_1745_FITNESS | 330.5296867114816 |
| 04 Diet and exercise + targeted selection of enzymes and transporters + top transcription factor regulated targets<br>(as therapy-03 above, but with additional set of parameters representing ESRRA regulated targets) | opt04_sel2diettopTF_0508_1749_FITNESS | 329.6195317070475 |
| 05 Diet and exercise + NAD supplementation + NADH cytosol-mitochondria shuttle capacity modulation + top transcription factor regulated targets<br>(diet represented by blood glucose and beta-hydroxybutyrate levels, exercise effects represented by blood lactate levels, NAD supplementation represented by total NAD, NADH cytosol-mitochondria shuttle capacity modulation, ESRRA regulated targets) | opt05_dietNadSuppShuttlestopTF_0508_1754_FITNESS | 321.4015401204615 |
| 06 Diet and exercise + NAD supplementation + top transcription factor regulated targets<br>(as therapy-05, but without NADH | opt06_dietNadSupptopTF_0508_1800_FITNESS | 326.52278564010174 |
| cytosol-mitochondria shuttle capacity modulation) |  |  |
| 07 Diet + exercise + NAD supplementation<br>(as 01 DEN-therapy, but without NADH cytosol-mitochondria shuttle capacity modulation) | opt07_dietNadSupp_0508_1805_FITNESS | 330.9287687071004 |
| 08 Diet and exercise + NAD supplementation + NADH cytosol-mitochondria shuttle capacity modulation + targeted selection of enzymes and transporters + top transcription factor regulated targets<br>(as therapy-04, but additionally with NAD supplementation represented by total NAD, NADH cytosol-mitochondria shuttle capacity modulation) | opt08_sel3fulltopTF_0508_1809_FITNESS | 322.00660105095636 |
| 09 Diet and exercise + NAD supplementation + NADH cytosol-mitochondria shuttle capacity modulation + targeted selection of enzymes and transporters<br>(as therapy-08, but without ESRRA regulated targets) | opt09_sel3full_0508_1814_FITNESS | 321.46368653203 |
| 10 NAD supplementation + NADH cytosol-mitochondria shuttle capacity modulation + targeted selection of enzymes and transporters + top transcription factor regulated targets<br>(as therapy-08, but without diet and exercise) | opt10_sel2nadfulltopTF_0508_1817_FITNESS | 325.9331563280608 |
| **11 Targeted selection of enzymes and transporters**<br>**+ NAD supplementation + NADH cytosol-mitochondria shuttle capacity modulation** | **opt11_sel2nadfull_0508_1818_FITNESS** | **320.2230577011635** |
| 12 NADH cytosol-mitochondria shuttle capacity modulation | opt12_NADshuttles_cm_0508_1832_FITNESS | 325.33028862691594 |
| 13 Diet and exercise + NAD supplementation + targeted selection of enzymes and transporters + top transcription factor regulated targets<br>(as therapy-08, but without NADH | opt13_sel3topTF_0508_1835_FITNESS | 332.51439560199486 |
| cytosol-mitochondria shuttle capacity modulation) |  |  |
| 14 Diet and exercise + NAD supplementation + targeted selection of enzymes and transporters (as therapy-13, but without top transcription factor regulated targets) | opt14_sel3_0508_1837_FITNESS | 330.8959136554414 |
| 15 NAD supplementation + targeted selection of enzymes and transporters + top transcription factor regulated targets (as therapy-13, but without diet and exercise) | opt15_sel2nadtopTF_0508_1838_FITNESS | 333.0797288342428 |
| 16 NAD supplementation + targeted selection of enzymes and transporters | opt16_sel2nad_0508_1840_FITNESS | 323.72448628397404 |
| 17 NAD supplementation represented by total NAD | opt17_NADsupplements_0508_1841_FITNESS | 325.9386339659428 |
| 18 Targeted selection of enzymes and transporters + top transcription factor (ESRRA) regulated targets | opt18_sel1topTF_0508_1844_FITNESS | 329.8385866817131 |
| 19 Targeted selection of enzymes and transporters (based on sensitivity analysis) | opt19_sel1_0508_1847_FITNESS | 330.32612175950476 |
| 20 Top transcription factor (ESRRA) regulated targets | opt20_sel_topTF_0508_1848_FITNESS | 330.564190608832 |

### Supplementary Figures

**Supplementary Fig. 1.**
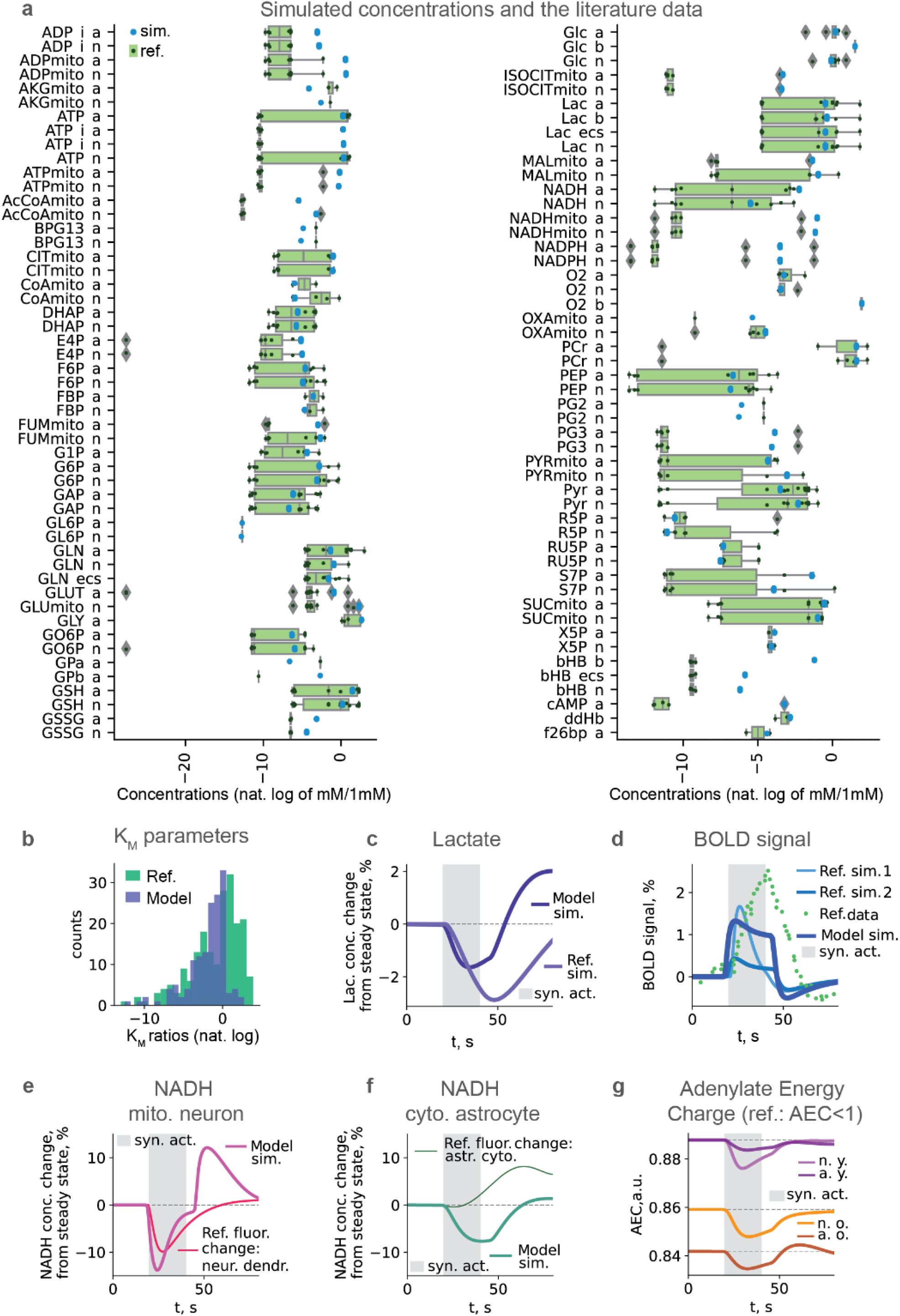
Validation, predicted energy budget. **a,** Simulated concentrations and the literature data. **b,** Comparison of ratios of model KM to average mammalian data KM for the same pairs enzyme-metabolite to min/mean and max/mean ratios of that mammalian data for the same pairs enzyme-metabolite, **c,** Lactate dynamics compared to simulation of the other model from the literature (Jolivet et al., 2015). **d,** BOLD dynamics (Jolivet et al., 2015; Winter et al., 2018; Jung et al., 2021). **e,** NADH dynamics in neuronal mitochondria compared to literature (Jolivet et al., 2015). **f,** NADH dynamics in astrocyte cytosol compared to literature (Jolivet et al., 2015). **g,** ATP consumption per AP in young and old ages.

**Supplementary Fig. 2.**
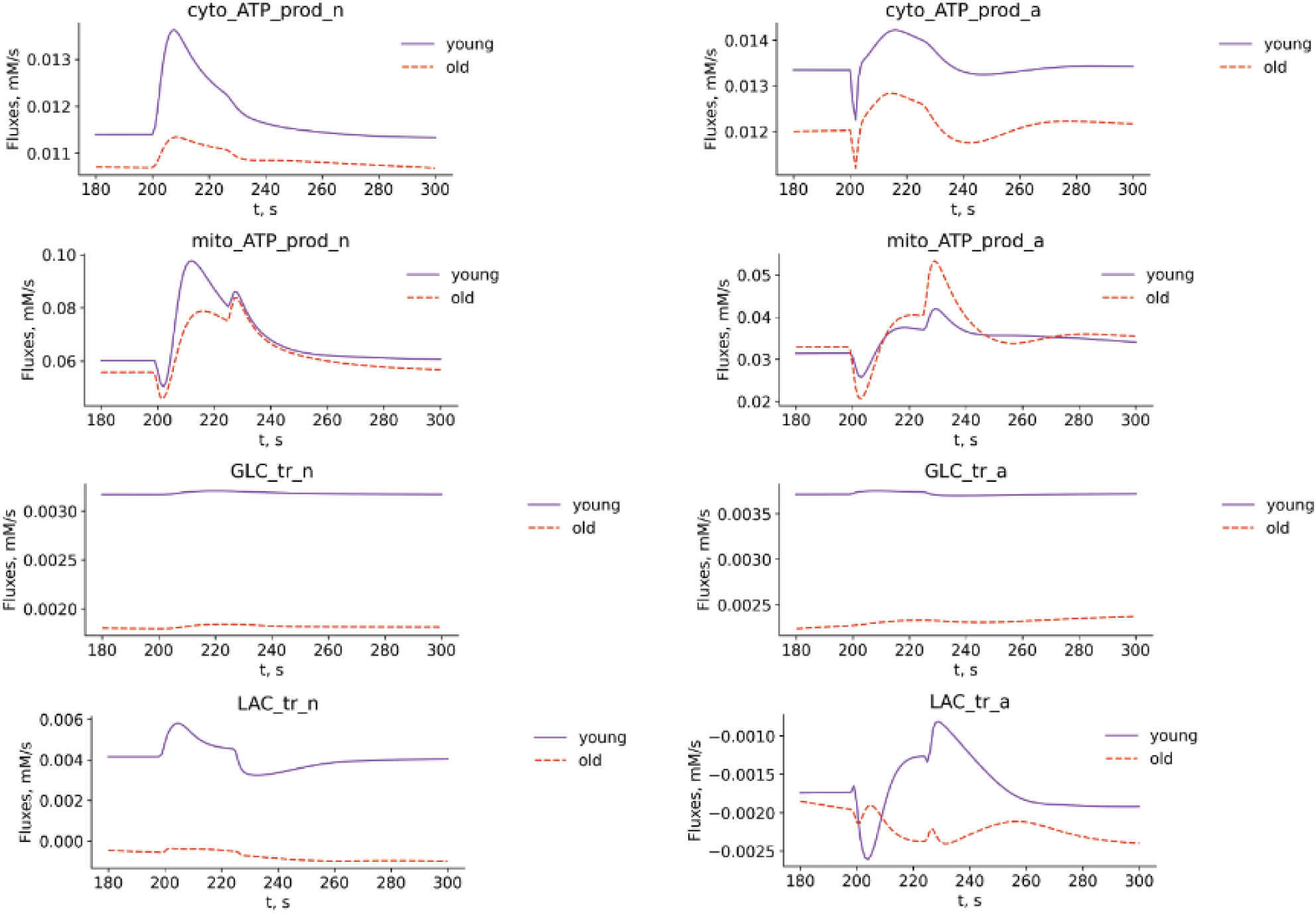
ATP production, glucose and lactate transport fluxes.

**Supplementary Fig. 3.**
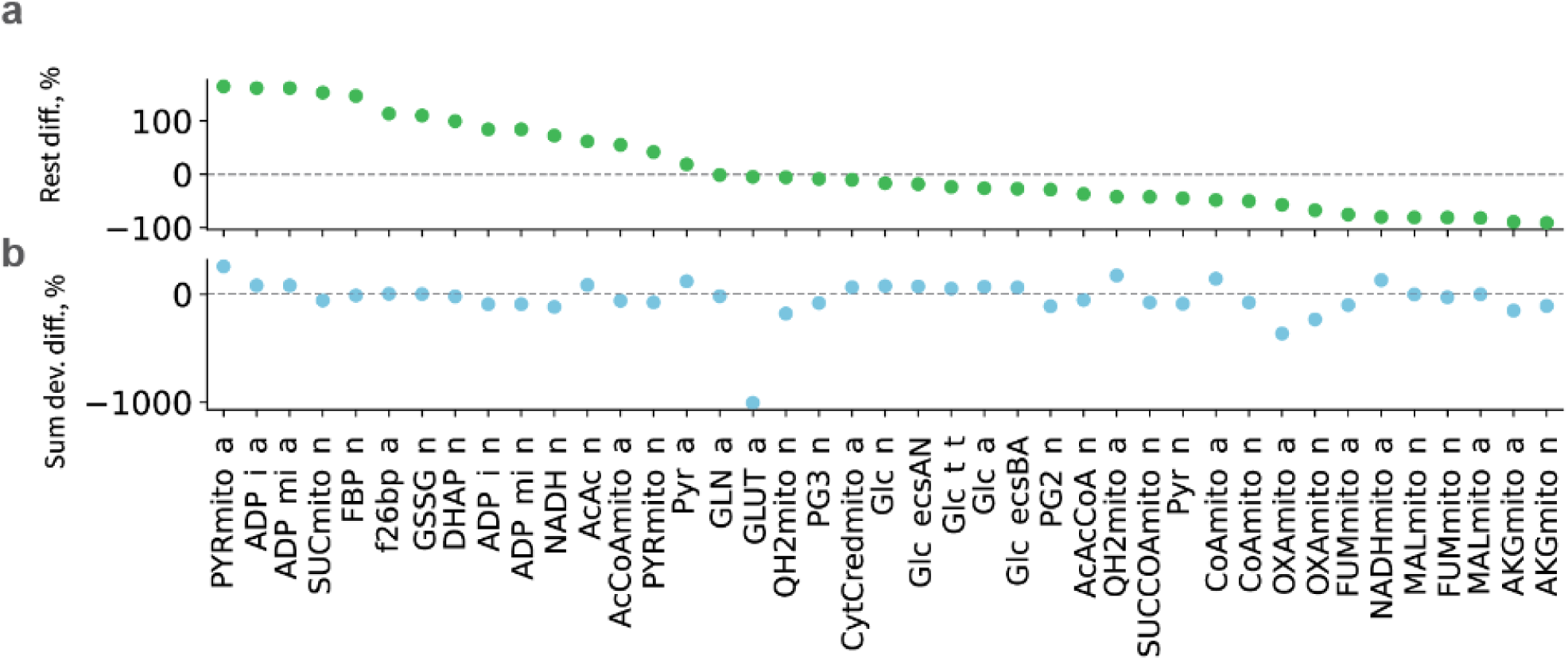
Differences between young and old in rest state concentrations (top) and in sum of relative deviations of concentration from rest (normalized by rest state) upon synaptic activation (bottom), both ranked by rest state differences (top), only top ranked are shown.

**Supplementary Fig. 4.**
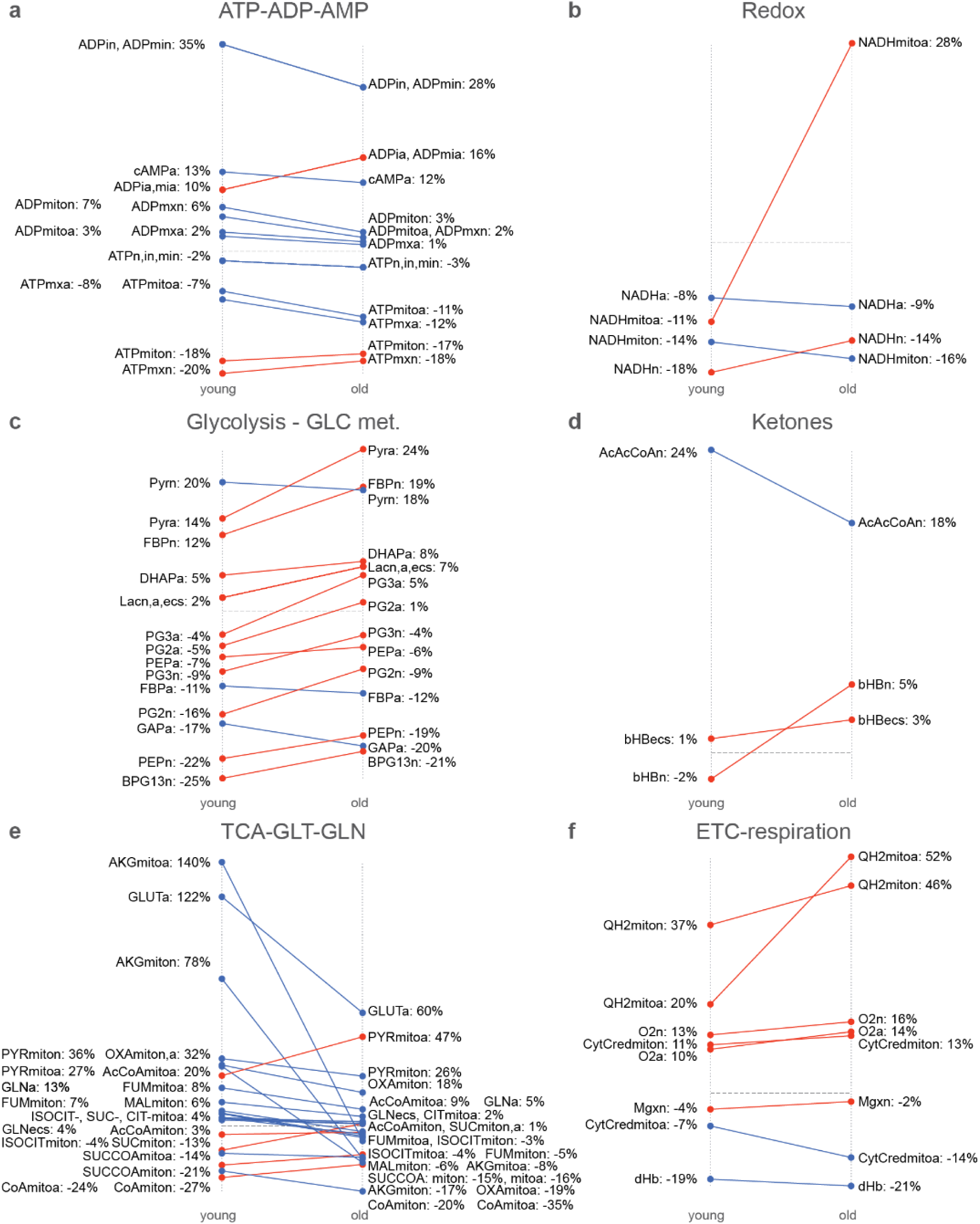
Comparison of amplitudes of metabolic response to synaptic activation in young and old ages (filtered by absolute values of deviations and difference in deviations of higher than 1%).

**Supplementary Fig. 5.**
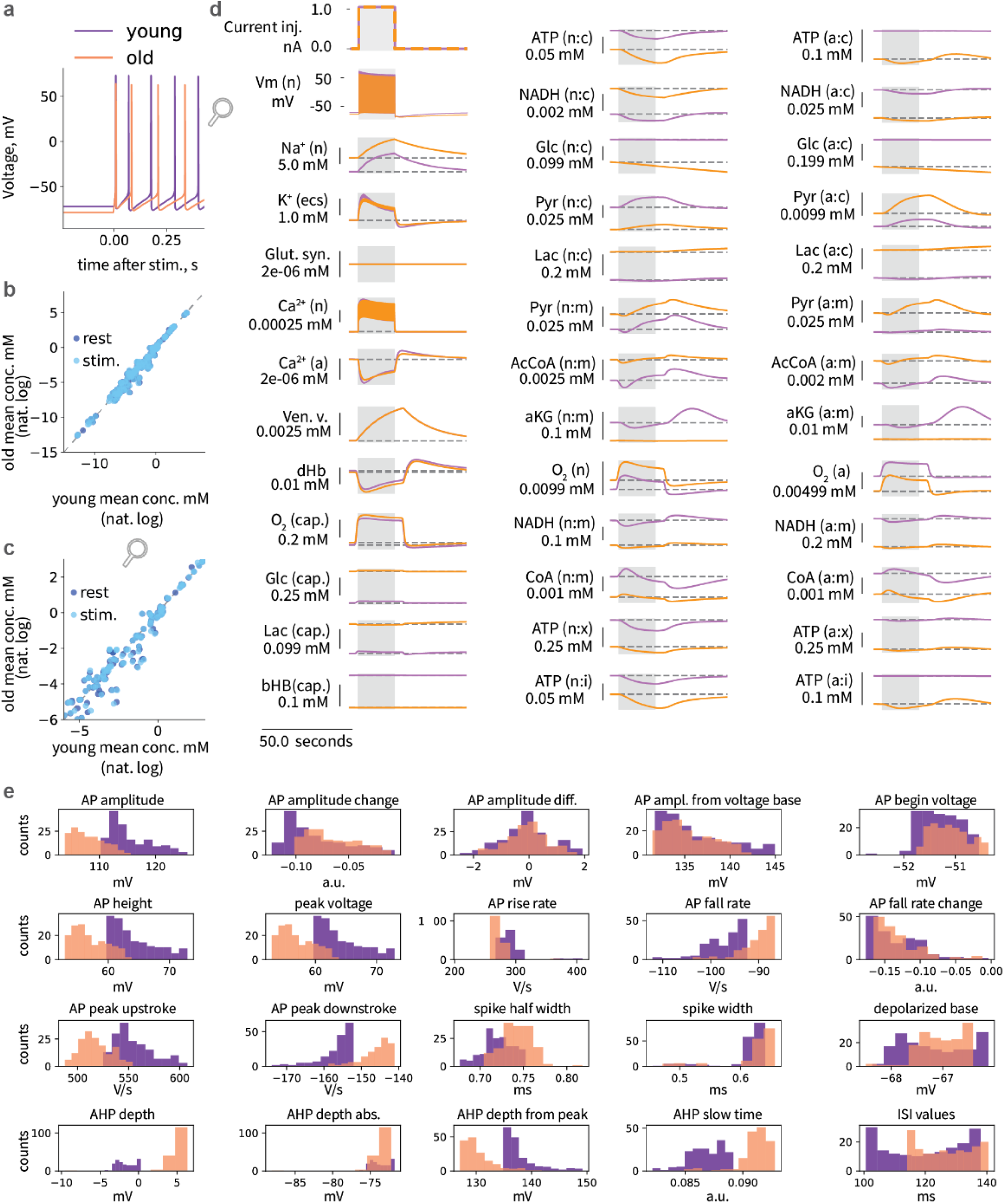
Train of APs evoked by 1 nA current injection simulations. **a,** Dynamics of metabolism in response to a train of APs evoked by current injection in different ages. **b,** Characteristics of neuronal firing in young and old ages evoked by current injection.

**Supplementary Fig. 6.**
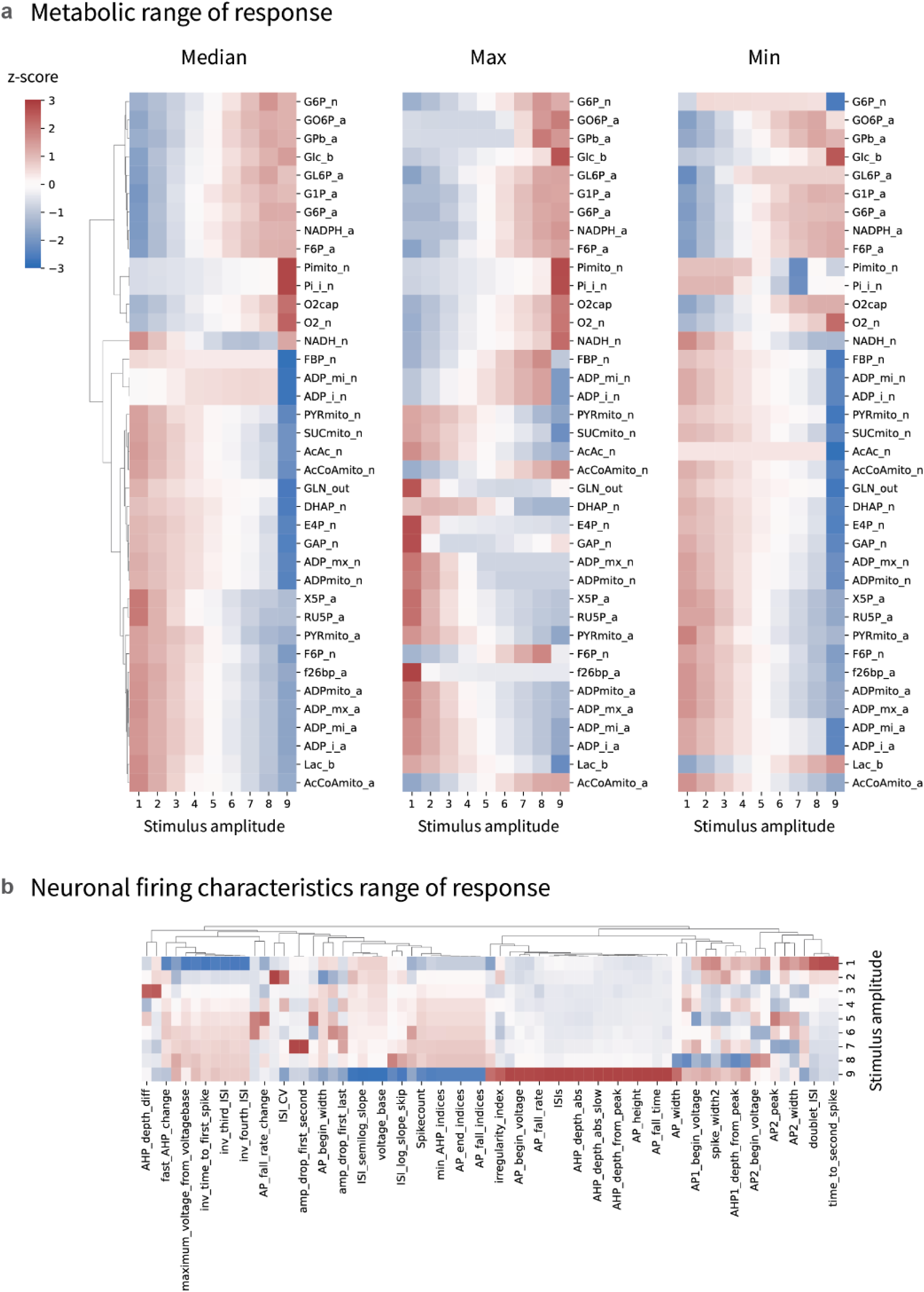
Aging-associated differences in range of response to the current injections of different amplitudes.

**Supplementary Fig. 7.**
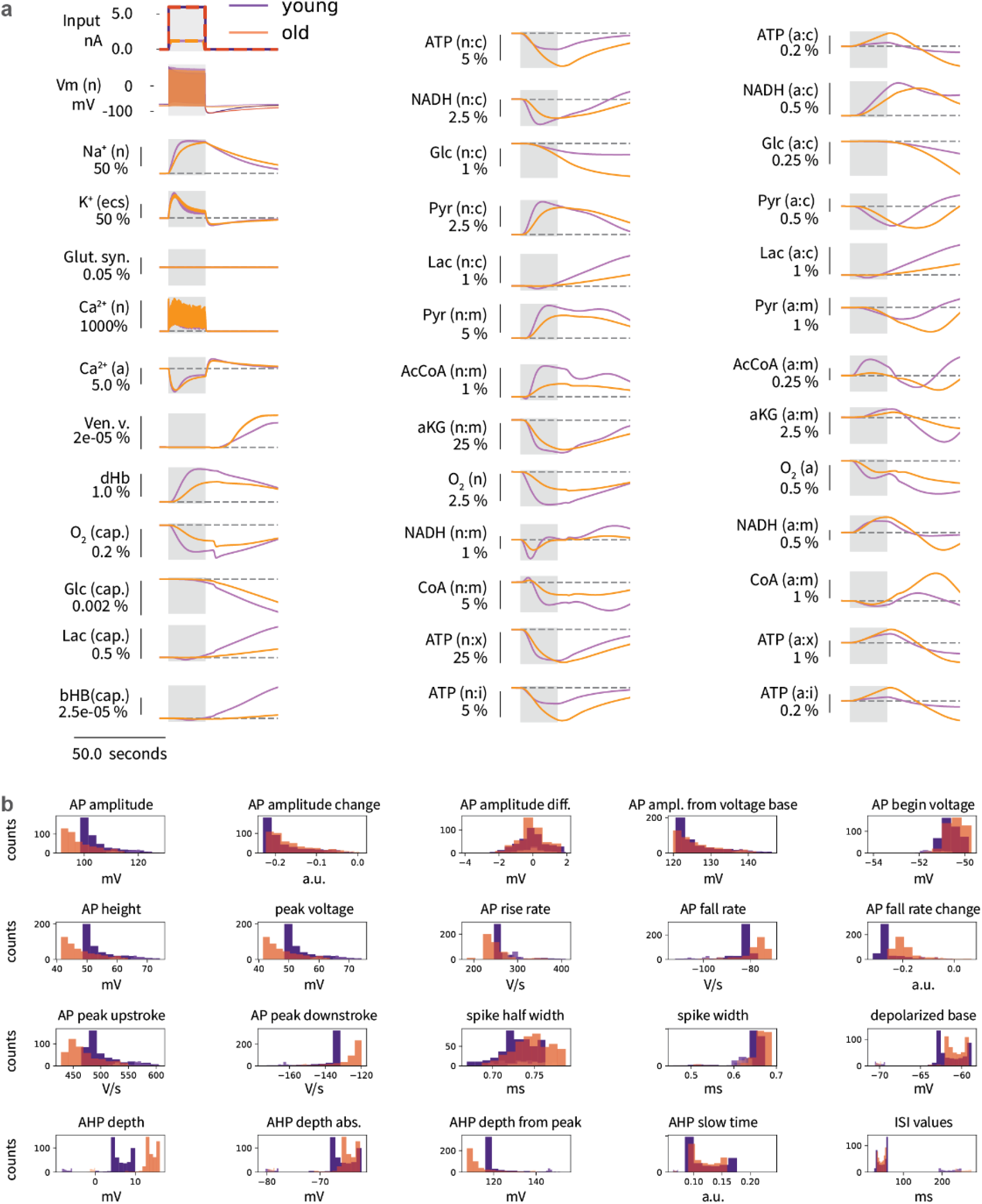
Dependence of metabolism and electrophysiology responses on the current injection amplitude in young and old ages. **a,** Young and old age responses to current injections of two different amplitudes: input current (top left in A), firing traces (left on second row in A), percent difference in metabolic response: 100*(m_IinjHigh - m_IinjLow)/m_IinjLow (all other figures in A). **b,** Characteristics of neuronal firing in young and old ages upon current injections of two different amplitudes.

**Supplementary Fig. 8.**
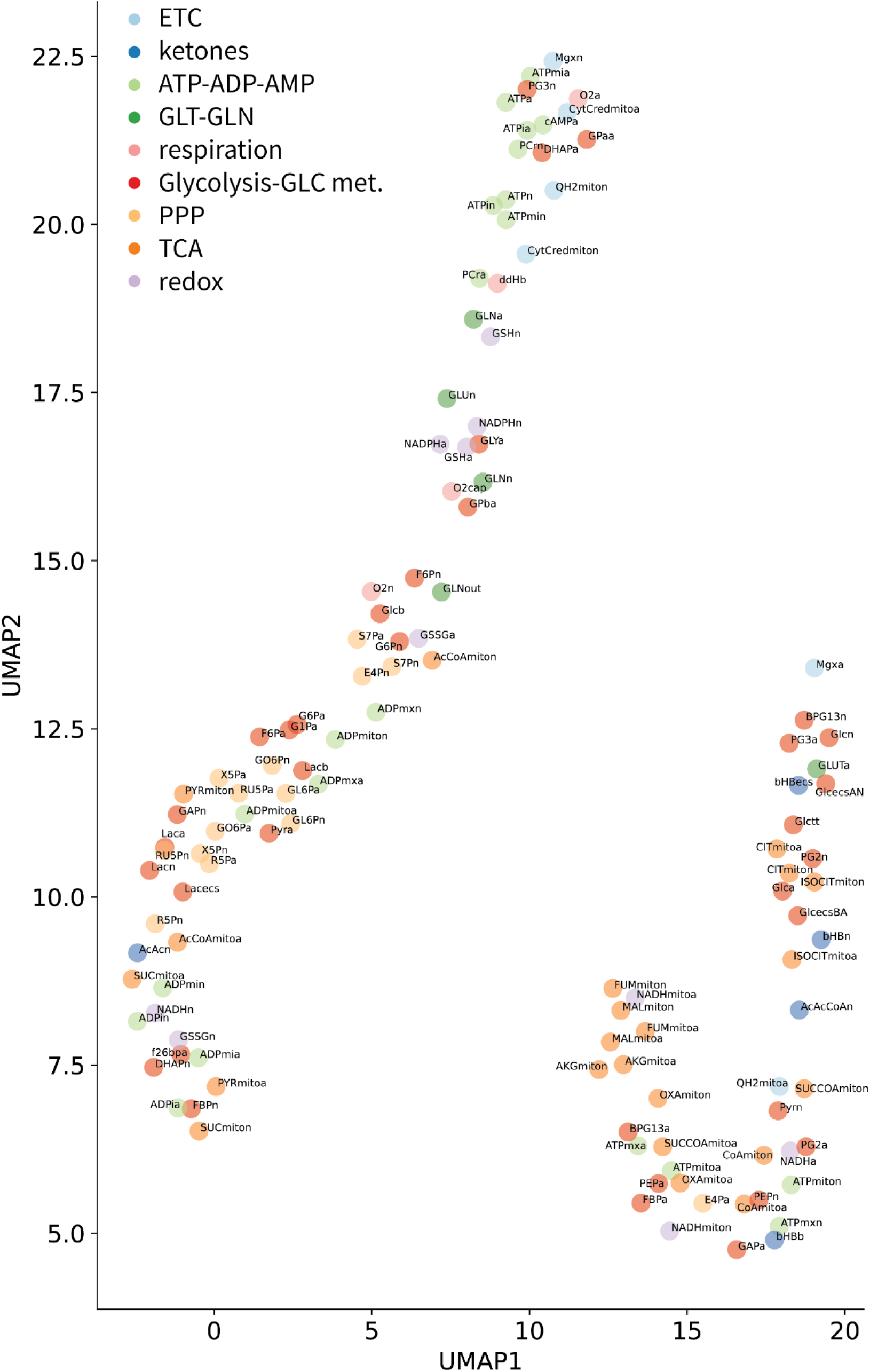
UMAP of relative differences in concentration traces in old compared to young.

**Supplementary Fig. 9.**
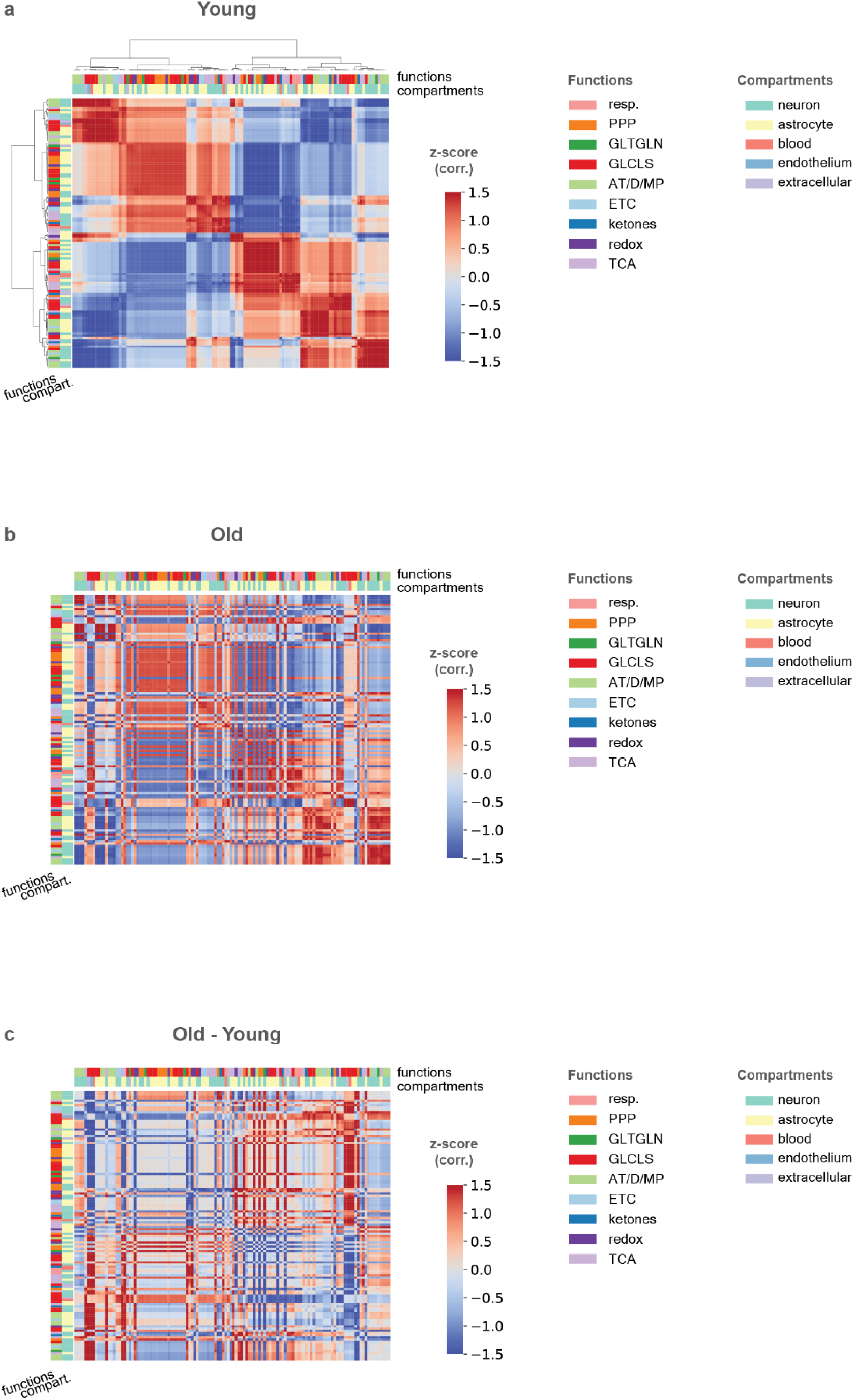
Kendall correlation of metabolite concentrations time series data in aging.

**Supplementary Fig. 10.**
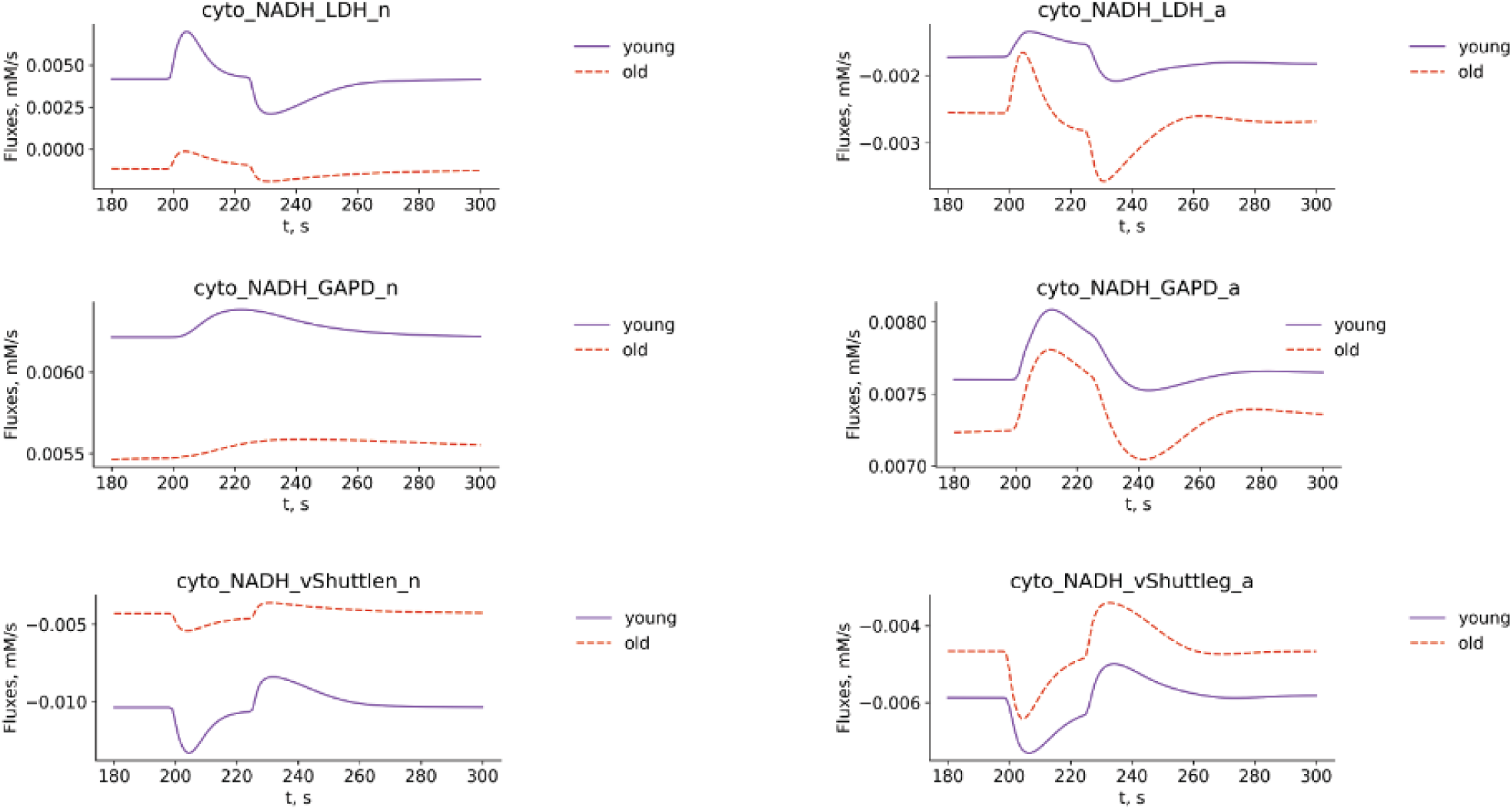
Cytosolic NADH fluxes.

**Supplementary Fig. 11.**
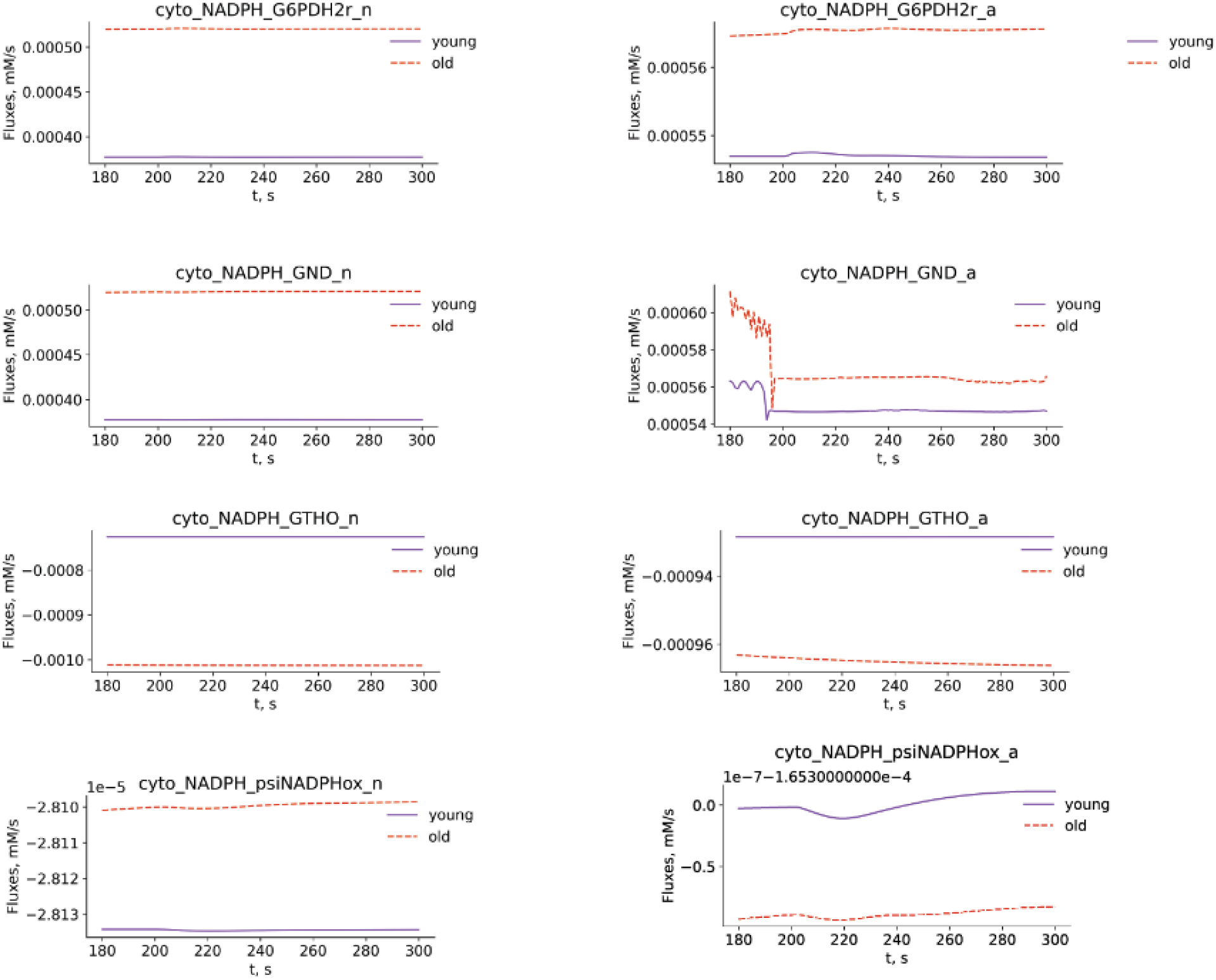
Cytosolic NADPH fluxes.

**Supplementary Fig. 12.**
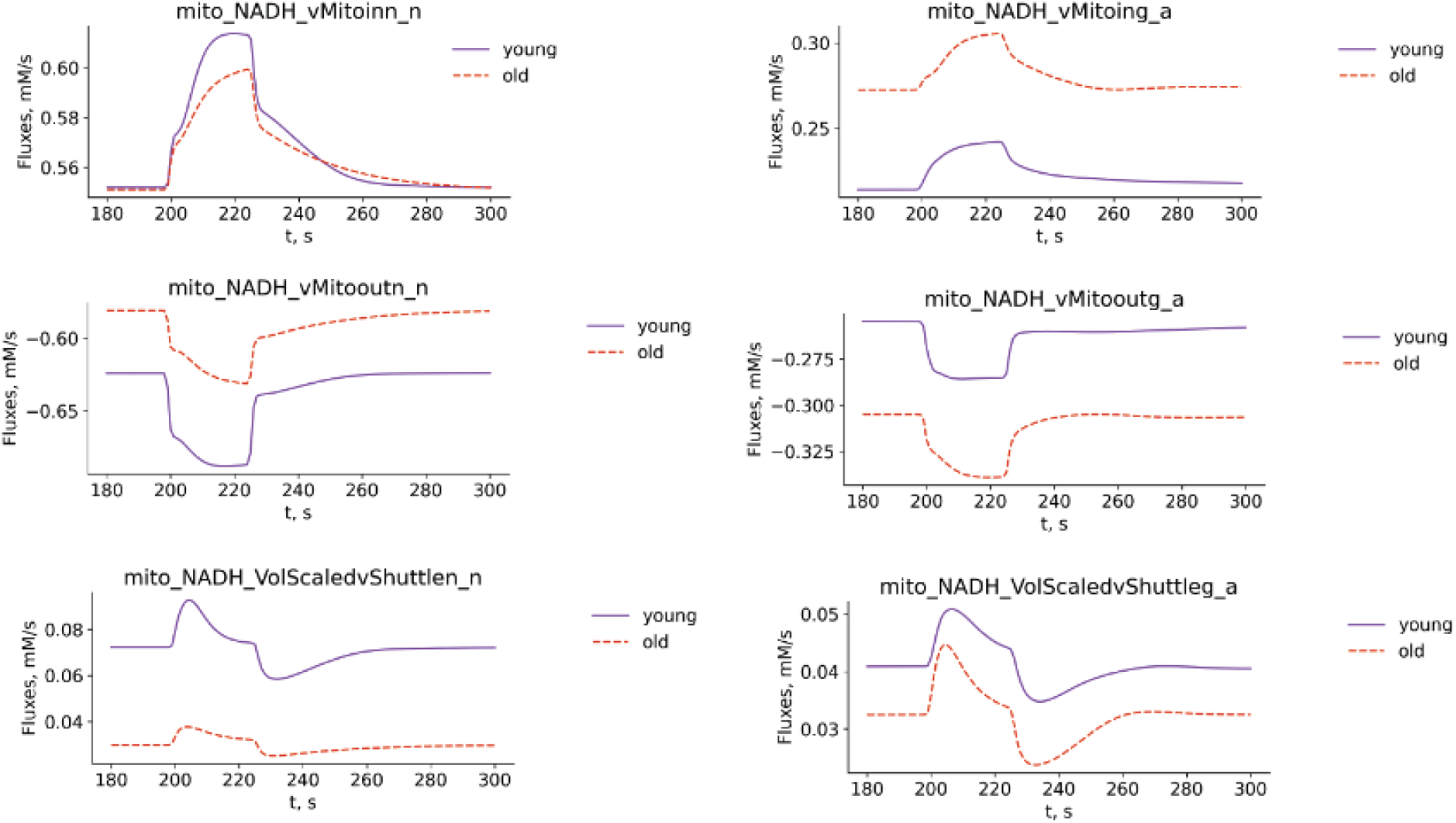
Mitochondrial NADH fluxes.

**Supplementary Fig. 13.**
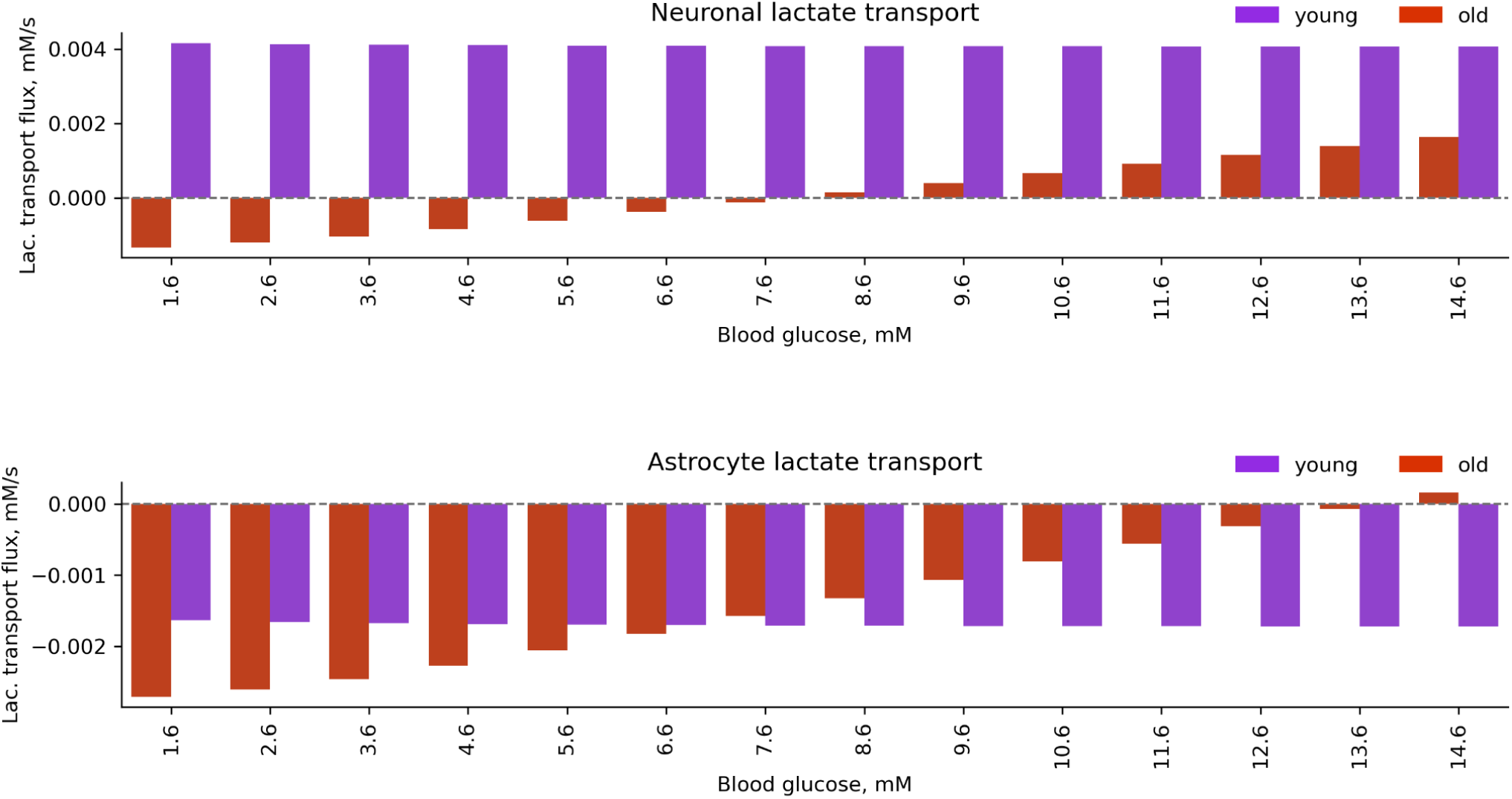
Lactate shuttle in conditions with different blood glucose levels.

**Supplementary Fig. 14.**
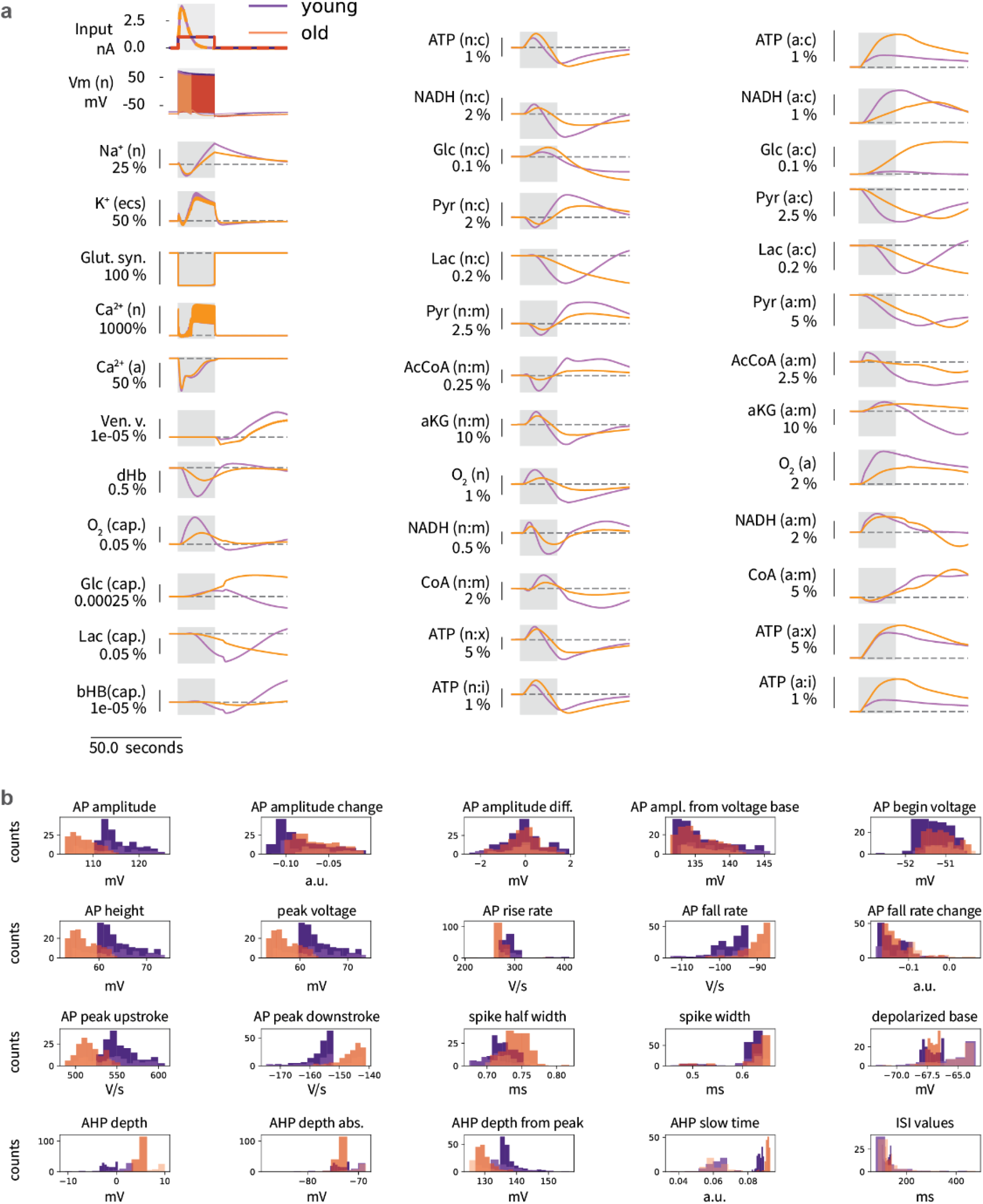
Comparison of synaptic activation and current injection evoked metabolic responses. **a,** Young and old age responses to synaptic input and current injection (approximately the same firing frequency): input current (top left in A), firing traces (left on second row in A), percent difference in metabolic response: 100*(m_Iinj - m_syn)/m_syn (all other figures in A). **b,** Characteristics of neuronal firing in young and old ages upon synaptic activation and current injection (approximately the same firing frequency).

**Supplementary Fig. 15.**
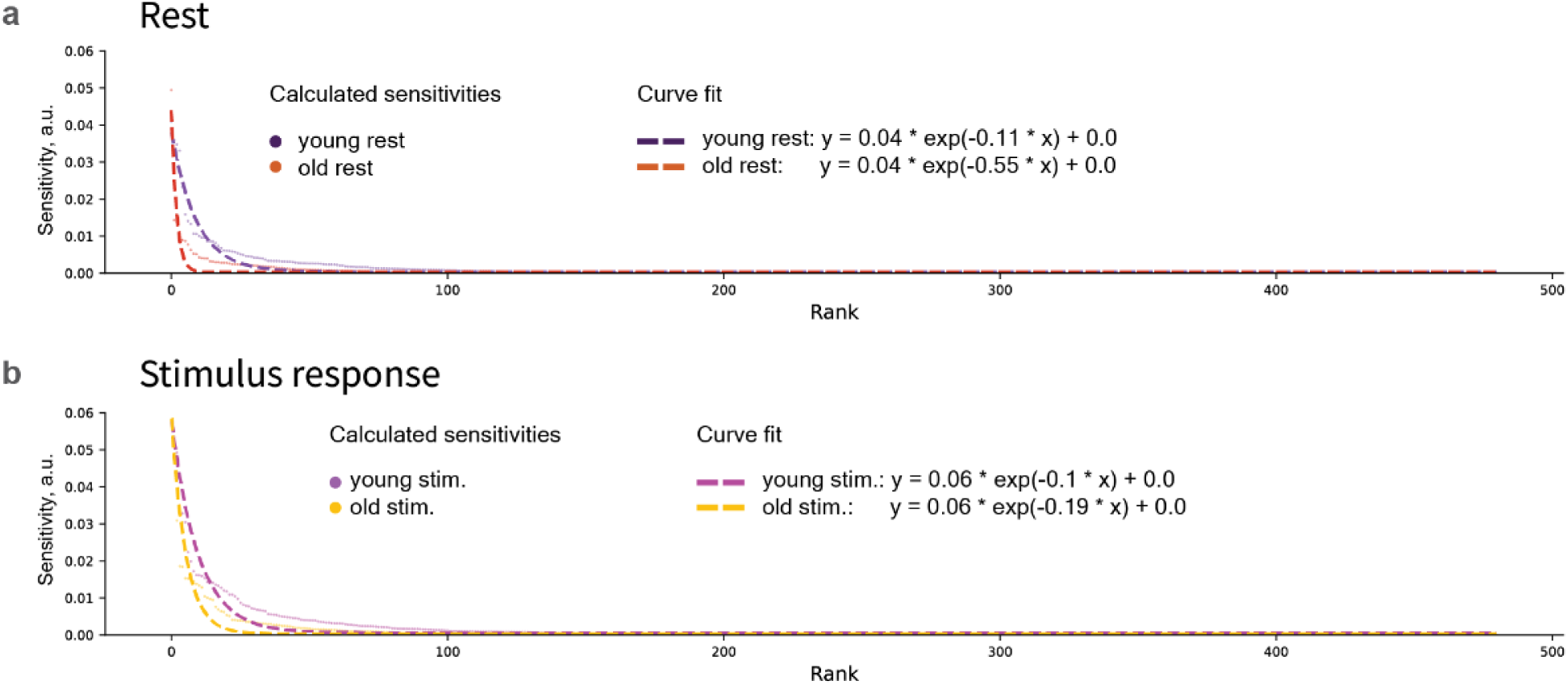
Sensitivities curve fit.

**Supplementary Fig. 16.**
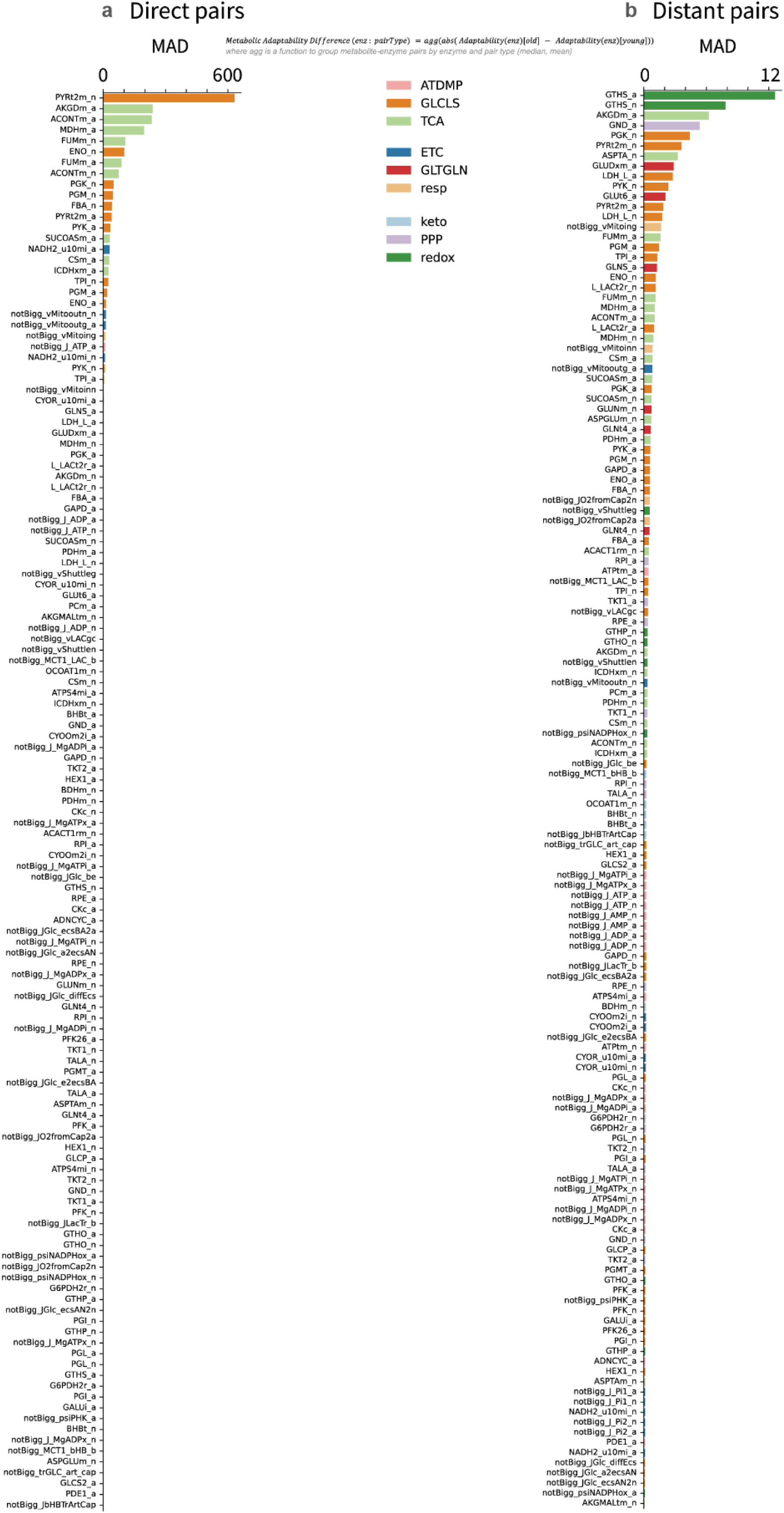
Metabolic adaptability difference.

**Supplementary Fig. 17.**
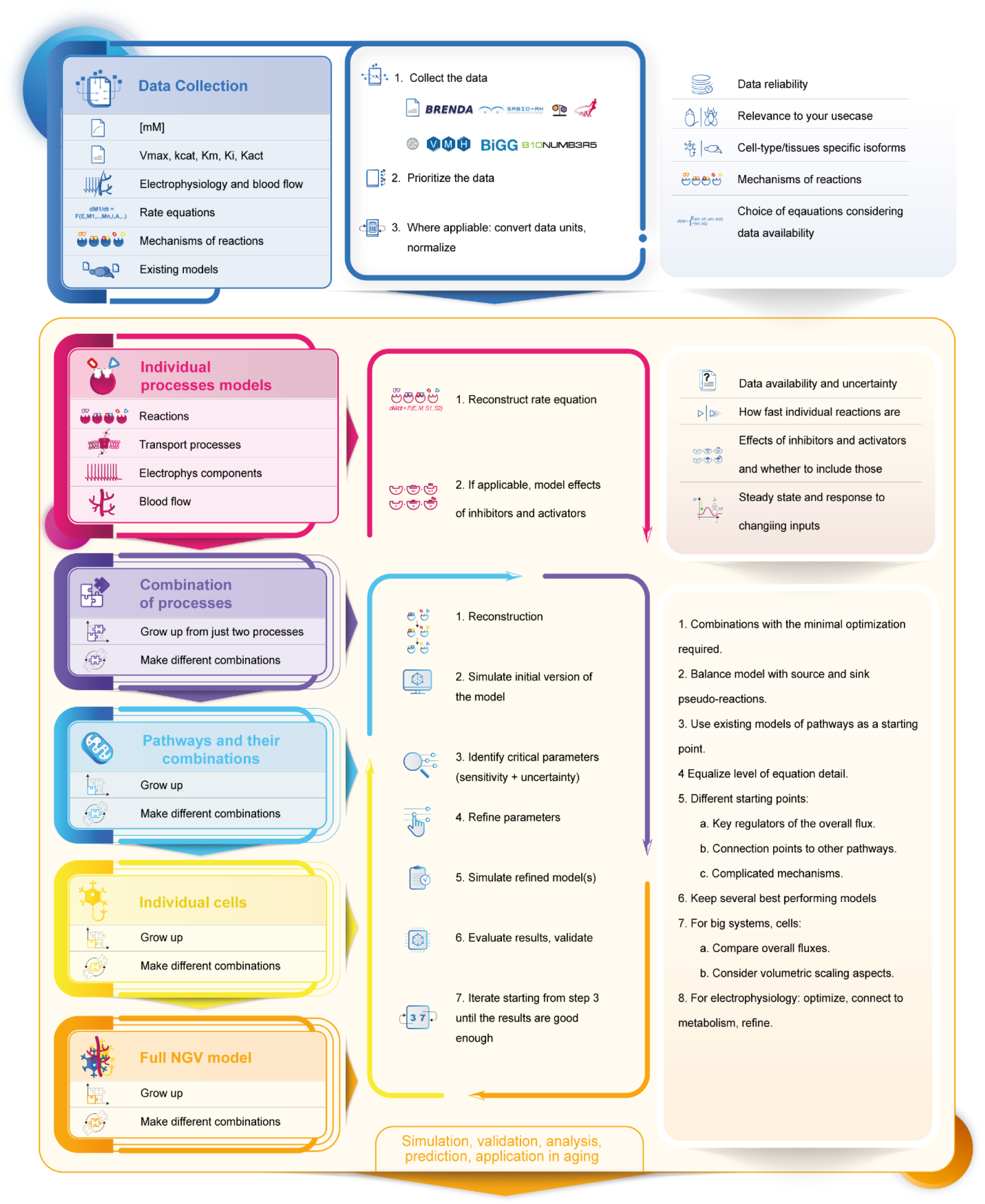
Bottom-up iterative model building workflow and the key considerations.

**Supplementary Fig. 18.**
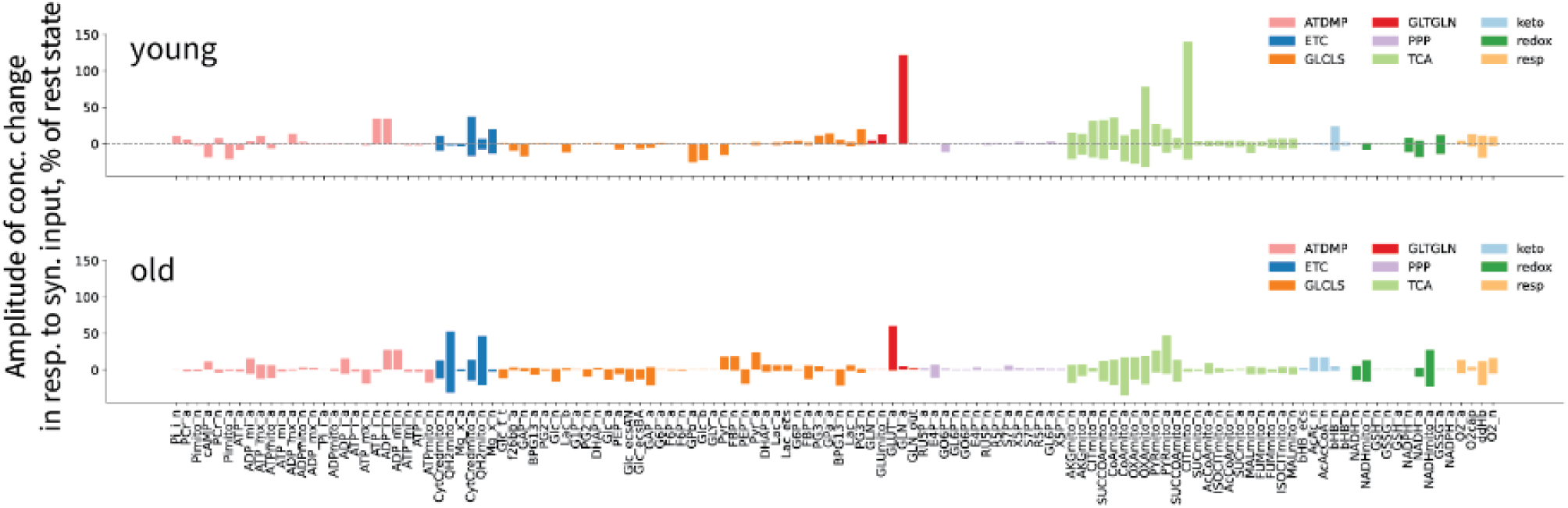
Labels of individual metabolites for Fig. 3d.

**Supplementary Fig. 19.**
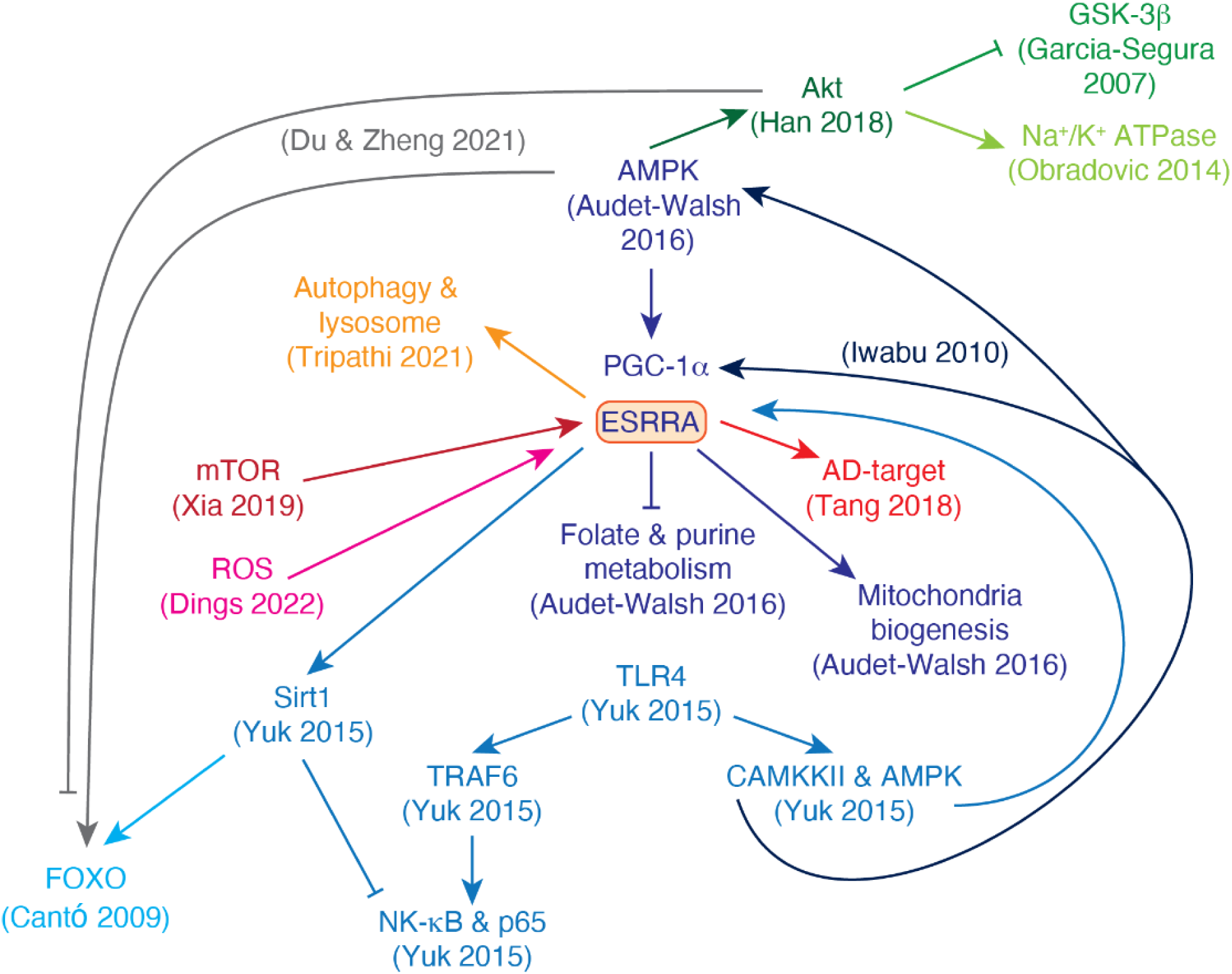
Literature evidence for ESRRA being a regulatory hub of aging-associated pathways (colored by reference).

### Supplementary Information File 1

#### Explanation of the Fruchterman-Reingold force-directed algorithm to position nodes

The Fruchterman-Reingold force-directed algorithm is used for a representation of the network. Edges correspond to springs which are holding nodes close, and nodes correspond to repelling objects. There are two types of forces which define the position of nodes as described below.

1. Attractive force, which acts between nodes that are connected by an edge. It is stronger for nodes connected by higher-weight edges, so these nodes are pulled closer together by the attractive force.
2. Repulsive force, which acts between all pairs of nodes (regardless of connecting edge presence). This force pushes nodes away from each other.

So the resulting edge lengths are not directly proportional to the weights. Instead, the edge weight affects the strength of the attractive force between connected nodes, where larger weight of edge (which in our case corresponds to smaller metabolic adaptability) means a stronger attractive force, and nodes being pulled together.

#### Centrality in context of metabolic adaptability

Centrality, or closeness centrality (CC), is calculated as a reciprocal of the sum of shortest path distances between the node and all other nodes. Higher centrality means smaller metabolic adaptability: CC is reciprocal to distance, while distance is reciprocal to weight, which makes CC proportional to the weight, while weight is defined as reciprocal of metabolic adaptability, which makes CC reciprocal of metabolic adaptability (Equation 1).

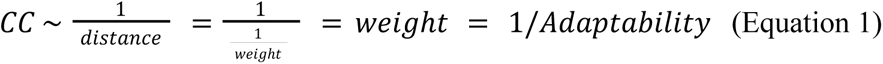

### Supplementary Information File 2

**Changes in other characteristics of neuronal firing (related to** **Fig.3**):

AHP1_depth_from_peak: -7.42%, old: 144.86, young: 156.48
AHP2_depth_from_peak: -7.23%, old: 140.48, young: 151.42
AP1_amp: -8.5%, old: 114.94, young: 125.62
AP1_begin_voltage: -0.35%, old: -51.64, young: -51.82
AP1_begin_width: 0.0%, old: 1.3, young: 1.3
AP1_peak: -14.23%, old: 63.3, young: 73.81
AP1_width: 3.59%, old: 0.76, young: 0.73
AP2_AP1_begin_width_diff: nan%, old: 0.0, young: 0.0
AP2_AP1_diff: -31.37%, old: -1.04, young: -1.52
AP2_AP1_peak_diff: -53.87%, old: -0.49, young: -1.05
AP2_amp: -8.22%, old: 113.9, young: 124.1
AP2_begin_voltage: -0.53%, old: -51.08, young: -51.35
AP2_begin_width: 0.0%, old: 1.3, young: 1.3
AP2_peak: -13.66%, old: 62.82, young: 72.75
AP2_width: 3.46%, old: 0.72, young: 0.7
APlast_amp: -6.17%, old: 105.99, young: 112.96
APlast_width: 3.5%, old: 0.79, young: 0.76
ISI_CV: -2.13%, old: 0.57, young: 0.58
ISI_log_slope: -0.57%, old: 0.16, young: 0.17
ISI_log_slope_skip: 0.24%, old: 0.22, young: 0.22
ISI_semilog_slope: 6.44%, old: 0.02, young: 0.02
Spikecount: -8.47%, old: 54, young: 59
Spikecount_stimint: -8.47%, old: 54, young: 59
adaptation_index: 11.23%, old: 0.01, young: 0.01
adaptation_index2: 11.31%, old: 0.01, young: 0.01
amp_drop_first_last: -29.32%, old: 8.64, young: 12.23
amp_drop_first_second: -53.87%, old: 0.49, young: 1.05
amp_drop_second_last: -27.0%, old: 8.16, young: 11.17
decay_time_constant_after_stim: -91.66%, old: 1603.71, young: 19220.41
doublet_ISI: 5.9%, old: 195.7, young: 184.8
inv_fifth_ISI: -6.26%, old: 8.94, young: 9.53
inv_first_ISI: -5.57%, old: 5.11, young: 5.41
inv_fourth_ISI: -6.06%, old: 8.31, young: 8.84
inv_last_ISI: -7.22%, old: 2.09, young: 2.26
inv_second_ISI: -5.68%, old: 6.53, young: 6.92
inv_third_ISI: -5.8%, old: 7.53, young: 7.99
inv_time_to_first_spike: -33.47%, old: 6.0, young: 9.02
irregularity_index: 16.38%, old: 8.87, young: 7.62
max_amp_difference: -16.07%, old: 1.14, young: 1.35
maximum_voltage: -14.23%, old: 63.3, young: 73.81
maximum_voltage_from_voltagebase: -2.75%, old: 141.84, young: 145.86
mean_AP_amplitude: -6.7%, old: 108.94, young: 116.76
mean_frequency: -8.46%, old: 7.55, young: 8.25
minimum_voltage: 3.33%, old: -85.43, young: -82.68
number_initial_spikes: -10.0%, old: 18, young: 20
steady_state_hyper: 6.37%, old: -85.14, young: -80.05
steady_state_voltage: 8.57%, old: -83.27, young: -76.7
time_to_first_spike: 50.32%, old: 166.7, young: 110.9
time_to_last_spike: -0.02%, old: 7150.2, young: 7151.4
time_to_second_spike: 22.56%, old: 362.4, young: 295.7
voltage_base: 9.0%, old: -78.54, young: -72.05

**Statistical tests for comparison of characteristics of neuronal firing (Figure 3).**

AP_amplitude

Sample sizes: 59,54

Levene statistic: 6.605499030481687 p-value: 0.011491236053743635
Levene: different variances (reject H0)
Fligner statistic: 6.630869011027655 p-value: 0.01002263560239055
Fligner: different variances (reject H0)
Wilcoxon-Mann-Whitney two-sided U test statistic: 3057.0 p-value: 4.0249411168018834e-17
Wilcoxon-Mann-Whitney two-sided U test: different distributions (reject H0)
Kolmogorov-Smirnov two-sided test statistic: 0.834902699309479 p-value: 6.661338147750939e-16
Kolmogorov-Smirnov two-sided test: different distributions (reject H0)

AP_amplitude_change

Sample sizes: 58,53

Levene statistic: 3.8760940845777174 p-value: 0.051516931692045034
Levene: same variances (fail to reject H0)
Fligner statistic: 4.545621778929398 p-value: 0.033003033701898096
Fligner: different variances (reject H0)
Wilcoxon-Mann-Whitney two-sided U test statistic: 920.0 p-value: 0.0002729809833335603
Wilcoxon-Mann-Whitney two-sided U test: different distributions (reject H0)
Kolmogorov-Smirnov two-sided test statistic: 0.42940793754066364 p-value: 3.8705542437011964e-05
Kolmogorov-Smirnov two-sided test: different distributions (reject H0)

AP_amplitude_diff

Sample sizes: 58,53

Levene statistic: 6.152257861376695 p-value: 0.014653671953606095
Levene: different variances (reject H0)
Fligner statistic: 5.75689520557301 p-value: 0.016424069262576897
Fligner: different variances (reject H0)
Wilcoxon-Mann-Whitney two-sided U test statistic: 1491.0 p-value: 0.7882208195357832
Wilcoxon-Mann-Whitney two-sided U test: same distribution (fail to reject H0)
Kolmogorov-Smirnov two-sided test statistic: 0.1486662329212752 p-value: 0.5084641118074855
Kolmogorov-Smirnov two-sided test: same distribution (fail to reject H0)

AP_amplitude_from_voltagebase

Sample sizes: 59,54

Levene statistic: 7.459710428818111 p-value: 0.007341447281952335
Levene: different variances (reject H0)
Fligner statistic: 7.391190337972734 p-value: 0.006554409829916771
Fligner: different variances (reject H0)
Wilcoxon-Mann-Whitney two-sided U test statistic: 1862.0 p-value: 0.12275003593521057
Wilcoxon-Mann-Whitney two-sided U test: same distribution (fail to reject H0)
Kolmogorov-Smirnov two-sided test statistic: 0.1864406779661017 p-value: 0.23771345375057895
Kolmogorov-Smirnov two-sided test: same distribution (fail to reject H0)

AP_begin_voltage

Sample sizes: 59,54

Levene statistic: 0.3716334196659708 p-value: 0.5433611341543954
Levene: same variances (fail to reject H0)
Fligner statistic: 0.3123751596228548 p-value: 0.5762263366963747
Fligner: same variances (fail to reject H0)
Wilcoxon-Mann-Whitney two-sided U test statistic: 1226.0 p-value: 0.03514923245426359
Wilcoxon-Mann-Whitney two-sided U test: different distributions (reject H0)
Kolmogorov-Smirnov two-sided test statistic: 0.21876961707470183 p-value: 0.1096654688128944
Kolmogorov-Smirnov two-sided test: same distribution (fail to reject H0)

AP_height

Sample sizes: 59,54

Levene statistic: 7.459710428818096 p-value: 0.007341447281952389
Levene: different variances (reject H0)
Fligner statistic: 7.391190337972734 p-value: 0.006554409829916771
Fligner: different variances (reject H0)
Wilcoxon-Mann-Whitney two-sided U test statistic: 3079.0 p-value: 1.3582777666804203e-17
Wilcoxon-Mann-Whitney two-sided U test: different distributions (reject H0)
Kolmogorov-Smirnov two-sided test statistic: 0.8549905838041432 p-value: 6.661338147750939e-16
Kolmogorov-Smirnov two-sided test: different distributions (reject H0)

peak_voltage

Sample sizes: 59,54

Levene statistic: 7.459710428818096 p-value: 0.007341447281952389
Levene: different variances (reject H0)
Fligner statistic: 7.391190337972734 p-value: 0.006554409829916771
Fligner: different variances (reject H0)
Wilcoxon-Mann-Whitney two-sided U test statistic: 3079.0 p-value: 1.3582777666804203e-17
Wilcoxon-Mann-Whitney two-sided U test: different distributions (reject H0)
Kolmogorov-Smirnov two-sided test statistic: 0.8549905838041432 p-value: 6.661338147750939e-16
Kolmogorov-Smirnov two-sided test: different distributions (reject H0)

AP_rise_rate

Sample sizes: 59,54

Levene statistic: 3.2120282887999094 p-value: 0.07582303196261797
Levene: same variances (fail to reject H0)
Fligner statistic: 8.500912633404996 p-value: 0.003549683927627355
Fligner: different variances (reject H0)
Wilcoxon-Mann-Whitney two-sided U test statistic: 3008.0 p-value: 4.274566678078413e-16
Wilcoxon-Mann-Whitney two-sided U test: different distributions (reject H0)
Kolmogorov-Smirnov two-sided test statistic: 0.8163841807909604 p-value: 6.661338147750939e-16
Kolmogorov-Smirnov two-sided test: different distributions (reject H0)

AP_fall_rate

Sample sizes: 59,54

Levene statistic: 3.066184321429485 p-value: 0.08269937892109969
Levene: same variances (fail to reject H0)
Fligner statistic: 6.71176089244189 p-value: 0.0095779100160061
Fligner: different variances (reject H0)
Wilcoxon-Mann-Whitney two-sided U test statistic: 442.0 p-value: 3.7647624938176844e-11
Wilcoxon-Mann-Whitney two-sided U test: different distributions (reject H0)
Kolmogorov-Smirnov two-sided test statistic: 0.7052730696798494 p-value: 3.952393967665557e-14
Kolmogorov-Smirnov two-sided test: different distributions (reject H0)

AP_fall_rate_change

Sample sizes: 58,53

Levene statistic: 0.6190034404678872 p-value: 0.433125124634122
Levene: same variances (fail to reject H0)
Fligner statistic: 3.187363730600531 p-value: 0.0742095978425819
Fligner: same variances (fail to reject H0)
Wilcoxon-Mann-Whitney two-sided U test statistic: 351.0 p-value: 2.5793688476458548e-12
Wilcoxon-Mann-Whitney two-sided U test: different distributions (reject H0)
Kolmogorov-Smirnov two-sided test statistic: 0.8103448275862069 p-value: 1.775156606426841e-20
Kolmogorov-Smirnov two-sided test: different distributions (reject H0)

AP_peak_upstroke

Sample sizes: 59,54

Levene statistic: 4.408526533521235 p-value: 0.038025854014361024
Levene: different variances (reject H0)
Fligner statistic: 4.601321053363534 p-value: 0.03194732985273627
Fligner: different variances (reject H0)
Wilcoxon-Mann-Whitney two-sided U test statistic: 3064.0 p-value: 2.853607187168821e-17
Wilcoxon-Mann-Whitney two-sided U test: different distributions (reject H0)
Kolmogorov-Smirnov two-sided test statistic: 0.7919020715630886 p-value: 6.661338147750939e-16
Kolmogorov-Smirnov two-sided test: different distributions (reject H0)

AP_peak_downstroke

Sample sizes: 59,54

Levene statistic: 1.2715834062751352 p-value: 0.2619014544184533
Levene: same variances (fail to reject H0)
Fligner statistic: 1.6194972442709932 p-value: 0.2031619039342202
Fligner: same variances (fail to reject H0)
Wilcoxon-Mann-Whitney two-sided U test statistic: 239.0 p-value: 7.258645224012553e-15
Wilcoxon-Mann-Whitney two-sided U test: different distributions (reject H0)
Kolmogorov-Smirnov two-sided test statistic: 0.834902699309479 p-value: 6.661338147750939e-16
Kolmogorov-Smirnov two-sided test: different distributions (reject H0)

spike_half_width

Sample sizes: 59,54

Levene statistic: 0.8356847163178224 p-value: 0.36261535212273177
Levene: same variances (fail to reject H0)
Fligner statistic: 0.9340883843599507 p-value: 0.3338028095263355
Fligner: same variances (fail to reject H0)
Wilcoxon-Mann-Whitney two-sided U test statistic: 716.0 p-value: 4.7016301099113494e-07
Wilcoxon-Mann-Whitney two-sided U test: different distributions (reject H0)
Kolmogorov-Smirnov two-sided test statistic: 0.5266792215944758 p-value: 1.0318489573890588e-07
Kolmogorov-Smirnov two-sided test: different distributions (reject H0)

spike_width2

Sample sizes: 58,53

Levene statistic: 0.4059195398968852 p-value: 0.5253838004179687
Levene: same variances (fail to reject H0)
Fligner statistic: 0.8000585301188278 p-value: 0.3710758705566992
Fligner: same variances (fail to reject H0)
Wilcoxon-Mann-Whitney two-sided U test statistic: 728.0 p-value: 1.8132084872745687e-06
Wilcoxon-Mann-Whitney two-sided U test: different distributions (reject H0)
Kolmogorov-Smirnov two-sided test statistic: 0.4362394274560833 p-value: 2.7784173407874313e-05
Kolmogorov-Smirnov two-sided test: different distributions (reject H0)

depolarized_base

Sample sizes: 58,53

Levene statistic: 0.21330792406056287 p-value: 0.6451074156436485
Levene: same variances (fail to reject H0)
Fligner statistic: 0.5099912340048506 p-value: 0.47514265083983753
Fligner: same variances (fail to reject H0)
Wilcoxon-Mann-Whitney two-sided U test statistic: 1622.0 p-value: 0.617871778194361
Wilcoxon-Mann-Whitney two-sided U test: same distribution (fail to reject H0)
Kolmogorov-Smirnov two-sided test statistic: 0.1724137931034483 p-value: 0.3298788555460255
Kolmogorov-Smirnov two-sided test: same distribution (fail to reject H0)

AHP_depth

Sample sizes: 59,54

Levene statistic: 0.08022401294287185 p-value: 0.7775216314033404
Levene: same variances (fail to reject H0)
Fligner statistic: 0.10770161746045517 p-value: 0.7427761389915951
Fligner: same variances (fail to reject H0)
Wilcoxon-Mann-Whitney two-sided U test statistic: 271.0 p-value: 3.055267399749676e-14
Wilcoxon-Mann-Whitney two-sided U test: different distributions (reject H0)
Kolmogorov-Smirnov two-sided test statistic: 0.8518518518518519 p-value: 6.661338147750939e-16
Kolmogorov-Smirnov two-sided test: different distributions (reject H0)

AHP_depth_abs

Sample sizes: 59,54

Levene statistic: 0.08022401294287124 p-value: 0.7775216314033404
Levene: same variances (fail to reject H0)
Fligner statistic: 0.10770132317169079 p-value: 0.7427764779807888
Fligner: same variances (fail to reject H0)
Wilcoxon-Mann-Whitney two-sided U test statistic: 1510.0 p-value: 0.6353512902776368
Wilcoxon-Mann-Whitney two-sided U test: same distribution (fail to reject H0)
Kolmogorov-Smirnov two-sided test statistic: 0.12962962962962962 p-value: 0.6655913571840438
Kolmogorov-Smirnov two-sided test: same distribution (fail to reject H0)

AHP_depth_from_peak

Sample sizes: 59,54

Levene statistic: 1.3399447901571637 p-value: 0.24952849723117262
Levene: same variances (fail to reject H0)
Fligner statistic: 2.424339978036405 p-value: 0.1194635491567324
Fligner: same variances (fail to reject H0)
Wilcoxon-Mann-Whitney two-sided U test statistic: 2893.0 p-value: 8.049943715324859e-14
Wilcoxon-Mann-Whitney two-sided U test: different distributions (reject H0)
Kolmogorov-Smirnov two-sided test statistic: 0.7777777777777778 p-value: 6.661338147750939e-16
Kolmogorov-Smirnov two-sided test: different distributions (reject H0)

AHP_slow_time

Sample sizes: 57,52

Levene statistic: 0.2773733483050493 p-value: 0.5995181094433077
Levene: same variances (fail to reject H0)
Fligner statistic: 0.3607862611307999 p-value: 0.5480698901731853
Fligner: same variances (fail to reject H0)
Wilcoxon-Mann-Whitney two-sided U test statistic: 1598.0 p-value: 0.48348010610693026
Wilcoxon-Mann-Whitney two-sided U test: same distribution (fail to reject H0)
Kolmogorov-Smirnov two-sided test statistic: 0.22604588394062078 p-value: 0.10176313080703958
Kolmogorov-Smirnov two-sided test: same distribution (fail to reject H0)

ISI_values

Sample sizes: 57,52

Levene statistic: 0.02132078025677348 p-value: 0.8841831976486665
Levene: same variances (fail to reject H0)
Fligner statistic: 0.07623370008066287 p-value: 0.7824677972955265
Fligner: same variances (fail to reject H0)
Wilcoxon-Mann-Whitney two-sided U test statistic: 1117.0 p-value: 0.027011518317369983
Wilcoxon-Mann-Whitney two-sided U test: different distributions (reject H0)
Kolmogorov-Smirnov two-sided test statistic: 0.3684210526315789 p-value: 0.0007881130606729458
Kolmogorov-Smirnov two-sided test: different distributions (reject H0)

### Supplementary Information File 3

The top-scoring TF was ESRRA (estrogen-related receptor alpha). This TF regulates expression of multiple metabolism-related genes, including those of mitochondrial function, biogenesis and turnover, as well as lipid catabolism (Tripathi et al., 2020). It is also linked to autophagy and NF-kB inflammatory response via Sirt1 signaling (Cantó et al., 2009; Yuk et al., 2015; Kim et al., 2018; Suresh et al., 2018). Mitochondrial dysfunction and autophagy impairments are consistently among the hallmarks of aging (López-Otín et al., 2013, 2023; Mattson and Arumugam, 2018). Notably, ESRRA expression is downregulated in aging according to various studies (Schaum et al., 2020; Tripathi et al., 2020).

The second-scoring TF was Nkx2-5 (NK2 homeobox 5), which is highly conserved among species and mostly studied in development and cardiac function (Takeda et al., 2009). Reduction of Nkx2-5 cardiac expression has been reported in aging (Volkova et al., 2005).

The third-ranked TF was the evolutionary conserved energy sensor NFE2L1 (nuclear factor erythroid 2-related factor 1, also called Nrf1 or nuclear respiratory factor 1). It is one of the key regulators of redox signaling and homeostasis. Dysfunction of this TF is associated with glucose metabolism reprogramming via AMPK signaling (Yang et al., 2021). NFE2L1 also upregulates expression of proteasomal genes in an ERK-signaling dependent manner (Zhang et al., 2021b), which is suggested to contribute to the development of neurodegenerative diseases (Lee et al., 2011).

The next TF was ZBED1 (zinc finger BED domain-containing protein 1). It acts as a small ubiquitin-like modifier (SUMO) ligase by SUMOylating Mi2-alpha during nucleosome remodeling and deacetylation (Yamashita et al., 2016).

The fifth TF, THAP4 (nitrobindin), detoxifies reactive nitrogen and oxygen species and scavenges peroxynitrite (De Simone et al., 2018).

The next TF was PREB (prolactin regulatory element binding). Interestingly, it has been reported as one of the links between aging and Alzheimer’s disease (Zhou et al., 2019).

The next TF was HIF1A (hypoxia inducible factor 1), which serves as a key regulator of the hypoxia response at both cellular and system scale. The role of HIF1a in brain aging and neurodegenerative diseases is convoluted, multifaceted and incompletely understood. On one hand, HIF1a promotes erythropoiesis, angiogenesis and exerts neuroprotection (Majmundar et al., 2010; Burtscher et al., 2021). On the other hand, there are contradictory reports on either the detrimental (Sun et al., 2006; March-Diaz et al., 2021; Lee et al., 2023) or protective (Ashok et al., 2017) role of HIF1a in neurodegeneration, Alzheimer’s disease in particular. To add the complexity, HIF1a is involved in inflammatory response and metabolism regulation (McGettrick and O’Neill, 2020; Taylor and Scholz, 2022). Even though mechanisms of HIF1a are so multifaceted, therapeutic potential of this TF has been recognized (Lee et al., 2019; Luo et al., 2022).

The next TF was MYRFL (myelin regulatory factor-like protein). Its biological function (and that of some other myelin regulatory factors, such as MYRF) is poorly understood, which is surprising given the importance of myelin for neuronal health (Huang et al., 2021).

The next TF was ZNF878 (zinc finger protein 878), which is one of the hundreds in the rapidly evolved KRAB-domain containing family (Shen et al., 2021).

The last out of top-10 TFs was SALL3 (spalt-like transcription factor 3). It modulates DNA methyltransferase activity and influences human induced pluripotent stem cell differentiation (Kuroda et al., 2019).

We further searched the STRING database (Szklarczyk et al., 2019) for the protein-protein associations of the top TF ESRRA (Fig. 5c), following which we performed a literature search for the top-10 proteins from this search: Hif1a, Sirt1, Hdac8, Ppargc1a, Ppargc1b, Mef2c, Nrip1, Ncoa1, Tfam, Perm1. Interestingly, numerous reports attribute roles in aging and neurodegeneration to these proteins as detailed below.

Hif1a is a common hit in the top-10 of STRING associations and ChEA3 enrichment with its implications in brain aging and neurodegeneration described above.

Sirt1 is a NAD+-dependent deacetylase, a member of the sirtuin family of proteins. It is largely studied for its role in aging, longevity, apoptosis, stress resistance, inflammation, linking nutrition and chromatin regulation, energy homeostasis and caloric restriction (Rodgers et al., 2005; Guarente, 2006; Milne et al., 2007; Cantó et al., 2009; Finkel et al., 2009; Bishop et al., 2010; Ledford, 2010; Gut and Verdin, 2013; O’Neill and Hardie, 2013; Ng et al., 2015; Shin et al., 2016; Satoh et al., 2017; Katsyuba et al., 2018; Yang et al., 2018).

Another protein found in STRING associations of ESRRA is HDAC8 (histone deacetylase 8), closely related to the sirtuins signaling. It is largely recognised as a promising drug target in several disorders (Chakrabarti et al., 2015; Mormino et al., 2021; Zhao et al., 2021; Emmons et al., 2022).

Another protein in our list, Ppargc1a (peroxisome proliferator-activated receptor gamma coactivator 1-alpha, also called PGC-1alpha), is an important regulator of energy metabolism and is implicated in aging (Rodgers et al., 2005; Anderson and Prolla, 2009; Garcia et al., 2018). It is associated with Parkinson’s (Li et al., 2022) and Huntington’s diseases (Cui et al., 2006).

Closely related to PGC-1alpha is the other protein on our list, Ppargc1b (peroxisome proliferator-activated receptor gamma coactivator 1-alpha, also called PGC-1beta), which is also involved in energy metabolism regulation, but less well studied and is an active area of research (Thiepold et al., 2017; Thibonnier et al., 2020).

The next protein on our list is Tfam (mitochondrial transcription factor 1), which is regulated by PGC-1alpha and is implicated in brain aging and neurodegeneration (Grimm and Eckert, 2017; Kang et al., 2018).

Another protein on the list is Perm1 (PGC-1 And ERR-Induced Regulator In Muscle Protein 1) is mostly studied in cardiac mitochondrial metabolism regulation (Huang et al., 2022).

The next on the list is Nrip1 (nuclear receptor interacting protein 1). It is an oxidative metabolism regulator and a potential therapeutic target in Down syndrome (Izzo et al., 2014).

The other protein on the list is Ncoa1 (nuclear receptor coactivator 1), which is involved in hormonal regulation, learning, memory and neurogenesis (Nishihara et al., 2007; Sun and Xu, 2020).

Next, brain-expressed Mef2c (myocyte enhancer factor 2C) is a TF downregulated in aging in an interferon signaling-dependent way (Deczkowska et al., 2017). Furthermore, therapeutic potential of this TF in neurodegeneration and aging has been recently shown with its effects in promoting cognitive resilience (Barker et al., 2021).

To sum up, our identified anti-aging targets largely align with the literature data on therapeutics for healthy aging (Campisi et al., 2019), but also suggest a role of a few poorly studied TFs in the brain energy metabolism aging and provide the insights on the links between molecular mechanisms implicated in aging and neurodegeneration.

### Supplementary Information File 4

This information will be available after peer-reviewed publication.

### Supplementary Information File 5

This information will be available after peer-reviewed publication.

### Supplementary Information File 6

This information will be available after peer-reviewed publication.

### Supplementary Information File 7

Mapping of model variables indexes to descriptive names and Bigg (King et al., 2016) nomenclature (where available).

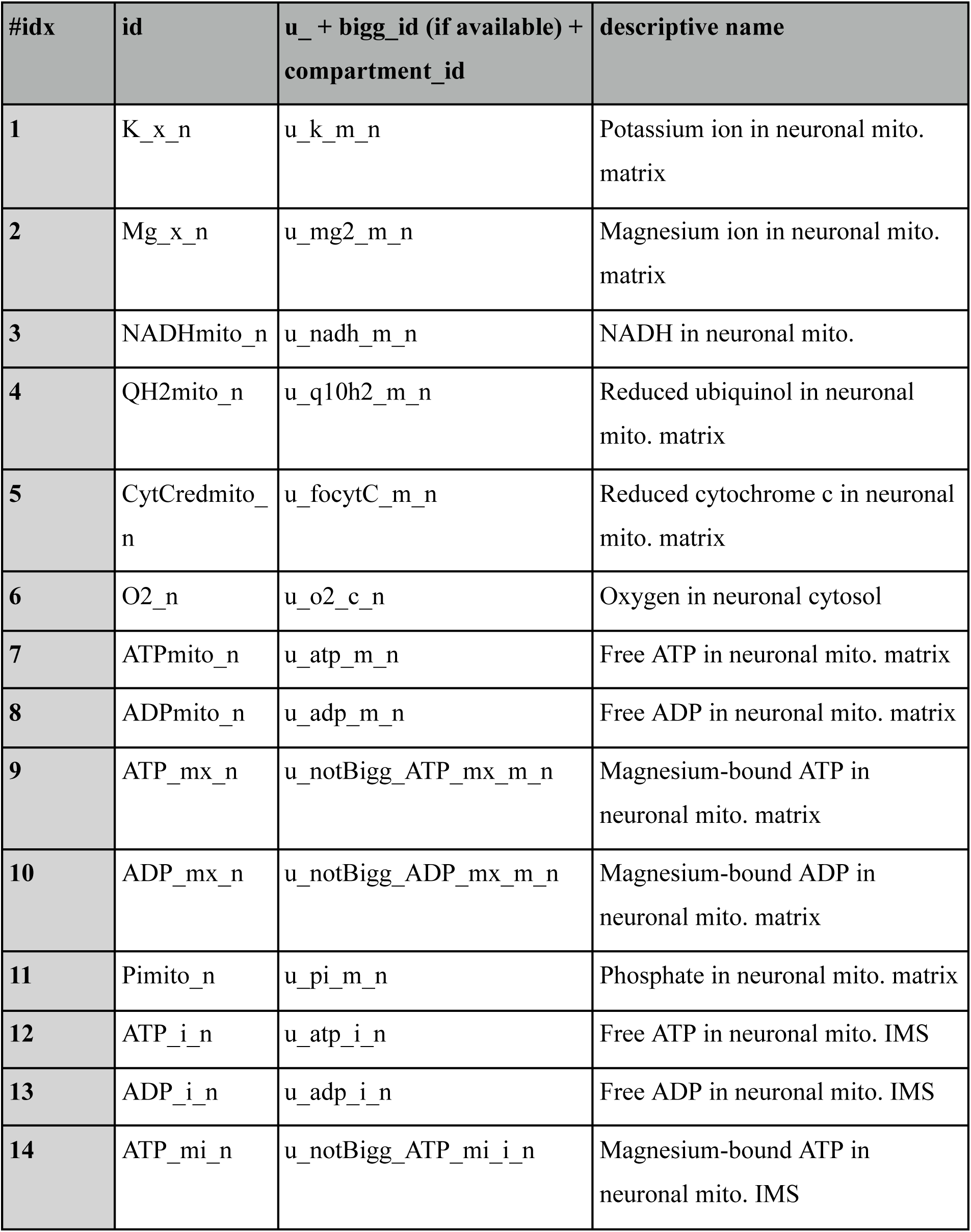
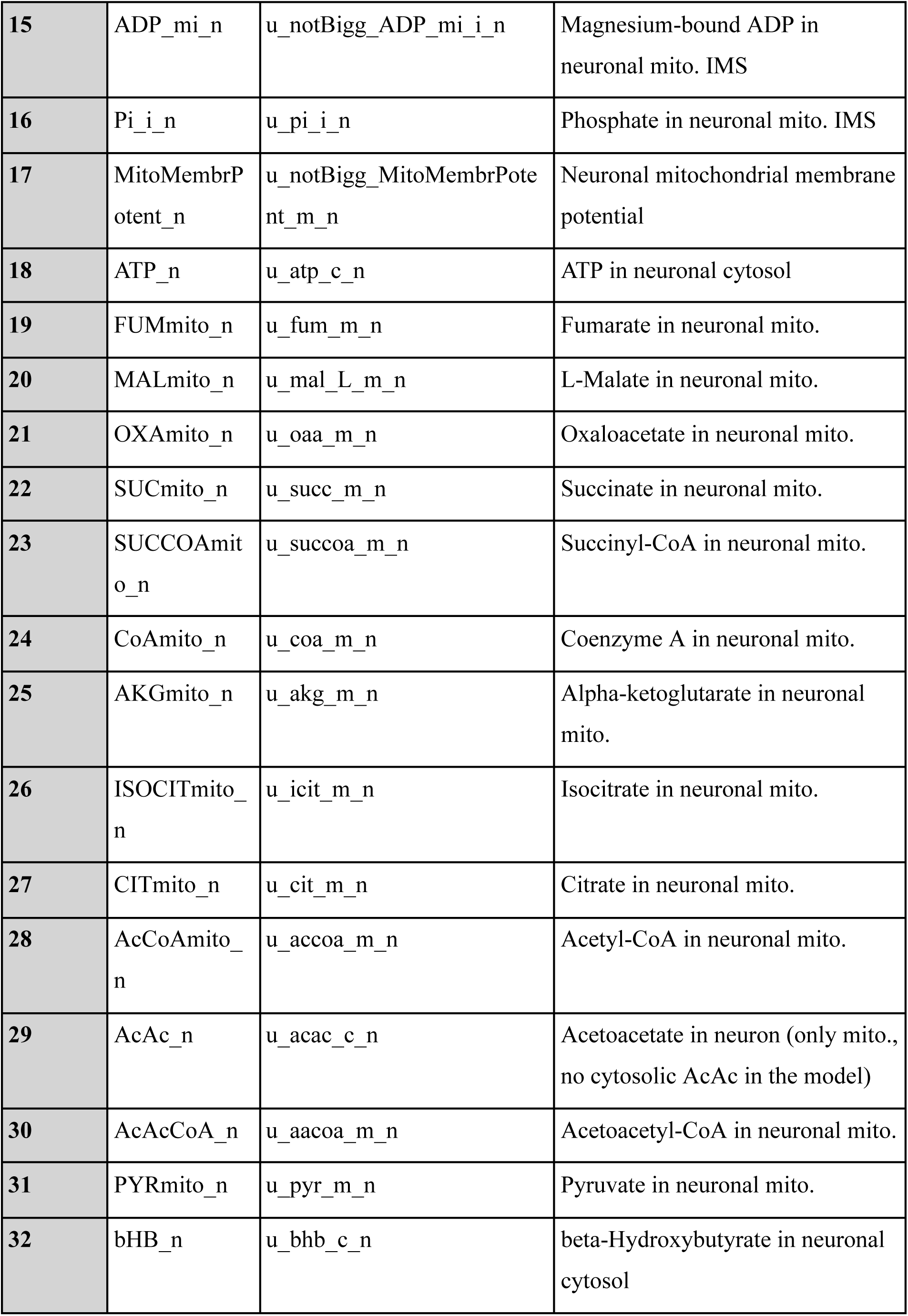
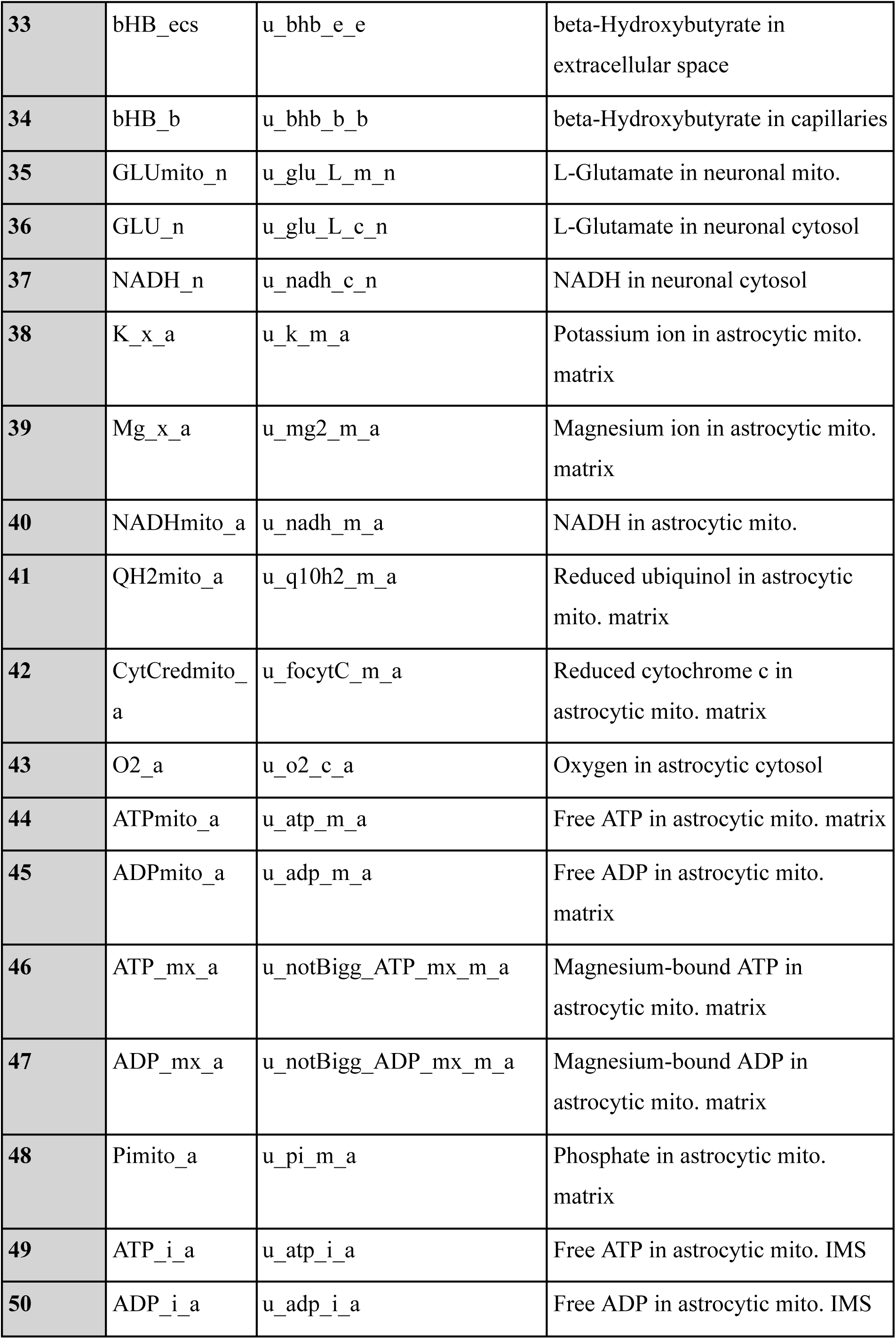
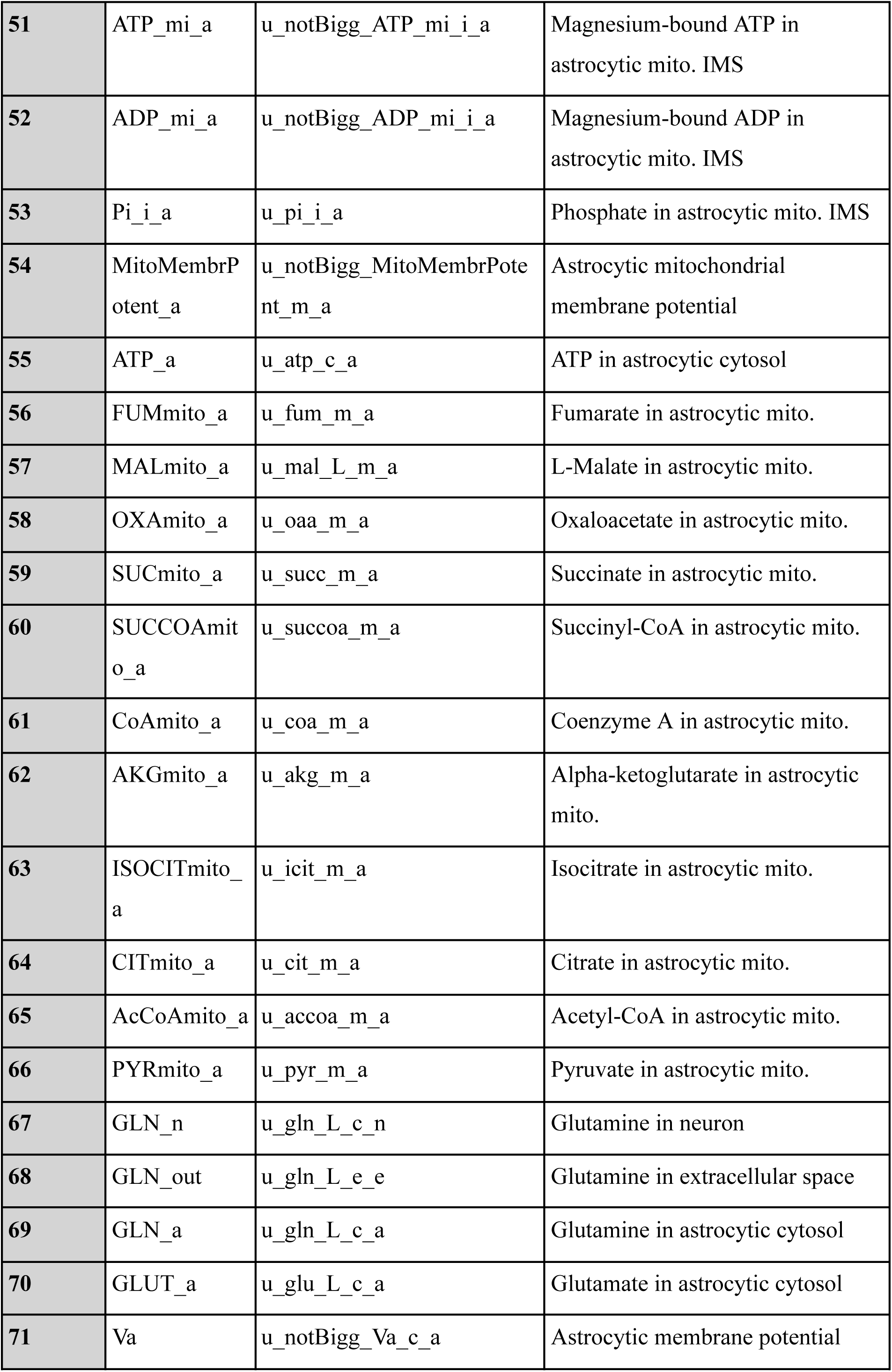
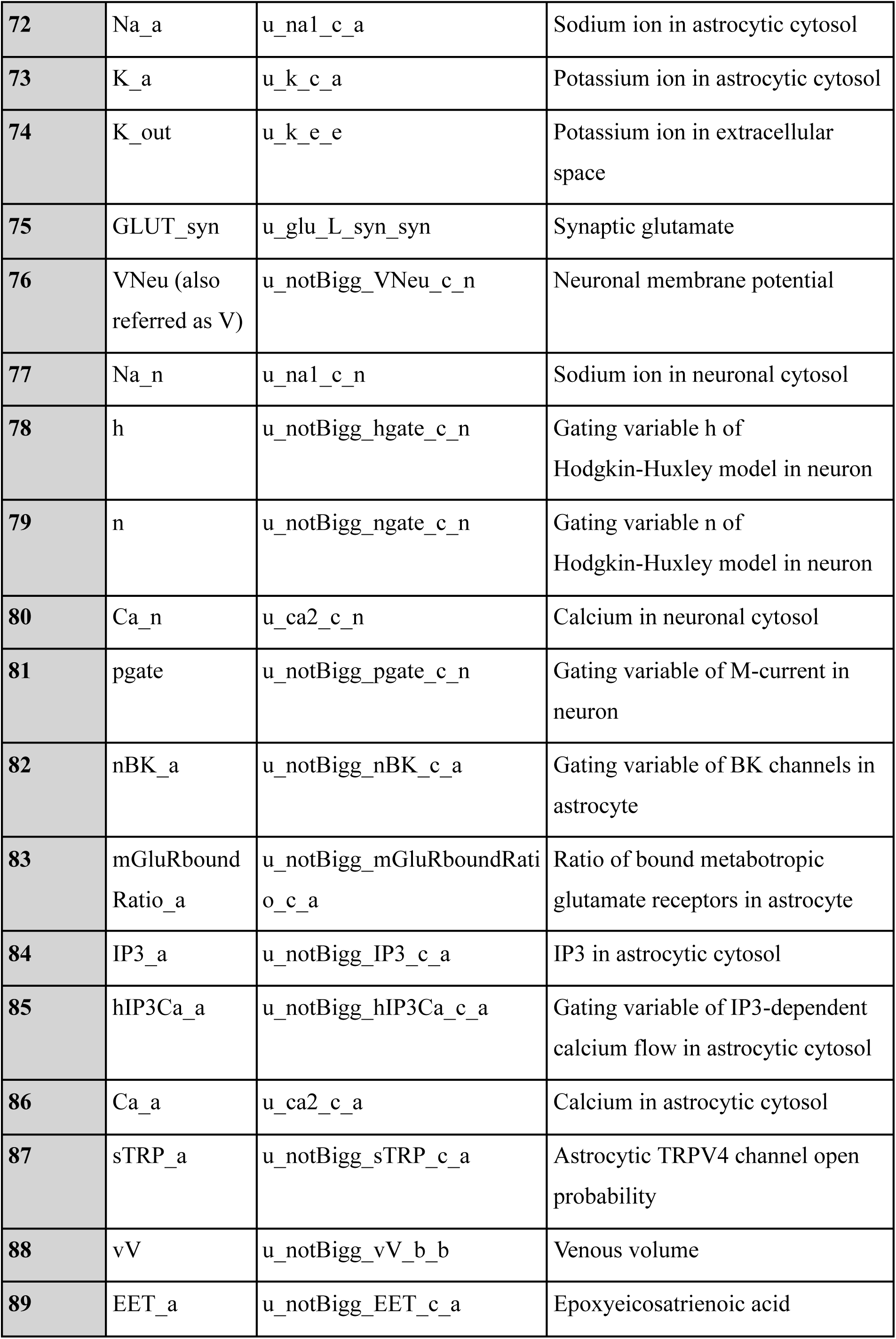
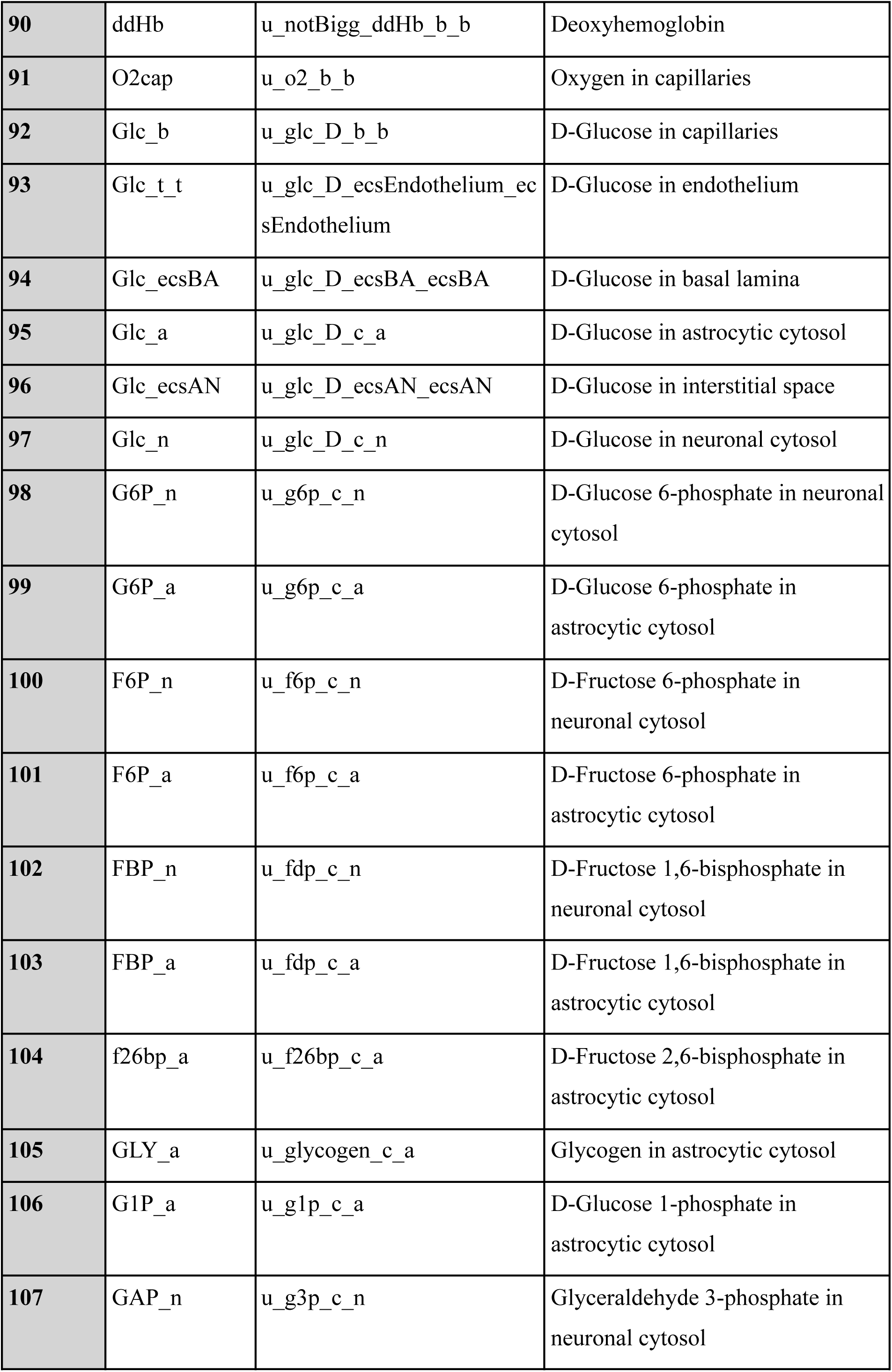
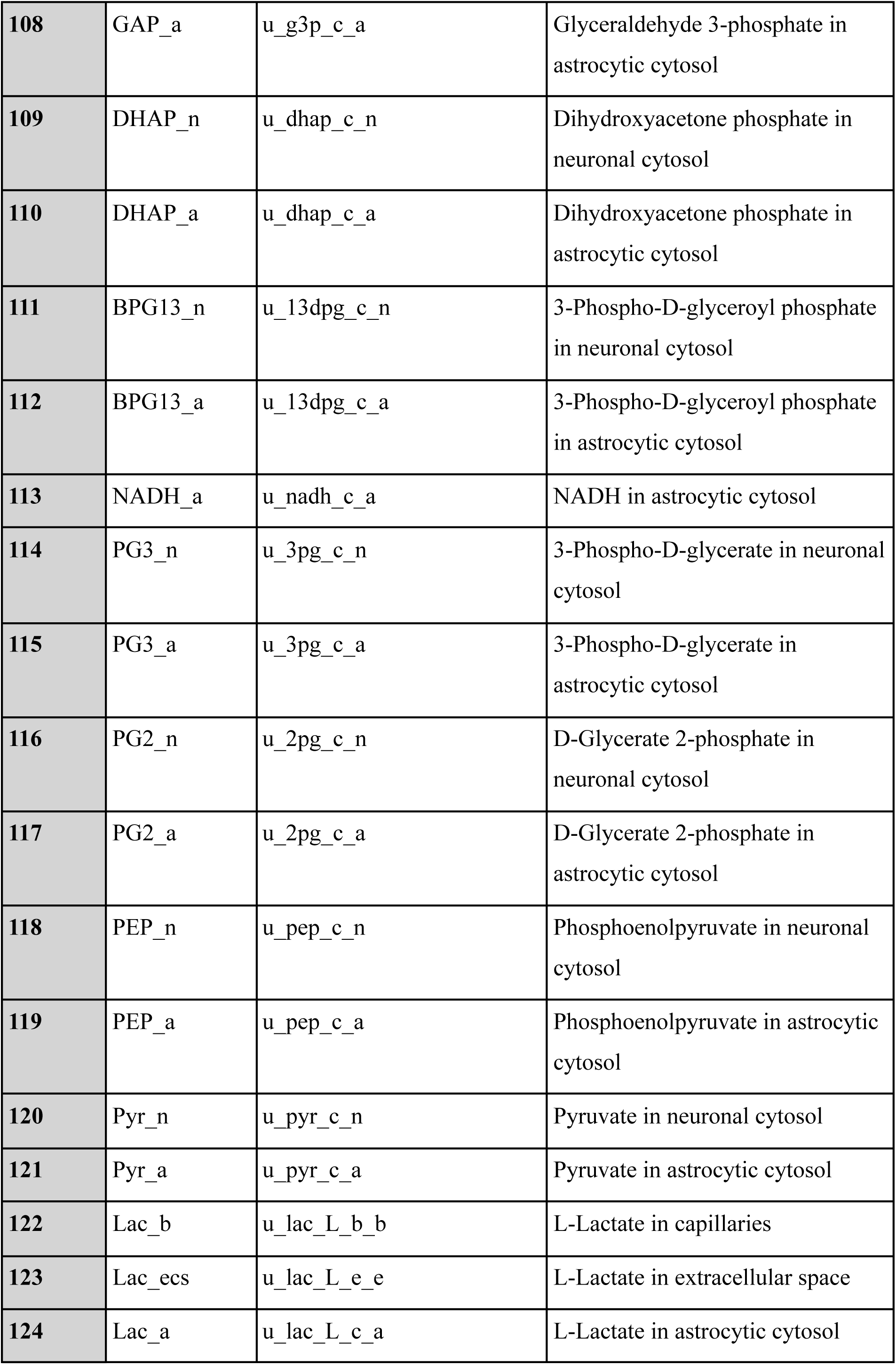
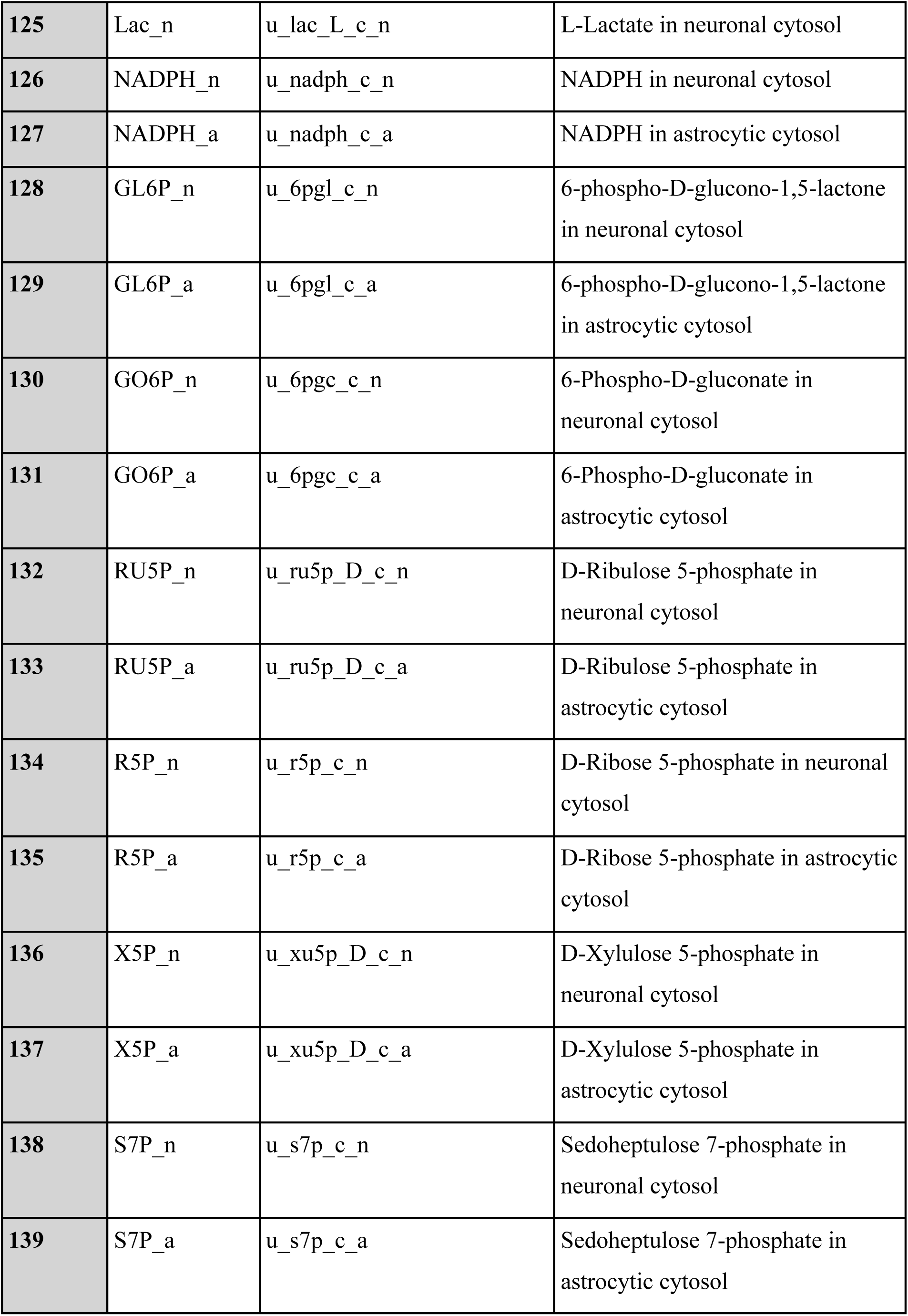
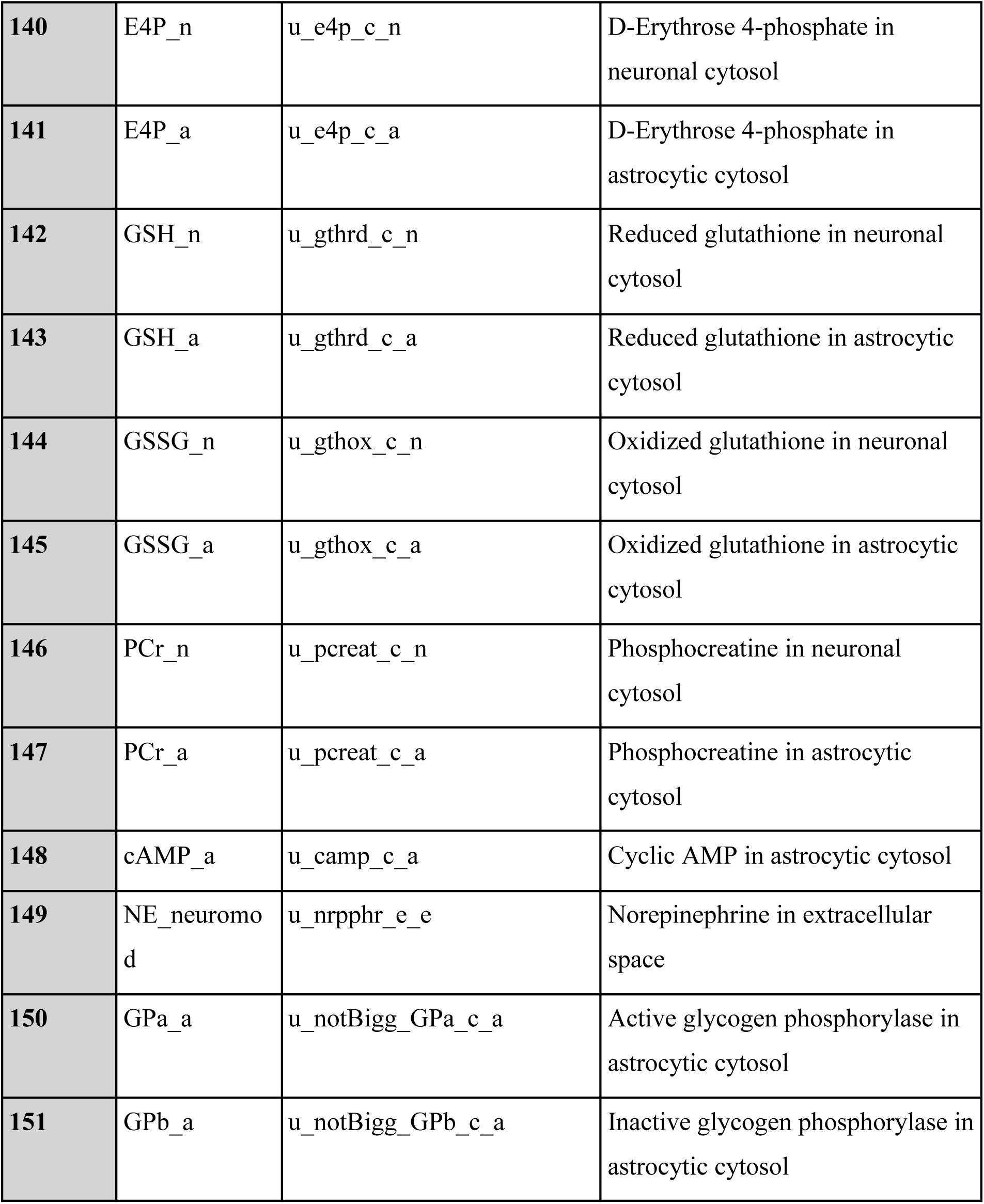

### Supplementary Information File 8

Model variables initial values.

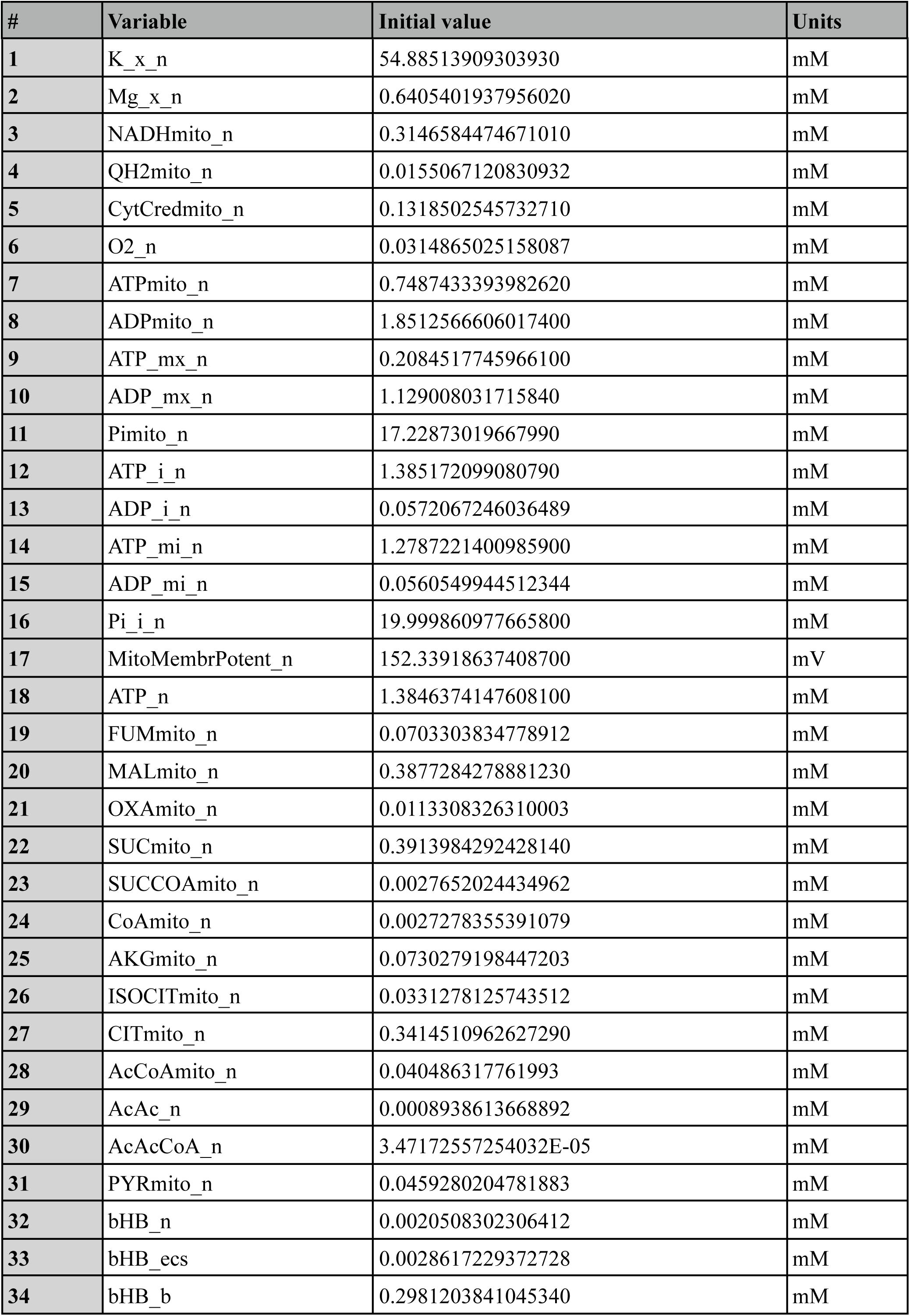
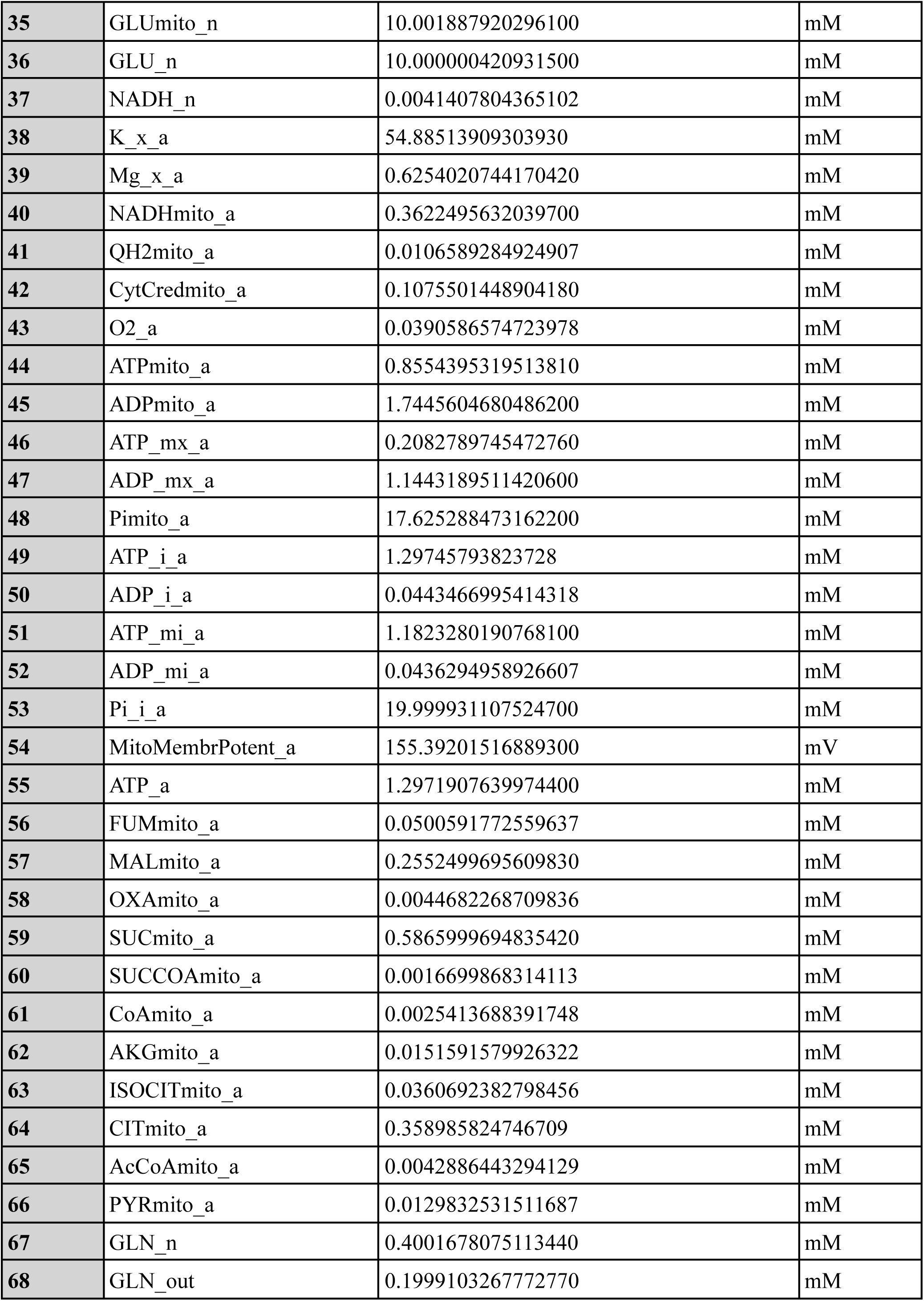
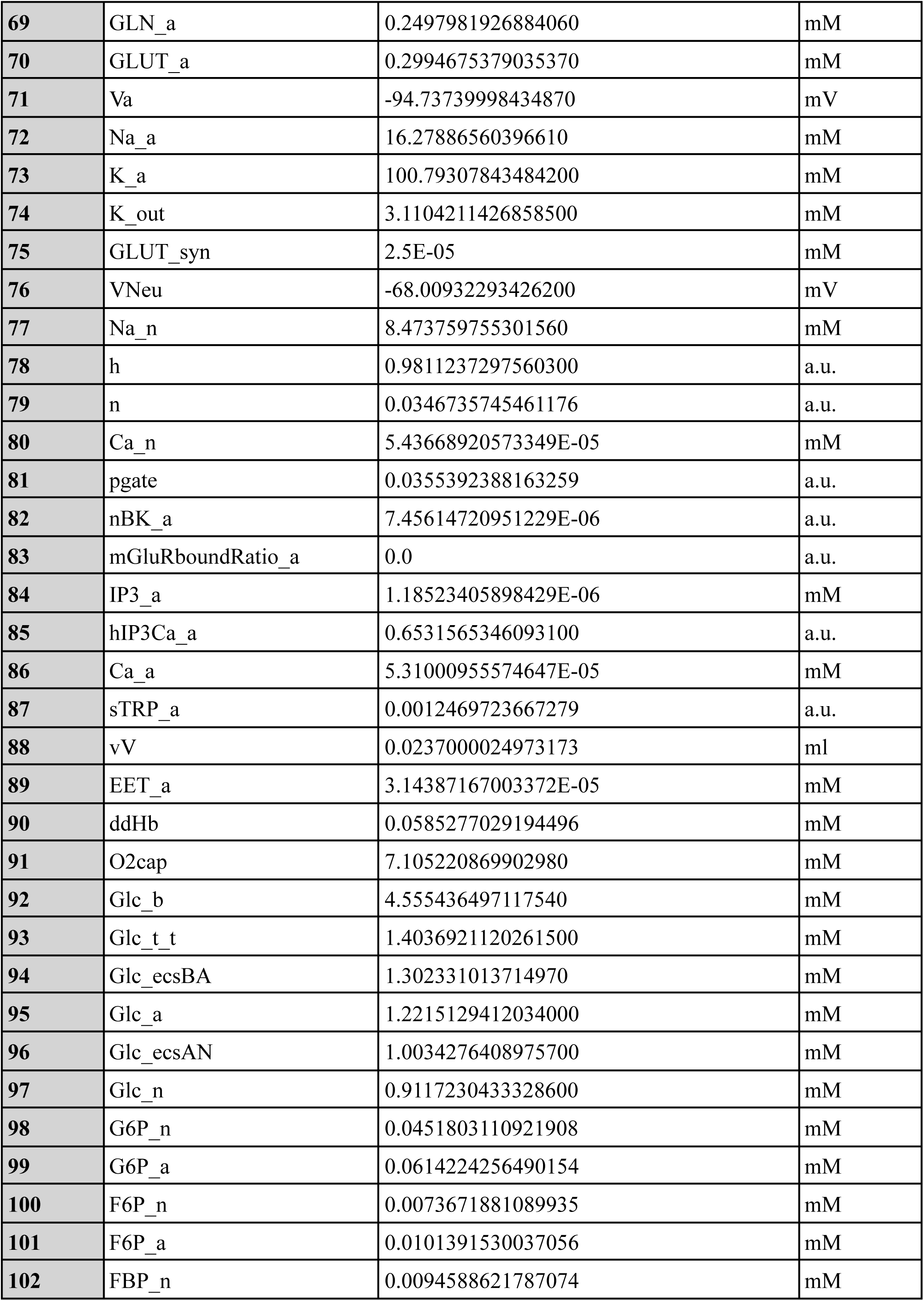
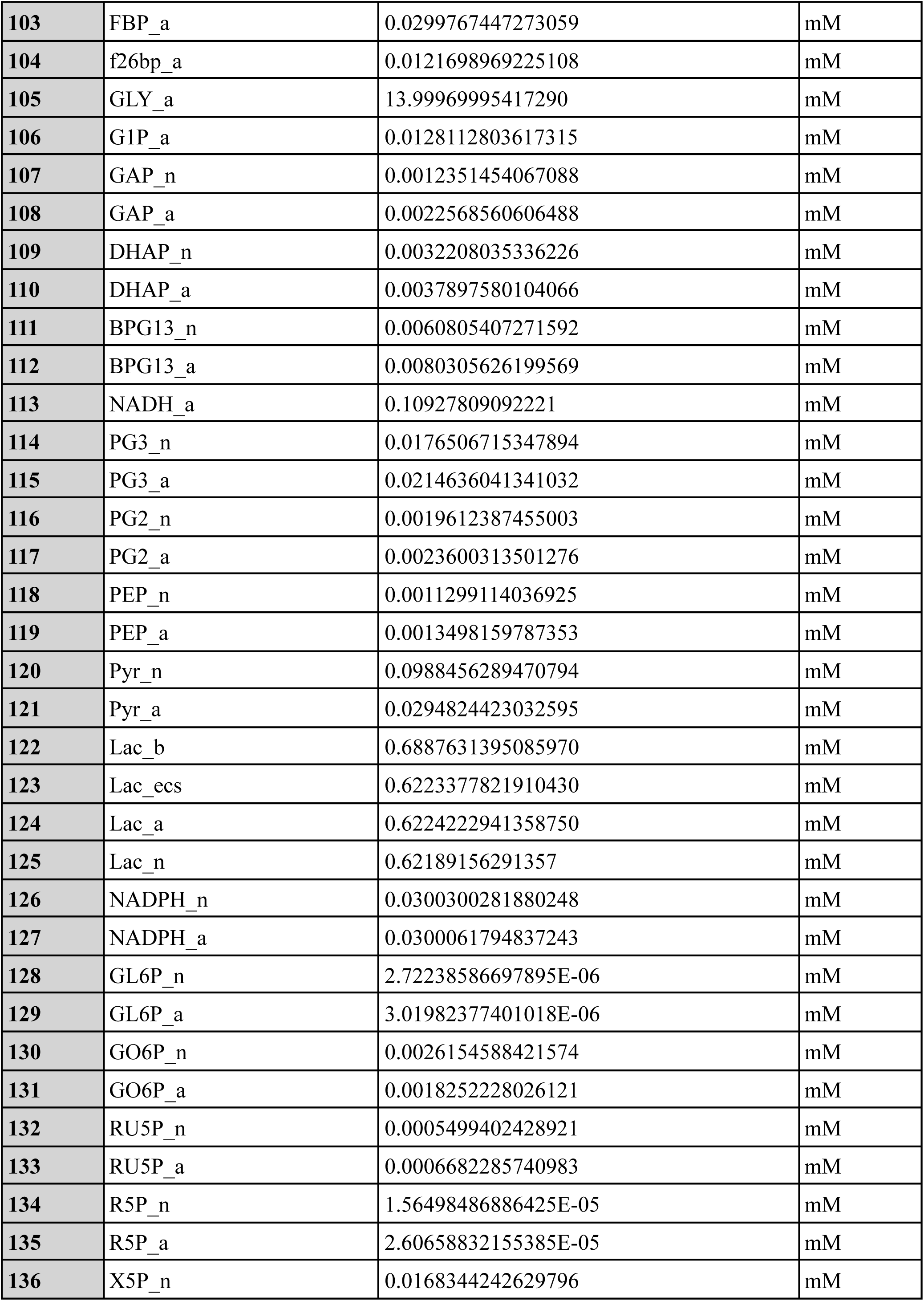
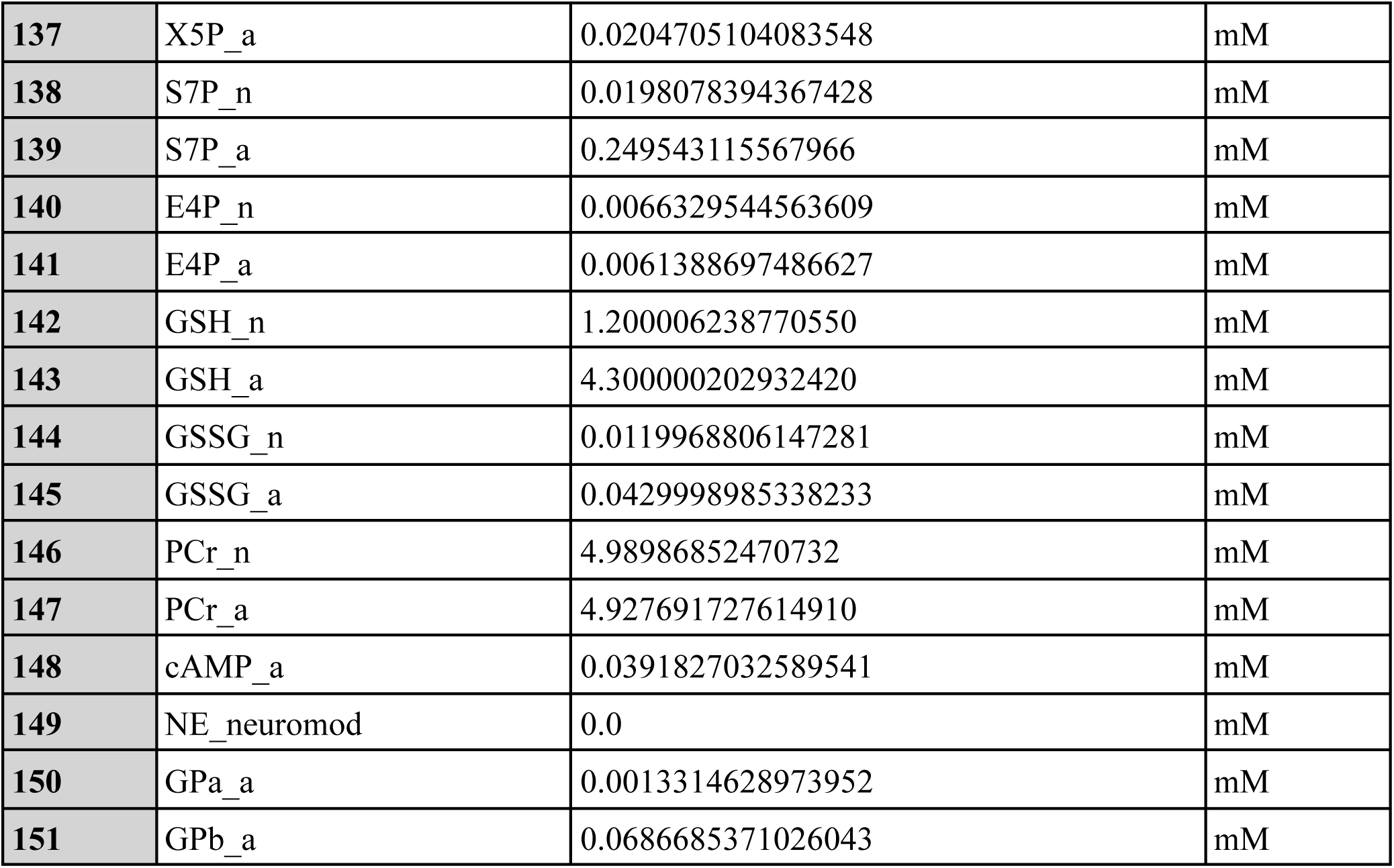

## Notes

### Competing Interest Statement

The authors have declared no competing interest.

### Summary of Updates

Added topological analysis, updated clustering method, text revisions.

## References

Acevedo, A., Torres, F., Kiwi, M., Baeza-Lehnert, F., Barros, L. F., Lee-Liu, D., et al. (2023). Metabolic switch in the aging astrocyte supported via integrative approach comprising network and transcriptome analyses. Aging 15. doi: 10.18632/aging.204663.

Achanta, L. B., and Rae, C. D. (2017). β-Hydroxybutyrate in the Brain: One Molecule, Multiple Mechanisms. Neurochem. Res. 42, 35–49. doi: 10.1007/s11064-016-2099-2.

Amorim, J. A., Coppotelli, G., Rolo, A. P., Palmeira, C. M., Ross, J. M., and Sinclair, D. A. (2022). Mitochondrial and metabolic dysfunction in ageing and age-related diseases. Nat. Rev. Endocrinol. 18, 243–258. doi: 10.1038/s41574-021-00626-7.

Anderson, P. J., and Wright, B. E. (1980). Kinetic models of glycogen metabolism in normal rat liver, morris Hepatom 7787 and host liver. Int. J. Biochem. 12, 361–369. doi: 10.1016/0020-711X(80)90115-9.

Anderson, R., and Prolla, T. (2009). PGC-1alpha in aging and anti-aging interventions. Biochim. Biophys. Acta 1790, 1059–1066. doi: 10.1016/j.bbagen.2009.04.005.

Antonenkov, V. D., Croes, K., Waelkens, E., Van Veldhoven, P. P., and Mannaerts, G. P. (2000). peroxisomes: Acetoacetyl-CoA thiolase from liver peroxisomes. Eur. J. Biochem. 267, 2981–2990. doi: 10.1046/j.1432-1033.2000.01314.x.

Arce-Molina, R., Cortés-Molina, F., Sandoval, P. Y., Galaz, A., Alegría, K., Schirmeier, S., et al. (2020). A highly responsive pyruvate sensor reveals pathway-regulatory role of the mitochondrial pyruvate carrier MPC. eLife 9, e53917. doi: 10.7554/eLife.53917.

Ashok, B. S., Ajith, T. A., and Sivanesan, S. (2017). Hypoxia-inducible factors as neuroprotective agent in Alzheimer’s disease. Clin. Exp. Pharmacol. Physiol. 44, 327–334. doi: 10.1111/1440-1681.12717.

Atkinson, D. E. (1968). Energy charge of the adenylate pool as a regulatory parameter. Interaction with feedback modifiers. Biochemistry 7, 4030–4034. doi: 10.1021/bi00851a033.

Aubert, A., Costalat, R., and Valabrègue, R. (2001). Modelling of the coupling between brain electrical activity and metabolism. Acta Biotheor. 49, 301–326. doi: 10.1023/a:1014286728421.

Baeza-Lehnert, F., Saab, A. S., Gutiérrez, R., Larenas, V., Díaz, E., Horn, M., et al. (2019). Non-Canonical Control of Neuronal Energy Status by the Na+ Pump. Cell Metab. 29, 668–680.e4. doi: 10.1016/j.cmet.2018.11.005.

Barbagallo, M., Belvedere, M., and Dominguez, L. J. (2009). Magnesium homeostasis and aging. Magnes. Res. 22, 235–246. doi: 10.1684/mrh.2009.0187.

Barbagallo, M., and Dominguez, L. (2010). Magnesium and Aging. Curr. Pharm. Des. 16, 832–839. doi: 10.2174/138161210790883679.

Barbagallo, M., Veronese, N., and Dominguez, L. J. (2021). Magnesium in Aging, Health and Diseases. Nutrients 13, 463. doi: 10.3390/nu13020463.

Barden, R. E., Fung, C. H., Utter, M. F., and Scrutton, M. C. (1972). Pyruvate carboxylase from chicken liver. Steady state kinetic studies indicate a “two-site” ping-pong mechanism. J. Biol. Chem. 247, 1323–1333.

Barker, S. J., Raju, R. M., Milman, N. E. P., Wang, J., Davila-Velderrain, J., Gunter-Rahman, F., et al. (2021). MEF2 is a key regulator of cognitive potential and confers resilience to neurodegeneration. Sci. Transl. Med. 13, eabd7695. doi: 10.1126/scitranslmed.abd7695.

Barros, L. (2022). How expensive is the astrocyte? J. Cereb. Blood Flow Metab. 42, 738–745. doi: 10.1177/0271678X221077343.

Barros, L. F., Bittner, C. X., Loaiza, A., and Porras, O. H. (2007). A quantitative overview of glucose dynamics in the gliovascular unit. Glia 55, 1222–1237. doi: 10.1002/glia.20375.

Barros, L. F., San Martín, A., Ruminot, I., Sandoval, P. Y., Fernández-Moncada, I., Baeza-Lehnert, F., et al. (2017). Near-critical GLUT1 and Neurodegeneration: Glucose Transport and Neurodegeneration. J. Neurosci. Res. 95, 2267–2274. doi: 10.1002/jnr.23998.

Bélanger, M., Allaman, I., and Magistretti, P. J. (2011). Brain energy metabolism: focus on astrocyte-neuron metabolic cooperation. Cell Metab. 14, 724–738. doi: 10.1016/j.cmet.2011.08.016.

Bennett, M. R., Farnell, L., and Gibson, W. G. (2008). Origins of blood volume change due to glutamatergic synaptic activity at astrocytes abutting on arteriolar smooth muscle cells. J. Theor. Biol. 250, 172–185. doi: 10.1016/j.jtbi.2007.08.024.

Benton, D., Parker, P. Y., and Donohoe, R. T. (1996). The supply of glucose to the brain and cognitive functioning. J. Biosoc. Sci. 28, 463–479. doi: 10.1017/S0021932000022537.

Berndt, N., Bulik, S., and Holzhütter, H.-G. (2012). Kinetic Modeling of the Mitochondrial Energy Metabolism of Neuronal Cells: The Impact of Reduced α-Ketoglutarate Dehydrogenase Activities on ATP Production and Generation of Reactive Oxygen Species. Int. J. Cell Biol. 2012, 757594. doi: 10.1155/2012/757594.

Berndt, N., Bulik, S., Wallach, I., Wünsch, T., König, M., Stockmann, M., et al. (2018). HEPATOKIN1 is a biochemistry-based model of liver metabolism for applications in medicine and pharmacology. Nat. Commun. 9, 2386. doi: 10.1038/s41467-018-04720-9.

Berndt, N., Kann, O., and Holzhütter, H.-G. (2015). Physiology-Based Kinetic Modeling of Neuronal Energy Metabolism Unravels the Molecular Basis of NAD(P)H Fluorescence Transients. J. Cereb. Blood Flow Metab. 35, 1494–1506. doi: 10.1038/jcbfm.2015.70.

Bezanson, J., Edelman, A., Karpinski, S., and Shah, V. B. (2017). Julia: A Fresh Approach to Numerical Computing. SIAM Rev. 59, 65–98. doi: 10.1137/141000671.

Bishop, N. A., Lu, T., and Yankner, B. A. (2010). Neural mechanisms of ageing and cognitive decline. Nature 464, 529–535. doi: 10.1038/nature08983.

Błaszczyk, J. W. (2020). Energy Metabolism Decline in the Aging Brain—Pathogenesis of Neurodegenerative Disorders. Metabolites 10, 450. doi: 10.3390/metabo10110450.

Bonvento, G., and Bolaños, J. P. (2021). Astrocyte-neuron metabolic cooperation shapes brain activity. Cell Metab. 33, 1546–1564. doi: 10.1016/j.cmet.2021.07.006.

Borst, P. (2020). The malate–aspartate shuttle (Borst cycle): How it started and developed into a major metabolic pathway. IUBMB Life 72, 2241–2259. doi: 10.1002/iub.2367.

Botman, D., Tigchelaar, W., and Van Noorden, C. J. F. (2014). Determination of glutamate dehydrogenase activity and its kinetics in mouse tissues using metabolic mapping (quantitative enzyme histochemistry). J. Histochem. Cytochem. Off. J. Histochem. Soc. 62, 802–812. doi: 10.1369/0022155414549071.

Bouzier-Sore, A.-K., and Bolaños, J. P. (2015). Uncertainties in pentose-phosphate pathway flux assessment underestimate its contribution to neuronal glucose consumption: relevance for neurodegeneration and aging. Front. Aging Neurosci. 7. doi: 10.3389/fnagi.2015.00089.

Bouzier-Sore, A.-K., Voisin, P., Bouchaud, V., Bezancon, E., Franconi, J.-M., and Pellerin, L. (2006). Competition between glucose and lactate as oxidative energy substrates in both neurons and astrocytes: a comparative NMR study. Eur. J. Neurosci. 24, 1687–1694. doi: 10.1111/j.1460-9568.2006.05056.x.

Bradshaw, P. (2019). Cytoplasmic and Mitochondrial NADPH-Coupled Redox Systems in the Regulation of Aging. Nutrients 11, 504. doi: 10.3390/nu11030504.

Braidy, N., Poljak, A., Grant, R., Jayasena, T., Mansour, H., Chan-Ling, T., et al. (2014). Mapping NAD+ metabolism in the brain of ageing Wistar rats: potential targets for influencing brain senescence. Biogerontology 15, 177–198. doi: 10.1007/s10522-013-9489-5.

Breslin, K., Wade, J. J., Wong-Lin, K., Harkin, J., Flanagan, B., Van Zalinge, H., et al. (2018). Potassium and sodium microdomains in thin astroglial processes: A computational model study. PLOS Comput. Biol. 14, e1006151. doi: 10.1371/journal.pcbi.1006151.

Brilkova, M., Nigri, M., Kumar, H. S., Moore, J., Mantovani, M., Keller, C., et al. (2022). Error-prone protein synthesis recapitulates early symptoms of Alzheimer disease in aging mice. Cell Rep. 40, 111433. doi: 10.1016/j.celrep.2022.111433.

Brocard, J. B., Tassetto, M., and Reynolds, I. J. (2001). Quantitative evaluation of mitochondrial calcium content in rat cortical neurones following a glutamate stimulus. J. Physiol. 531, 793–805. doi: 10.1111/j.1469-7793.2001.0793h.x.

Bröer, S., and Brookes, N. (2001). Transfer of glutamine between astrocytes and neurons: Glutamine transfer between astrocytes and neurons. J. Neurochem. 77, 705–719. doi: 10.1046/j.1471-4159.2001.00322.x.

Brunk, E., Sahoo, S., Zielinski, D. C., Altunkaya, A., Dräger, A., Mih, N., et al. (2018). Recon3D enables a three-dimensional view of gene variation in human metabolism. Nat. Biotechnol. 36, 272–281. doi: 10.1038/nbt.4072.

Burtscher, J., Mallet, R. T., Burtscher, M., and Millet, G. P. (2021). Hypoxia and brain aging: Neurodegeneration or neuroprotection? Ageing Res. Rev. 68, 101343. doi: 10.1016/j.arr.2021.101343.

Byrne, J. H., Heidelberger, R., and Waxham, M. N. eds. (2014). From molecules to networks: an introduction to cellular and molecular neuroscience. Third edition. Amsterdam ; Boston: Elsevier/AP, Academic Press is an imprint of Elsevier.

Cai, L., Sutter, B. M., Li, B., and Tu, B. P. (2011). Acetyl-CoA Induces Cell Growth and Proliferation by Promoting the Acetylation of Histones at Growth Genes. Mol. Cell 42, 426–437. doi: 10.1016/j.molcel.2011.05.004.

Calvetti, D., Capo Rangel, G., Gerardo Giorda, L., and Somersalo, E. (2018). A computational model integrating brain electrophysiology and metabolism highlights the key role of extracellular potassium and oxygen. J. Theor. Biol. 446, 238–258. doi: 10.1016/j.jtbi.2018.02.029.

Calvetti, D., and Somersalo, E. (2011). Dynamic activation model for a glutamatergic neurovascular unit. J. Theor. Biol. 274, 12–29. doi: 10.1016/j.jtbi.2010.12.007.

Campisi, J., Kapahi, P., Lithgow, G. J., Melov, S., Newman, J. C., and Verdin, E. (2019). From discoveries in ageing research to therapeutics for healthy ageing. Nature 571, 183–192. doi: 10.1038/s41586-019-1365-2.

Cantó, C., Gerhart-Hines, Z., Feige, J. N., Lagouge, M., Noriega, L., Milne, J. C., et al. (2009). AMPK regulates energy expenditure by modulating NAD+ metabolism and SIRT1 activity. Nature 458, 1056–1060. doi: 10.1038/nature07813.

Chakrabarti, A., Oehme, I., Witt, O., Oliveira, G., Sippl, W., Romier, C., et al. (2015). HDAC8: a multifaceted target for therapeutic interventions. Trends Pharmacol. Sci. 36, 481–492. doi: 10.1016/j.tips.2015.04.013.

Chang, A., Jeske, L., Ulbrich, S., Hofmann, J., Koblitz, J., Schomburg, I., et al. (2021). BRENDA, the ELIXIR core data resource in 2021: new developments and updates. Nucleic Acids Res. 49, D498–D508. doi: 10.1093/nar/gkaa1025.

Chaudhry, F. A., Reimer, R. J., Krizaj, D., Barber, D., Storm-Mathisen, J., Copenhagen, D. R., et al. (1999). Molecular Analysis of System N Suggests Novel Physiological Roles in Nitrogen Metabolism and Synaptic Transmission. Cell 99, 769–780. doi: 10.1016/S0092-8674(00)81674-8.

Choi, I.-Y., and Gruetter, R. eds. (2012). Neural Metabolism In Vivo. Boston, MA: Springer US doi: 10.1007/978-1-4614-1788-0.

Chowdhury, G. M., Jiang, L., Rothman, D. L., and Behar, K. L. (2014). The Contribution of Ketone Bodies to Basal and Activity-Dependent Neuronal Oxidation in Vivo. J. Cereb. Blood Flow Metab. 34, 1233–1242. doi: 10.1038/jcbfm.2014.77.

Cloutier, M., Bolger, F. B., Lowry, J. P., and Wellstead, P. (2009). An integrative dynamic model of brain energy metabolism using in vivo neurochemical measurements. J. Comput. Neurosci. 27, 391–414. doi: 10.1007/s10827-009-0152-8.

Coggan, J. S., Keller, D., Calì, C., Lehväslaiho, H., Markram, H., Schürmann, F., et al. (2018). Norepinephrine stimulates glycogenolysis in astrocytes to fuel neurons with lactate. PLOS Comput. Biol. 14, e1006392. doi: 10.1371/journal.pcbi.1006392.

Coggan, J. S., Keller, D., Markram, H., Schürmann, F., and Magistretti, P. J. (2020). Excitation states of metabolic networks predict dose-response fingerprinting and ligand pulse phase signalling. J. Theor. Biol. 487, 110123. doi: 10.1016/j.jtbi.2019.110123.

Cox, M. F., Hascup, E. R., Bartke, A., and Hascup, K. N. (2022). Friend or Foe? Defining the Role of Glutamate in Aging and Alzheimer’s Disease. Front. Aging 3, 929474. doi: 10.3389/fragi.2022.929474.

Cui, L., Jeong, H., Borovecki, F., Parkhurst, C. N., Tanese, N., and Krainc, D. (2006). Transcriptional repression of PGC-1alpha by mutant huntingtin leads to mitochondrial dysfunction and neurodegeneration. Cell 127, 59–69. doi: 10.1016/j.cell.2006.09.015.

Curtis, W. M., Seeds, W. A., Mattson, M. P., and Bradshaw, P. C. (2022). NADPH and Mitochondrial Quality Control as Targets for a Circadian-Based Fasting and Exercise Therapy for the Treatment of Parkinson’s Disease. Cells 11, 2416. doi: 10.3390/cells11152416.

Datta, S., and Chakrabarti, N. (2018). Age related rise in lactate and its correlation with lactate dehydrogenase (LDH) status in post-mitochondrial fractions isolated from different regions of brain in mice. Neurochem. Int. 118, 23–33. doi: 10.1016/j.neuint.2018.04.007.

De Simone, G., di Masi, A., Polticelli, F., and Ascenzi, P. (2018). Human nitrobindin: the first example of an all-β-barrel ferric heme-protein that catalyzes peroxynitrite detoxification. FEBS Open Bio 8, 2002–2010. doi: 10.1002/2211-5463.12534.

Deczkowska, A., Matcovitch-Natan, O., Tsitsou-Kampeli, A., Ben-Hamo, S., Dvir-Szternfeld, R., Spinrad, A., et al. (2017). Mef2C restrains microglial inflammatory response and is lost in brain ageing in an IFN-I-dependent manner. Nat. Commun. 8, 717. doi: 10.1038/s41467-017-00769-0.

Dienel, G. A. (2012). Brain Lactate Metabolism: The Discoveries and the Controversies. J. Cereb. Blood Flow Metab. 32, 1107–1138. doi: 10.1038/jcbfm.2011.175.

DiNuzzo, M., Mangia, S., Maraviglia, B., and Giove, F. (2010). Glycogenolysis in astrocytes supports blood-borne glucose channeling not glycogen-derived lactate shuttling to neurons: evidence from mathematical modeling. J. Cereb. Blood Flow Metab. Off. J. Int. Soc. Cereb. Blood Flow Metab. 30, 1895–1904. doi: 10.1038/jcbfm.2010.151.

Disterhoft, J. F., and Oh, M. M. (2007). Alterations in intrinsic neuronal excitability during normal aging. Aging Cell 6, 327–336. doi: 10.1111/j.1474-9726.2007.00297.x.

Dombrowski, G. J., Cheung, G. P., and Swiatek, K. R. (1977). Evidence for the existence of enzymatic variants of β-hydroxybutyrate dehydrogenase from rat liver and brain mitochondria. Life Sci. 21, 1821–1829. doi: 10.1016/0024-3205(77)90164-3.

Dong, Y., and Brewer, G. J. (2019). Global Metabolic Shifts in Age and Alzheimer’s Disease Mouse Brains Pivot at NAD+/NADH Redox Sites. J. Alzheimers Dis. 71, 119–140. doi: 10.3233/JAD-190408.

Duarte, J. M. N., and Gruetter, R. (2013). Glutamatergic and GABAergic energy metabolism measured in the rat brain by 13 C NMR spectroscopy at 14.1 T. J. Neurochem. 126, 579–590. doi: 10.1111/jnc.12333.

Dyson, F. (2004). A meeting with Enrico Fermi. Nature 427, 297–297. doi: 10.1038/427297a.

Eap, B., Nomura, M., Panda, O., Garcia, T. Y., King, C. D., Rose, J. P., et al. (2022). Ketone body metabolism declines with age in mice in a sex-dependent manner. Physiology doi: 10.1101/2022.10.05.511032.

Emmons, M. F., Bennett, R. L., Riva, A., Zhang, C., Macaulay, R., Dupéré-Richér, D., et al. (2022). HDAC8-mediated inhibition of EP300 drives a neural crest-like transcriptional state that increases melanoma brain metastasis. Cancer Biology doi: 10.1101/2022.10.12.511971.

Erecińska, M., and Silver, I. A. (1989). ATP and brain function. J. Cereb. Blood Flow Metab. Off. J. Int. Soc. Cereb. Blood Flow Metab. 9, 2–19. doi: 10.1038/jcbfm.1989.2.

Erecińska, M., and Silver, I. A. (1994). Ions and energy in mammalian brain. Prog. Neurobiol. 43, 37–71. doi: 10.1016/0301-0082(94)90015-9.

Fang, E. F., Lautrup, S., Hou, Y., Demarest, T. G., Croteau, D. L., Mattson, M. P., et al. (2017). NAD + in Aging: Molecular Mechanisms and Translational Implications. Trends Mol. Med. 23, 899–916. doi: 10.1016/j.molmed.2017.08.001.

Featherstone, D. E. (2010). Intercellular Glutamate Signaling in the Nervous System and Beyond. ACS Chem. Neurosci. 1, 4–12. doi: 10.1021/cn900006n.

Fillenz, M., and Lowry, J. P. (1998). Studies of the source of glucose in the extracellular compartment of the rat brain. Dev. Neurosci. 20, 365–368. doi: 10.1159/000017332.

Fink, B. D., Bai, F., Yu, L., Sheldon, R. D., Sharma, A., Taylor, E. B., et al. (2018). Oxaloacetic acid mediates ADP-dependent inhibition of mitochondrial complex II–driven respiration. J. Biol. Chem. 293, 19932–19941. doi: 10.1074/jbc.RA118.005144.

Finkel, T., Deng, C.-X., and Mostoslavsky, R. (2009). Recent progress in the biology and physiology of sirtuins. Nature 460, 587–591. doi: 10.1038/nature08197.

Flanagan, B., McDaid, L., Wade, J., Wong-Lin, K., and Harkin, J. (2018). A computational study of astrocytic glutamate influence on post-synaptic neuronal excitability. PLoS Comput. Biol. 14, e1006040. doi: 10.1371/journal.pcbi.1006040.

Fraser, C. L., and Arieff, A. I. (2001). Na-K-ATPase activity decreases with aging in female rat brain synaptosomes. Am. J. Physiol. Renal Physiol. 281, F674–678. doi: 10.1152/ajprenal.2001.281.4.F674.

Frezza, C. (2017). Mitochondrial metabolites: undercover signalling molecules. Interface Focus 7, 20160100. doi: 10.1098/rsfs.2016.0100.

Gaitonde, M. K., Murray, E., and Cunningham, V. J. (1989). Effect of 6-phosphogluconate on phosphoglucose isomerase in rat brain in vitro and in vivo. J. Neurochem. 52, 1348–1352. doi: 10.1111/j.1471-4159.1989.tb09178.x.

Garcia, S., Nissanka, N., Mareco, E. A., Rossi, S., Peralta, S., Diaz, F., et al. (2018). Overexpression of PGC-1α in aging muscle enhances a subset of young-like molecular patterns. Aging Cell 17, e12707. doi: 10.1111/acel.12707.

Garrett, R., and Grisham, C. M. (2013). Biochemistry. 5th ed. Belmont, CA: Brooks/Cole, Cengage Learning.

Ge, I., Kirschen, G. W., and Wang, X. (2021). Shifted Dynamics of Glucose Metabolism in the Hippocampus During Aging. Front. Aging Neurosci. 13, 700306. doi: 10.3389/fnagi.2021.700306.

Ghosh, D., Levault, K. R., and Brewer, G. J. (2014). Relative importance of redox buffers GSH and NAD(P)H in age-related neurodegeneration and Alzheimer disease-like mouse neurons. Aging Cell 13, 631–640. doi: 10.1111/acel.12216.

Gilbert, H. F., Lennox, B. J., Mossman, C. D., and Carle, W. C. (1981). The relation of acyl transfer to the overall reaction of thiolase I from porcine heart. J. Biol. Chem. 256, 7371–7377.

Grimm, A., and Eckert, A. (2017). Brain aging and neurodegeneration: from a mitochondrial point of view. J. Neurochem. 143, 418–431. doi: 10.1111/jnc.14037.

Guarente, L. (2006). Sirtuins as potential targets for metabolic syndrome. Nature 444, 868–874. doi: 10.1038/nature05486.

Gut, P., and Verdin, E. (2013). The nexus of chromatin regulation and intermediary metabolism. Nature 502, 489–498. doi: 10.1038/nature12752.

Hädel, S., Wirth, C., Rapp, M., Gallinat, J., and Schubert, F. (2013). Effects of age and sex on the concentrations of glutamate and glutamine in the human brain: Brain Glutamate and Glutamine With Age and Sex. J. Magn. Reson. Imaging 38, 1480–1487. doi: 10.1002/jmri.24123.

Hagberg, A. A., Schult, D. A., and Swart, P. J. (2008). Exploring Network Structure, Dynamics, and Function using NetworkX. in Proceedings of the 7th Python in Science Conference, eds. G. Varoquaux, T. Vaught, and J. Millman (Pasadena, CA USA), 11–15.

Halestrap, A. P., and Denton, R. M. (1974). Specific inhibition of pyruvate transport in rat liver mitochondria and human erythrocytes by α-cyano-4-hydroxycinnamate (Short Communication). Biochem. J. 138, 313–316. doi: 10.1042/bj1380313.

Hersh, L. B., and Jencks, W. P. (1967). Isolation of an enzyme-coenzyme A intermediate from succinyl coenzyme A-acetoacetate coenzyme A transferase. J. Biol. Chem. 242, 339–340.

Hertz, L., and Rothman, D. (2017). Glutamine-Glutamate Cycle Flux Is Similar in Cultured Astrocytes and Brain and Both Glutamate Production and Oxidation Are Mainly Catalyzed by Aspartate Aminotransferase. Biology 6, 17. doi: 10.3390/biology6010017.

Horowitz, A. M., Fan, X., Bieri, G., Smith, L. K., Sanchez-Diaz, C. I., Schroer, A. B., et al. (2020). Blood factors transfer beneficial effects of exercise on neurogenesis and cognition to the aged brain. Science 369, 167–173. doi: 10.1126/science.aaw2622.

Hosseini, L., Majdi, A., Sadigh-Eteghad, S., Farajdokht, F., Ziaee, M., Rahigh Aghsan, S., et al. (2022). Coenzyme Q10 ameliorates aging-induced memory deficits via modulation of apoptosis, oxidative stress, and mitophagy in aged rats. Exp. Gerontol. 168, 111950. doi: 10.1016/j.exger.2022.111950.

Hou, Y., Dan, X., Babbar, M., Wei, Y., Hasselbalch, S. G., Croteau, D. L., et al. (2019). Ageing as a risk factor for neurodegenerative disease. Nat. Rev. Neurol. 15, 565–581. doi: 10.1038/s41582-019-0244-7.

Howarth, C., Gleeson, P., and Attwell, D. (2012). Updated Energy Budgets for Neural Computation in the Neocortex and Cerebellum. J. Cereb. Blood Flow Metab. 32, 1222–1232. doi: 10.1038/jcbfm.2012.35.

Huang, C.-Y., Oka, S.-I., Xu, X., Chen, C.-F., Tung, C.-Y., Chang, Y.-Y., et al. (2022). PERM1 regulates genes involved in fatty acid metabolism in the heart by interacting with PPARα and PGC-1α. Sci. Rep. 12, 14576. doi: 10.1038/s41598-022-18885-3.

Huang, H., Zhou, F., Zhou, S., and Qiu, M. (2021). MYRF: A Mysterious Membrane-Bound Transcription Factor Involved in Myelin Development and Human Diseases. Neurosci. Bull. 37, 881–884. doi: 10.1007/s12264-021-00678-9.

Hupfeld, K. E., Hyatt, H. W., Alvarez Jerez, P., Mikkelsen, M., Hass, C. J., Edden, R. A. E., et al. (2021). In Vivo Brain Glutathione is Higher in Older Age and Correlates with Mobility. Cereb. Cortex 31, 4576–4594. doi: 10.1093/cercor/bhab107.

Huth, W., and Menke, R. (1982). Regulation of ketogenesis. Mitochondrial acetyl-CoA acetyltransferase from rat liver: initial-rate kinetics in the presence of the product CoASH reveal intermediary plateau regions. Eur. J. Biochem. 128, 413–419.

Huynh, Q. K., Sakakibara, R., Watanabe, T., and Wada, H. (1980). Glutamic oxaloacetic transaminase isozymes from rat liver. Purification and physicochemical characterization. J. Biochem. (Tokyo) 88, 231–239.

Iskusnykh, I. Y., Zakharova, A. A., and Pathak, D. (2022). Glutathione in Brain Disorders and Aging. Mol. Basel Switz. 27, 324. doi: 10.3390/molecules27010324.

Ivanisevic, J., Stauch, K., Petrascheck, M., Benton, H., Epstein, A., Fang, M., et al. (2016). Metabolic drift in the aging brain. Aging 8, 1000–1020. doi: 10.18632/aging.100961.

Izzo, A., Manco, R., Bonfiglio, F., Calì, G., De Cristofaro, T., Patergnani, S., et al. (2014). NRIP1/RIP140 siRNA-mediated attenuation counteracts mitochondrial dysfunction in Down syndrome. Hum. Mol. Genet. 23, 4406–4419. doi: 10.1093/hmg/ddu157.

Jolivet, R., Allaman, I., Pellerin, L., Magistretti, P. J., and Weber, B. (2010). Comment on Recent Modeling Studies of Astrocyte–Neuron Metabolic Interactions. J. Cereb. Blood Flow Metab. 30, 1982–1986. doi: 10.1038/jcbfm.2010.132.

Jolivet, R., Coggan, J. S., Allaman, I., and Magistretti, P. J. (2015). Multi-timescale Modeling of Activity-Dependent Metabolic Coupling in the Neuron-Glia-Vasculature Ensemble. PLOS Comput. Biol. 11, e1004036. doi: 10.1371/journal.pcbi.1004036.

Jones, T. T., and Brewer, G. J. (2009). Critical age-related loss of cofactors of neuron cytochrome C oxidase reversed by estrogen. Exp. Neurol. 215, 212–219. doi: 10.1016/j.expneurol.2008.09.011.

Jung, W. B., Im, G. H., Jiang, H., and Kim, S.-G. (2021). Early fMRI responses to somatosensory and optogenetic stimulation reflect neural information flow. Proc. Natl. Acad. Sci. 118, e2023265118. doi: 10.1073/pnas.2023265118.

Kaiser, L. G., Schuff, N., Cashdollar, N., and Weiner, M. W. (2005). Age-related glutamate and glutamine concentration changes in normal human brain: 1H MR spectroscopy study at 4 T. Neurobiol. Aging 26, 665–672. doi: 10.1016/j.neurobiolaging.2004.07.001.

Kang, I., Chu, C. T., and Kaufman, B. A. (2018). The mitochondrial transcription factor TFAM in neurodegeneration: emerging evidence and mechanisms. FEBS Lett. 592, 793–811. doi: 10.1002/1873-3468.12989.

Katsyuba, E., Mottis, A., Zietak, M., De Franco, F., van der Velpen, V., Gariani, K., et al. (2018). De novo NAD+ synthesis enhances mitochondrial function and improves health. Nature 563, 354–359. doi: 10.1038/s41586-018-0645-6.

Kauffman, F. C., Brown, J. G., Passonneau, J. V., and Lowry, O. H. (1969). Effects of changes in brain metabolism on levels of pentose phosphate pathway intermediates. J. Biol. Chem. 244, 3647–3653.

Keenan, A. B., Torre, D., Lachmann, A., Leong, A. K., Wojciechowicz, M. L., Utti, V., et al. (2019). ChEA3: transcription factor enrichment analysis by orthogonal omics integration. Nucleic Acids Res. 47, W212–W224. doi: 10.1093/nar/gkz446.

Kelly, M. P. (2018). Cyclic nucleotide signaling changes associated with normal aging and age-related diseases of the brain. Cell. Signal. 42, 281–291. doi: 10.1016/j.cellsig.2017.11.004.

Kety, S. S. (1957). “The general metabolism of the brain in vivo,” in Metabolism of the nervous system (Elsevier), 221–237.

Kim, S. Y., Yang, C.-S., Lee, H.-M., Kim, J. K., Kim, Y.-S., Kim, Y.-R., et al. (2018). ESRRA (estrogen-related receptor α) is a key coordinator of transcriptional and post-translational activation of autophagy to promote innate host defense. Autophagy 14, 152–168. doi: 10.1080/15548627.2017.1339001.

King, Z. A., Lu, J., Dräger, A., Miller, P., Federowicz, S., Lerman, J. A., et al. (2016). BiGG Models: A platform for integrating, standardizing and sharing genome-scale models. Nucleic Acids Res. 44, D515–D522. doi: 10.1093/nar/gkv1049.

Kiyatkin, E. A., and Lenoir, M. (2012). Rapid fluctuations in extracellular brain glucose levels induced by natural arousing stimuli and intravenous cocaine: fueling the brain during neural activation. J. Neurophysiol. 108, 1669–1684. doi: 10.1152/jn.00521.2012.

Koga, M., Serritella, A. V., Messmer, M. M., Hayashi-Takagi, A., Hester, L. D., Snyder, S. H., et al. (2011). Glutathione is a physiologic reservoir of neuronal glutamate. Biochem. Biophys. Res. Commun. 409, 596–602. doi: 10.1016/j.bbrc.2011.04.087.

Köhler, S., Schmidt, H., Fülle, P., Hirrlinger, J., and Winkler, U. (2020). A Dual Nanosensor Approach to Determine the Cytosolic Concentration of ATP in Astrocytes. Front. Cell. Neurosci. 14, 565921. doi: 10.3389/fncel.2020.565921.

Krishnan, G. P., Filatov, G., Shilnikov, A., and Bazhenov, M. (2015). Electrogenic properties of the Na+/K + ATPase control transitions between normal and pathological brain states. J. Neurophysiol. 113, 3356–3374. doi: 10.1152/jn.00460.2014.

Kulkarni, A. S., Gubbi, S., and Barzilai, N. (2020). Benefits of Metformin in Attenuating the Hallmarks of Aging. Cell Metab. 32, 15–30. doi: 10.1016/j.cmet.2020.04.001.

Kumar, A., and Foster, T. C. (2007). “Neurophysiology of Old Neurons and Synapses,” in Brain Aging: Models, Methods, and Mechanisms Frontiers in Neuroscience., ed. D. R. Riddle (Boca Raton (FL): CRC Press/Taylor & Francis). Available at: http://www.ncbi.nlm.nih.gov/books/NBK3882/ [Accessed June 28, 2023].

Kuroda, T., Yasuda, S., Tachi, S., Matsuyama, S., Kusakawa, S., Tano, K., et al. (2019). SALL3 expression balance underlies lineage biases in human induced pluripotent stem cell differentiation. Nat. Commun. 10, 2175. doi: 10.1038/s41467-019-09511-4.

Lajtha, A., and Reith, M. E. A. eds. (2007). *Handbook of Neurochemistry and Molecular Neurobiology*. Boston, MA: Springer US doi: 10.1007/978-0-387-30380-2.

Lambeth, M. J., and Kushmerick, M. J. (2002). A computational model for glycogenolysis in skeletal muscle. Ann. Biomed. Eng. 30, 808–827.

Ledford, H. (2010). Ageing: Much ado about ageing. Nature 464, 480–481. doi: 10.1038/464480a.

Lee, C. S., Lee, C., Hu, T., Nguyen, J. M., Zhang, J., Martin, M. V., et al. (2011). Loss of nuclear factor E2-related factor 1 in the brain leads to dysregulation of proteasome gene expression and neurodegeneration. Proc. Natl. Acad. Sci. U. S. A. 108, 8408–8413. doi: 10.1073/pnas.1019209108.

Lee, J. V., Carrer, A., Shah, S., Snyder, N. W., Wei, S., Venneti, S., et al. (2014). Akt-dependent metabolic reprogramming regulates tumor cell histone acetylation. Cell Metab. 20, 306–319. doi: 10.1016/j.cmet.2014.06.004.

Lee, J. W., Ko, J., Ju, C., and Eltzschig, H. K. (2019). Hypoxia signaling in human diseases and therapeutic targets. Exp. Mol. Med. 51, 1–13. doi: 10.1038/s12276-019-0235-1.

Lee, S., Devanney, N. A., Golden, L. R., Smith, C. T., Schwartz, J. L., Walsh, A. E., et al. (2023). APOE modulates microglial immunometabolism in response to age, amyloid pathology, and inflammatory challenge. Cell Rep. 42, 112196. doi: 10.1016/j.celrep.2023.112196.

Lerchundi, R., Fernández-Moncada, I., Contreras-Baeza, Y., Sotelo-Hitschfeld, T., Mächler, P., Wyss, M. T., et al. (2015). NH4+ triggers the release of astrocytic lactate via mitochondrial pyruvate shunting. Proc. Natl. Acad. Sci. 112, 11090–11095. doi: 10.1073/pnas.1508259112.

Li, L.-Z., Zhao, Y.-W., Pan, H.-X., Xiang, Y.-Q., Wang, Y.-G., Xu, Q., et al. (2022). Association of rare PPARGC1A variants with Parkinson’s disease risk. J. Hum. Genet. 67, 687–690. doi: 10.1038/s10038-022-01074-5.

Li, W., Yu, J., Liu, Y., Huang, X., Abumaria, N., Zhu, Y., et al. (2014). Elevation of brain magnesium prevents synaptic loss and reverses cognitive deficits in Alzheimer’s disease mouse model. Mol. Brain 7, 65. doi: 10.1186/s13041-014-0065-y.

Listrom, D. C., Morizono, H., Rajagopal, S. B., McCANN, T. M., Tuchman, M., and Allewell, M. N. (1997). Expression, purification, and characterization of recombinant human glutamine synthetase. Biochem. J. 328, 159–163. doi: 10.1042/bj3280159.

López-Otín, C., Blasco, M. A., Partridge, L., Serrano, M., and Kroemer, G. (2013). The Hallmarks of Aging. Cell 153, 1194–1217. doi: 10.1016/j.cell.2013.05.039.

López-Otín, C., Blasco, M. A., Partridge, L., Serrano, M., and Kroemer, G. (2023). Hallmarks of aging: An expanding universe. Cell 186, 243–278. doi: 10.1016/j.cell.2022.11.001.

Luder, A. S., Parks, J. K., Frerman, F., and Parker, W. D. (1990). Inactivation of beef brain alpha-ketoglutarate dehydrogenase complex by valproic acid and valproic acid metabolites. Possible mechanism of anticonvulsant and toxic actions. J. Clin. Invest. 86, 1574–1581. doi: 10.1172/JCI114877.

Luo, Z., Tian, M., Yang, G., Tan, Q., Chen, Y., Li, G., et al. (2022). Hypoxia signaling in human health and diseases: implications and prospects for therapeutics. Signal Transduct. Target. Ther. 7, 218. doi: 10.1038/s41392-022-01080-1.

Mächler, P., Wyss, M. T., Elsayed, M., Stobart, J., Gutierrez, R., von Faber-Castell, A., et al. (2016). In Vivo Evidence for a Lactate Gradient from Astrocytes to Neurons. Cell Metab. 23, 94–102. doi: 10.1016/j.cmet.2015.10.010.

Mahan, D. E., Mushahwar, I. K., and Koeppe, R. E. (1975). Purification and properties of rat brain pyruvate carboxylase. Biochem. J. 145, 25–35. doi: 10.1042/bj1450025.

Mahmoud, S., Gharagozloo, M., Simard, C., and Gris, D. (2019). Astrocytes Maintain Glutamate Homeostasis in the CNS by Controlling the Balance between Glutamate Uptake and Release. Cells 8, 184. doi: 10.3390/cells8020184.

Majmundar, A. J., Wong, W. J., and Simon, M. C. (2010). Hypoxia-inducible factors and the response to hypoxic stress. Mol. Cell 40, 294–309. doi: 10.1016/j.molcel.2010.09.022.

Maletic-Savatic, M., Vingara, L. K., Manganas, L. N., Li, Y., Zhang, S., Sierra, A., et al. (2008). Metabolomics of Neural Progenitor Cells: A Novel Approach to Biomarker Discovery. Cold Spring Harb. Symp. Quant. Biol. 73, 389–401. doi: 10.1101/sqb.2008.73.021.

Mann, K., Deny, S., Ganguli, S., and Clandinin, T. R. (2021). Coupling of activity, metabolism and behaviour across the Drosophila brain. Nature 593, 244–248. doi: 10.1038/s41586-021-03497-0.

Mantle, D., Heaton, R. A., and Hargreaves, I. P. (2021). Coenzyme Q10, Ageing and the Nervous System: An Overview. Antioxidants 11, 2. doi: 10.3390/antiox11010002.

March-Diaz, R., Lara-Ureña, N., Romero-Molina, C., Heras-Garvin, A., Ortega-de San Luis, C., Alvarez-Vergara, M. I., et al. (2021). Hypoxia compromises the mitochondrial metabolism of Alzheimer’s disease microglia via HIF1. *Nat*. Aging 1, 385–399. doi: 10.1038/s43587-021-00054-2.

Mattson, M. P., and Arumugam, T. V. (2018). Hallmarks of Brain Aging: Adaptive and Pathological Modification by Metabolic States. Cell Metab. 27, 1176–1199. doi: 10.1016/j.cmet.2018.05.011.

McBean, G. (2017). Cysteine, Glutathione, and Thiol Redox Balance in Astrocytes. Antioxidants 6, 62. doi: 10.3390/antiox6030062.

McCormack, J. G., and Denton, R. M. (1979). The effects of calcium ions and adenine nucleotides on the activity of pig heart 2-oxoglutarate dehydrogenase complex. Biochem. J. 180, 533–544. doi: 10.1042/bj1800533.

McGettrick, A. F., and O’Neill, L. A. J. (2020). The Role of HIF in Immunity and Inflammation. Cell Metab. 32, 524–536. doi: 10.1016/j.cmet.2020.08.002.

Meidenbauer, J. J., Ta, N., and Seyfried, T. N. (2014). Influence of a ketogenic diet, fish-oil, and calorie restriction on plasma metabolites and lipids in C57BL/6J mice. Nutr. Metab. 11, 23. doi: 10.1186/1743-7075-11-23.

Menahan, L. A., Hron, W. T., Hinkelman, D. G., and Miziorko, H. M. (1981). Interrelationships between 3-Hydroxy-3-Methylglutaryl-CoA Synthase, Acetoacetyl-CoA and Ketogenesis. Eur. J. Biochem. 119, 287–294. doi: 10.1111/j.1432-1033.1981.tb05606.x.

Meyer, D. J., Díaz-García, C. M., Nathwani, N., Rahman, M., and Yellen, G. (2022). The Na+/K+ pump dominates control of glycolysis in hippocampal dentate granule cells. eLife 11, e81645. doi: 10.7554/eLife.81645.

Milne, J. C., Lambert, P. D., Schenk, S., Carney, D. P., Smith, J. J., Gagne, D. J., et al. (2007). Small molecule activators of SIRT1 as therapeutics for the treatment of type 2 diabetes. Nature 450, 712–716. doi: 10.1038/nature06261.

Mink, J. W., Blumenschine, R. J., and Adams, D. B. (1981). Ratio of central nervous system to body metabolism in vertebrates: its constancy and functional basis. Am. J. Physiol. 241, R203–212. doi: 10.1152/ajpregu.1981.241.3.R203.

Mironov, S. L. (2007). ADP Regulates Movements of Mitochondria in Neurons. Biophys. J. 92, 2944–2952. doi: 10.1529/biophysj.106.092981.

Mogilevskaya, E., Demin, O., and Goryanin, I. (2006). Kinetic model of mitochondrial Krebs cycle: unraveling the mechanism of salicylate hepatotoxic effects. J. Biol. Phys. 32, 245–271. doi: 10.1007/s10867-006-9015-y.

Mongeon, R., Venkatachalam, V., and Yellen, G. (2016). Cytosolic NADH-NAD+ Redox Visualized in Brain Slices by Two-Photon Fluorescence Lifetime Biosensor Imaging. Antioxid. Redox Signal. 25, 553–563. doi: 10.1089/ars.2015.6593.

Mormino, A., Cocozza, G., Fontemaggi, G., Valente, S., Esposito, V., Santoro, A., et al. (2021). Histone-deacetylase 8 drives the immune response and the growth of glioma. Glia 69, 2682–2698. doi: 10.1002/glia.24065.

Mueggler, P. A., and Wolfe, R. G. (1978). Malate dehydrogenase. Kinetic studies of substrate activation of supernatant enzyme by L-malate. Biochemistry 17, 4615–4620. doi: 10.1021/bi00615a006.

Mulukutla, B. C., Yongky, A., Daoutidis, P., and Hu, W.-S. (2014). Bistability in Glycolysis Pathway as a Physiological Switch in Energy Metabolism. PLoS ONE 9, e98756. doi: 10.1371/journal.pone.0098756.

Mulukutla, B. C., Yongky, A., Grimm, S., Daoutidis, P., and Hu, W.-S. (2015). Multiplicity of Steady States in Glycolysis and Shift of Metabolic State in Cultured Mammalian Cells. PLOS ONE 10, e0121561. doi: 10.1371/journal.pone.0121561.

Muraleedharan, R., Gawali, M. V., Tiwari, D., Sukumaran, A., Oatman, N., Anderson, J., et al. (2020). AMPK-Regulated Astrocytic Lactate Shuttle Plays a Non-Cell-Autonomous Role in Neuronal Survival. Cell Rep. 32, 108092. doi: 10.1016/j.celrep.2020.108092.

Nazaret, C., Heiske, M., Thurley, K., and Mazat, J.-P. (2009). Mitochondrial energetic metabolism: A simplified model of TCA cycle with ATP production. J. Theor. Biol. 258, 455–464. doi: 10.1016/j.jtbi.2008.09.037.

Nehlig, A. (2004). Brain uptake and metabolism of ketone bodies in animal models. Prostaglandins Leukot. Essent. Fatty Acids 70, 265–275. doi: 10.1016/j.plefa.2003.07.006.

Neves, A., Costalat, R., and Pellerin, L. (2012). Determinants of brain cell metabolic phenotypes and energy substrate utilization unraveled with a modeling approach. PLoS Comput. Biol. 8, e1002686. doi: 10.1371/journal.pcbi.1002686.

Neves, S. R. (2011). Obtaining and Estimating Kinetic Parameters from the Literature. Sci. Signal. 4. doi: 10.1126/scisignal.2001988.

Ng, F., Wijaya, L., and Tang, B. L. (2015). SIRT1 in the brain-connections with aging-associated disorders and lifespan. Front. Cell. Neurosci. 9, 64. doi: 10.3389/fncel.2015.00064.

Niccoli, T., and Partridge, L. (2012). Ageing as a Risk Factor for Disease. Curr. Biol. 22, R741–R752. doi: 10.1016/j.cub.2012.07.024.

Nielsen, N. C., Zahler, W. L., and Fleischer, S. (1973). Mitochondrial D- -hydroxybutyrate dehydrogenase. IV. Kinetic analysis of reaction mechanism. J. Biol. Chem. 248, 2556–2562.

Nishihara, E., Moriya, T., and Shinohara, K. (2007). Expression of steroid receptor coactivator-1 is elevated during neuronal differentiation of murine neural stem cells. Brain Res. 1135, 22–30. doi: 10.1016/j.brainres.2006.12.026.

Niven, J. E. (2016). Neuronal energy consumption: biophysics, efficiency and evolution. Curr. Opin. Neurobiol. 41, 129–135. doi: 10.1016/j.conb.2016.09.004.

Oka, M., Suzuki, E., Asada, A., Saito, T., Iijima, K. M., and Ando, K. (2021). Increasing neuronal glucose uptake attenuates brain aging and promotes life span under dietary restriction in Drosophila. iScience 24, 101979. doi: 10.1016/j.isci.2020.101979.

O’Neill, L. A. J., and Hardie, D. G. (2013). Metabolism of inflammation limited by AMPK and pseudo-starvation. Nature 493, 346–355. doi: 10.1038/nature11862.

Orosz, F., Wágner, G., Ortega, F., Cascante, M., and Ovádi, J. (2003). Glucose conversion by multiple pathways in brain extract: theoretical and experimental analysis. Biochem. Biophys. Res. Commun. 309, 792–797. doi: 10.1016/j.bbrc.2003.08.072.

Øyehaug, L., Østby, I., Lloyd, C. M., Omholt, S. W., and Einevoll, G. T. (2012). Dependence of spontaneous neuronal firing and depolarisation block on astroglial membrane transport mechanisms. J. Comput. Neurosci. 32, 147–165. doi: 10.1007/s10827-011-0345-9.

Palla, A. R., Ravichandran, M., Wang, Y. X., Alexandrova, L., Yang, A. V., Kraft, P., et al. (2021). Inhibition of prostaglandin-degrading enzyme 15-PGDH rejuvenates aged muscle mass and strength. Science 371, eabc8059. doi: 10.1126/science.abc8059.

Pamiljans, V., Krishnaswamy, P. R., Dumville, G., and Meister, A. (1962). Studies on the Mechanism of Glutamine Synthesis; Isolation and Properties of the Enzyme from Sheep Brain. Biochemistry 1, 153–158. doi: 10.1021/bi00907a023.

Park, J. O., Rubin, S. A., Xu, Y.-F., Amador-Noguez, D., Fan, J., Shlomi, T., et al. (2016). Metabolite concentrations, fluxes and free energies imply efficient enzyme usage. Nat. Chem. Biol. 12, 482–489. doi: 10.1038/nchembio.2077.

Pathak, D., Shields, L. Y., Mendelsohn, B. A., Haddad, D., Lin, W., Gerencser, A. A., et al. (2015). The Role of Mitochondrially Derived ATP in Synaptic Vesicle Recycling. J. Biol. Chem. 290, 22325–22336. doi: 10.1074/jbc.M115.656405.

Pérez-Escuredo, J., Van Hée, V. F., Sboarina, M., Falces, J., Payen, V. L., Pellerin, L., et al. (2016). Monocarboxylate transporters in the brain and in cancer. Biochim. Biophys. Acta BBA - Mol. Cell Res. 1863, 2481–2497. doi: 10.1016/j.bbamcr.2016.03.013.

Pochini, L., Scalise, M., Galluccio, M., and Indiveri, C. (2014). Membrane transporters for the special amino acid glutamine: structure/function relationships and relevance to human health. Front. Chem. 2. doi: 10.3389/fchem.2014.00061.

Poliquin, P. O., Chen, J., Cloutier, M., Trudeau, L.-É., and Jolicoeur, M. (2013). Metabolomics and In-Silico Analysis Reveal Critical Energy Deregulations in Animal Models of Parkinson’s Disease. PLoS ONE 8, e69146. doi: 10.1371/journal.pone.0069146.

Pospischil, M., Toledo-Rodriguez, M., Monier, C., Piwkowska, Z., Bal, T., Frégnac, Y., et al. (2008). Minimal Hodgkin–Huxley type models for different classes of cortical and thalamic neurons. Biol. Cybern. 99, 427–441. doi: 10.1007/s00422-008-0263-8.

Power, J. M., Wu, W. W., Sametsky, E., Oh, M. M., and Disterhoft, J. F. (2002). Age-related enhancement of the slow outward calcium-activated potassium current in hippocampal CA1 pyramidal neurons in vitro. J. Neurosci. Off. J. Soc. Neurosci. 22, 7234–7243. doi: 10.1523/JNEUROSCI.22-16-07234.2002.

Pritchard, J. B. (1995). Intracellular alpha-ketoglutarate controls the efficacy of renal organic anion transport. J. Pharmacol. Exp. Ther. 274, 1278–1284.

Rackauckas, C., and Nie, Q. (2017). DifferentialEquations.jl – A Performant and Feature-Rich Ecosystem for Solving Differential Equations in Julia. J. Open Res. Softw. 5, 15. doi: 10.5334/jors.151.

Recasens, M., Benezra, R., Basset, P., and Mandel, P. (1980). Cysteine sulfinate aminotransferase and aspartate aminotransferase isoenzymes of rat brain. Purification, characterization, and further evidence for identity. Biochemistry 19, 4583–4589. doi: 10.1021/bi00561a007.

Reimann, M. W., Nolte, M., Scolamiero, M., Turner, K., Perin, R., Chindemi, G., et al. (2017). Cliques of Neurons Bound into Cavities Provide a Missing Link between Structure and Function. Front. Comput. Neurosci. 11, 48. doi: 10.3389/fncom.2017.00048.

Riddle, D. R. ed. (2007). Brain Aging: Models, Methods, and Mechanisms. Boca Raton (FL): CRC Press/Taylor & Francis Available at: http://www.ncbi.nlm.nih.gov/books/NBK1834/ [Accessed August 29, 2023].

Rizzo, V., Richman, J., and Puthanveettil, S. V. (2015). Dissecting mechanisms of brain aging by studying the intrinsic excitability of neurons. Front. Aging Neurosci. 6. doi: 10.3389/fnagi.2014.00337.

Roberg, B., Torgner, I. A., and Kvamme, E. (1999). Inhibition of glutamine transport in rat brain mitochondria by some amino acids and tricarboxylic acid cycle intermediates. Neurochem. Res. 24, 809–814. doi: 10.1023/a:1020941510764.

Robinson, M. B., and Jackson, J. G. (2016). Astroglial glutamate transporters coordinate excitatory signaling and brain energetics. Neurochem. Int. 98, 56–71. doi: 10.1016/j.neuint.2016.03.014.

Rock, C. O., Calder, R. B., Karim, M. A., and Jackowski, S. (2000). Pantothenate Kinase Regulation of the Intracellular Concentration of Coenzyme A. J. Biol. Chem. 275, 1377–1383. doi: 10.1074/jbc.275.2.1377.

Rodgers, J. T., Lerin, C., Haas, W., Gygi, S. P., Spiegelman, B. M., and Puigserver, P. (2005). Nutrient control of glucose homeostasis through a complex of PGC-1alpha and SIRT1. Nature 434, 113–118. doi: 10.1038/nature03354.

Roeder, L. M., Poduslo, S. E., and Tildon, J. T. (1982). Utilization of ketone bodies and glucose by established neural cell lines. J. Neurosci. Res. 8, 671–682. doi: 10.1002/jnr.490080412.

Rolfe, D. F., and Brown, G. C. (1997). Cellular energy utilization and molecular origin of standard metabolic rate in mammals. Physiol. Rev. 77, 731–758. doi: 10.1152/physrev.1997.77.3.731.

Ronowska, A., Szutowicz, A., Bielarczyk, H., Gul-Hinc, S., Klimaszewska-Łata, J., Dyś, A., et al. (2018). The Regulatory Effects of Acetyl-CoA Distribution in the Healthy and Diseased Brain. Front. Cell. Neurosci. 12. doi: 10.3389/fncel.2018.00169.

Ross, J. M., Oberg, J., Brene, S., Coppotelli, G., Terzioglu, M., Pernold, K., et al. (2010). High brain lactate is a hallmark of aging and caused by a shift in the lactate dehydrogenase A/B ratio. Proc. Natl. Acad. Sci. 107, 20087–20092. doi: 10.1073/pnas.1008189107.

Rybalkin, S. D., Hinds, T. R., and Beavo, J. A. (2013). “Enzyme Assays for cGMP Hydrolyzing Phosphodiesterases,” in Guanylate Cyclase and Cyclic GMP Methods in Molecular Biology., eds. T. Krieg and R. Lukowski (Totowa, NJ: Humana Press), 51–62. doi: 10.1007/978-1-62703-459-3_3.

Satoh, A., Imai, S., and Guarente, L. (2017). The brain, sirtuins, and ageing. Nat. Rev. Neurosci. 18, 362–374. doi: 10.1038/nrn.2017.42.

Savtchenko, L. P., Bard, L., Jensen, T. P., Reynolds, J. P., Kraev, I., Medvedev, N., et al. (2018). Disentangling astroglial physiology with a realistic cell model in silico. Nat. Commun. 9. doi: 10.1038/s41467-018-05896-w.

Schaum, N., Lehallier, B., Hahn, O., Pálovics, R., Hosseinzadeh, S., Lee, S. E., et al. (2020). Ageing hallmarks exhibit organ-specific temporal signatures. Nature 583, 596–602. doi: 10.1038/s41586-020-2499-y.

Schousboe, A., Waagepetersen, H. S., and Sonnewald, U. (2019). Astrocytic pyruvate carboxylation: Status after 35 years. J. Neurosci. Res. 97, 890–896. doi: 10.1002/jnr.24402.

Sedlak, T. W., Paul, B. D., Parker, G. M., Hester, L. D., Snowman, A. M., Taniguchi, Y., et al. (2019). The glutathione cycle shapes synaptic glutamate activity. Proc. Natl. Acad. Sci. 116, 2701–2706. doi: 10.1073/pnas.1817885116.

Seelig, M. S., and Preuss, H. G. (1994). Magnesium metabolism and perturbations in the elderly. Geriatr. Nephrol. Urol. 4, 101–111. doi: 10.1007/BF01436050.

Sharma, H. K., and Rothstein, M. (1984). Altered brain phosphoglycerate kinase from aging rats. Mech. Ageing Dev. 25, 285–296. doi: 10.1016/0047-6374(84)90002-2.

Shen, P., Xu, A., Hou, Y., Wang, H., Gao, C., He, F., et al. (2021). Conserved paradoxical relationships among the evolutionary, structural and expressional features of KRAB zinc-finger proteins reveal their special functional characteristics. BMC Mol. Cell Biol. 22, 7. doi: 10.1186/s12860-021-00346-w.

Shestov, A. A., Valette, J., Uğurbil, K., and Henry, P.-G. (2007). On the reliability of13C metabolic modeling with two-compartment neuronal-glial models. J. Neurosci. Res. 85, 3294–3303. doi: 10.1002/jnr.21269.

Shichkova, P., Coggan, J. S., Markram, H., and Keller, D. (2021). A Standardized Brain Molecular Atlas: A Resource for Systems Modeling and Simulation. Front. Mol. Neurosci. 14, 604559. doi: 10.3389/fnmol.2021.604559.

Shin, H.-J. R., Kim, H., Oh, S., Lee, J.-G., Kee, M., Ko, H.-J., et al. (2016). AMPK-SKP2-CARM1 signalling cascade in transcriptional regulation of autophagy. Nature 534, 553–557. doi: 10.1038/nature18014.

Simpson, I. A., Carruthers, A., and Vannucci, S. J. (2007). Supply and Demand in Cerebral Energy Metabolism: The Role of Nutrient Transporters. J. Cereb. Blood Flow Metab. 27, 1766–1791. doi: 10.1038/sj.jcbfm.9600521.

Singh, A., D’Amico, D., Andreux, P. A., Fouassier, A. M., Blanco-Bose, W., Evans, M., et al. (2022). Urolithin A improves muscle strength, exercise performance, and biomarkers of mitochondrial health in a randomized trial in middle-aged adults. Cell Rep. Med. 3, 100633. doi: 10.1016/j.xcrm.2022.100633.

Sizemore, A. E., Giusti, C., Kahn, A., Vettel, J. M., Betzel, R. F., and Bassett, D. S. (2018). Cliques and cavities in the human connectome. J. Comput. Neurosci. 44, 115–145. doi: 10.1007/s10827-017-0672-6.

Smith, C. M., Bryla, J., and Williamson, J. R. (1974). Regulation of mitochondrial alpha-ketoglutarate metabolism by product inhibition at alpha-ketoglutarate dehydrogenase. J. Biol. Chem. 249, 1497–1505.

Smith, R. N., Agharkar, A. S., and Gonzales, E. B. (2014). A review of creatine supplementation in age-related diseases: more than a supplement for athletes. F1000Research 3, 222. doi: 10.12688/f1000research.5218.1.

Smithers, H. E., Terry, J. R., Brown, J. T., and Randall, A. D. (2017). Aging-Associated Changes to Intrinsic Neuronal Excitability in the Bed Nucleus of the Stria Terminalis Is Cell Type-Dependent. Front. Aging Neurosci. 9, 424. doi: 10.3389/fnagi.2017.00424.

Sokoloff, L. (1996). “Cerebral Metabolism and Visualization of Cerebral Activity,” in Comprehensive Human Physiology, eds. R. Greger and U. Windhorst (Berlin, Heidelberg: Springer Berlin Heidelberg), 579–602. doi: 10.1007/978-3-642-60946-6_30.

Stillman, C. M., Esteban-Cornejo, I., Brown, B., Bender, C. M., and Erickson, K. I. (2020). Effects of Exercise on Brain and Cognition Across Age Groups and Health States. Trends Neurosci. 43, 533–543. doi: 10.1016/j.tins.2020.04.010.

Sugrue, M. M., and Tatton, W. G. (2001). Mitochondrial Membrane Potential in Aging Cells. Neurosignals 10, 176–188. doi: 10.1159/000046886.

Sun, X., He, G., Qing, H., Zhou, W., Dobie, F., Cai, F., et al. (2006). Hypoxia facilitates Alzheimer’s disease pathogenesis by up-regulating BACE1 gene expression. Proc. Natl. Acad. Sci. U. S. A. 103, 18727–18732. doi: 10.1073/pnas.0606298103.

Sun, Z., and Xu, Y. (2020). Nuclear Receptor Coactivators (NCOAs) and Corepressors (NCORs) in the Brain. Endocrinology 161, bqaa083. doi: 10.1210/endocr/bqaa083.

Suresh, S. N., Chavalmane, A. K., Pillai, M., Ammanathan, V., Vidyadhara, D. J., Yarreiphang, H., et al. (2018). Modulation of Autophagy by a Small Molecule Inverse Agonist of ERRα Is Neuroprotective. Front. Mol. Neurosci. 11, 109. doi: 10.3389/fnmol.2018.00109.

Szklarczyk, D., Gable, A. L., Lyon, D., Junge, A., Wyder, S., Huerta-Cepas, J., et al. (2019). STRING v11: protein–protein association networks with increased coverage, supporting functional discovery in genome-wide experimental datasets. Nucleic Acids Res. 47, D607–D613. doi: 10.1093/nar/gky1131.

Takahashi, H., Manaka, S., and Sano, K. (1981). Changes in extracellular potassium concentration in cortex and brain stem during the acute phase of experimental closed head injury. J. Neurosurg. 55, 708–717. doi: 10.3171/jns.1981.55.5.0708.

Takeda, M., Briggs, L. E., Wakimoto, H., Marks, M. H., Warren, S. A., Lu, J. T., et al. (2009). Slow progressive conduction and contraction defects in loss of Nkx2-5 mice after cardiomyocyte terminal differentiation. Lab. Investig. J. Tech. Methods Pathol. 89, 983–993. doi: 10.1038/labinvest.2009.59.

Tantama, M., Martínez-François, J. R., Mongeon, R., and Yellen, G. (2013). Imaging energy status in live cells with a fluorescent biosensor of the intracellular ATP-to-ADP ratio. Nat. Commun. 4, 2550. doi: 10.1038/ncomms3550.

Taylor, C. T., and Scholz, C. C. (2022). The effect of HIF on metabolism and immunity. Nat. Rev. Nephrol. 18, 573–587. doi: 10.1038/s41581-022-00587-8.

Theurey, P., Connolly, N. M. C., Fortunati, I., Basso, E., Lauwen, S., Ferrante, C., et al. (2019). Systems biology identifies preserved integrity but impaired metabolism of mitochondria due to a glycolytic defect in Alzheimer’s disease neurons. Aging Cell 18, e12924. doi: 10.1111/acel.12924.

Thibonnier, M., Esau, C., Ghosh, S., Wargent, E., and Stocker, C. (2020). Metabolic and energetic benefits of microRNA-22 inhibition. BMJ Open Diabetes Res. Care 8, e001478. doi: 10.1136/bmjdrc-2020-001478.

Thiepold, A.-L., Lorenz, N. I., Foltyn, M., Engel, A. L., Divé, I., Urban, H., et al. (2017). Mammalian target of rapamycin complex 1 activation sensitizes human glioma cells to hypoxia-induced cell death. Brain J. Neurol. 140, 2623–2638. doi: 10.1093/brain/awx196.

Tiveci, S., Akın, A., Çakır, T., Saybaşılı, H., and Ülgen, K. (2005). Modelling of calcium dynamics in brain energy metabolism and Alzheimer’s disease. Comput. Biol. Chem. 29, 151–162. doi: 10.1016/j.compbiolchem.2005.03.002.

Tong, J., Fitzmaurice, P. S., Moszczynska, A., Mattina, K., Ang, L.-C., Boileau, I., et al. (2016). Do glutathione levels decline in aging human brain? Free Radic. Biol. Med. 93, 110–117. doi: 10.1016/j.freeradbiomed.2016.01.029.

Tretter, L., Patocs, A., and Chinopoulos, C. (2016). Succinate, an intermediate in metabolism, signal transduction, ROS, hypoxia, and tumorigenesis. Biochim. Biophys. Acta 1857, 1086–1101. doi: 10.1016/j.bbabio.2016.03.012.

Tripathi, M., Yen, P. M., and Singh, B. K. (2020). Estrogen-Related Receptor Alpha: An Under-Appreciated Potential Target for the Treatment of Metabolic Diseases. Int. J. Mol. Sci. 21, 1645. doi: 10.3390/ijms21051645.

Tsuboi, K. K., Fukunaga, K., and Petricciani, J. C. (1969). Purification and specific kinetic properties of erythrocyte uridine diphosphate glucose pyrophosphorylase. J. Biol. Chem. 244, 1008–1015.

Vali, S., Mythri, R. B., Jagatha, B., Padiadpu, J., Ramanujan, K. S., Andersen, J. K., et al. (2007). Integrating glutathione metabolism and mitochondrial dysfunction with implications for Parkinson’s disease: a dynamic model. Neuroscience 149, 917–930. doi: 10.1016/j.neuroscience.2007.08.028.

Verkhratsky, A., and Nedergaard, M. (2018). Physiology of Astroglia. Physiol. Rev. 98, 239–389. doi: 10.1152/physrev.00042.2016.

Vitale, P., Salgueiro-Pereira, A. R., Lupascu, C. A., Willem, M., Migliore, R., Migliore, M., et al. (2021). Analysis of Age-Dependent Alterations in Excitability Properties of CA1 Pyramidal Neurons in an APPPS1 Model of Alzheimer’s Disease. Front. Aging Neurosci. 13, 668948. doi: 10.3389/fnagi.2021.668948.

Volkova, M., Garg, R., Dick, S., and Boheler, K. R. (2005). Aging-associated changes in cardiac gene expression. Cardiovasc. Res. 66, 194–204. doi: 10.1016/j.cardiores.2004.11.016.

Waitt, A. E., Reed, L., Ransom, B. R., and Brown, A. M. (2017). Emerging Roles for Glycogen in the CNS. Front. Mol. Neurosci. 10. doi: 10.3389/fnmol.2017.00073.

Weber, B., and Barros, L. F. (2015). The Astrocyte: Powerhouse and Recycling Center. Cold Spring Harb. Perspect. Biol. 7, a020396. doi: 10.1101/cshperspect.a020396.

White, H., and Jencks, W. P. (1976). Mechanism and specificity of succinyl-CoA:3-ketoacid coenzyme A transferase. J. Biol. Chem. 251, 1688–1699.

Wilcock, A. R., Sharpe, D. M., and Goldberg, D. M. (1973). Kinetic similarity of enzymes in human blood serum and cerebrospinal fluid: Aspartate aminotransferase and lactate dehydrogenase. J. Neurol. Sci. 20, 97–101. doi: 10.1016/0022-510X(73)90121-4.

Williamson, D., Lund, P., and Krebs, H. (1967). The redox state of free nicotinamide-adenine dinucleotide in the cytoplasm and mitochondria of rat liver. Biochem. J. 103, 514–527. doi: 10.1042/bj1030514.

Winter, F., Bludszuweit-Philipp, C., and Wolkenhauer, O. (2018). Mathematical analysis of the influence of brain metabolism on the BOLD signal in Alzheimer’s disease. J. Cereb. Blood Flow Metab. 38, 304–316. doi: 10.1177/0271678X17693024.

Witthoft, A., Filosa, J. A., and Karniadakis, G. E. (2013). Potassium Buffering in the Neurovascular Unit: Models and Sensitivity Analysis. Biophys. J. 105, 2046–2054. doi: 10.1016/j.bpj.2013.09.012.

Wittig, U., Rey, M., Weidemann, A., Kania, R., and Müller, W. (2018). SABIO-RK: an updated resource for manually curated biochemical reaction kinetics. Nucleic Acids Res. 46, D656–D660. doi: 10.1093/nar/gkx1065.

Wu, F., Yang, F., Vinnakota, K. C., and Beard, D. A. (2007). Computer modeling of mitochondrial tricarboxylic acid cycle, oxidative phosphorylation, metabolite transport, and electrophysiology. J. Biol. Chem. 282, 24525–24537. doi: 10.1074/jbc.M701024200.

Xu, K., Morgan, K. T., Todd Gehris, A., Elston, T. C., and Gomez, S. M. (2011). A Whole-Body Model for Glycogen Regulation Reveals a Critical Role for Substrate Cycling in Maintaining Blood Glucose Homeostasis. PLoS Comput. Biol. 7, e1002272. doi: 10.1371/journal.pcbi.1002272.

Xue, X., Liu, B., Hu, J., Bian, X., and Lou, S. (2022). The potential mechanisms of lactate in mediating exercise-enhanced cognitive function: a dual role as an energy supply substrate and a signaling molecule. Nutr. Metab. 19, 52. doi: 10.1186/s12986-022-00687-z.

Yamashita, D., Moriuchi, T., Osumi, T., and Hirose, F. (2016). Transcription Factor hDREF Is a Novel SUMO E3 Ligase of Mi2α. J. Biol. Chem. 291, 11619–11634. doi: 10.1074/jbc.M115.713370.

Yang, J.-S., Hsu, J.-W., Park, S.-Y., Li, J., Oldham, W. M., Beznoussenko, G. V., et al. (2018). GAPDH inhibits intracellular pathways during starvation for cellular energy homeostasis. Nature 561, 263–267. doi: 10.1038/s41586-018-0475-6.

Yang, Q., Zhao, W., Xing, Y., Li, P., Zhou, X., Ning, H., et al. (2021). Dysfunction of an energy sensor NFE2L1 triggers uncontrollable AMPK signal and glucose metabolism reprogramming. Molecular Biology doi: 10.1101/2021.09.07.459348.

Yang, S. Y., He, X. Y., and Schulz, H. (1987). Fatty acid oxidation in rat brain is limited by the low activity of 3-ketoacyl-coenzyme A thiolase. J. Biol. Chem. 262, 13027–13032.

Yi, G., and Grill, W. M. (2019). Average firing rate rather than temporal pattern determines metabolic cost of activity in thalamocortical relay neurons. Sci. Rep. 9. doi: 10.1038/s41598-019-43460-8.

Yoshino, J., Baur, J. A., and Imai, S.-I. (2018). NAD+ Intermediates: The Biology and Therapeutic Potential of NMN and NR. Cell Metab. 27, 513–528. doi: 10.1016/j.cmet.2017.11.002.

Yuk, J.-M., Kim, T. S., Kim, S. Y., Lee, H.-M., Han, J., Dufour, C. R., et al. (2015). Orphan Nuclear Receptor ERRα Controls Macrophage Metabolic Signaling and A20 Expression to Negatively Regulate TLR-Induced Inflammation. Immunity 43, 80–91. doi: 10.1016/j.immuni.2015.07.003.

Zhang, M. J., Pisco, A. O., Darmanis, S., and Zou, J. (2021a). Mouse aging cell atlas analysis reveals global and cell type-specific aging signatures. eLife 10, e62293. doi: 10.7554/eLife.62293.

Zhang, P., Qu, H.-Y., Wu, Z., Na, H., Hourihan, J., Zhang, F., et al. (2021b). ERK signaling licenses SKN-1A/NRF1 for proteasome production and proteasomal stress resistance. Cell Biology doi: 10.1101/2021.01.04.425272.

Zhang, X., Dash, R. K., Jacobs, E. R., Camara, A. K. S., Clough, A. V., and Audi, S. H. (2018). Integrated computational model of the bioenergetics of isolated lung mitochondria. PloS One 13, e0197921. doi: 10.1371/journal.pone.0197921.

Zhao, T., Kee, H. J., Bai, L., Kim, M.-K., Kee, S.-J., and Jeong, M. H. (2021). Selective HDAC8 Inhibition Attenuates Isoproterenol-Induced Cardiac Hypertrophy and Fibrosis via p38 MAPK Pathway. Front. Pharmacol. 12, 677757. doi: 10.3389/fphar.2021.677757.

Zhou, M., Xia, X., Yan, H., Li, S., Bian, S., Sha, X., et al. (2019). The Model of Aging Acceleration Network Reveals the Correlation of Alzheimer’s Disease and Aging at System Level. BioMed Res. Int. 2019, 4273108. doi: 10.1155/2019/4273108.

Zhu, F., Wang, R., Pan, X., and Zhu, Z. (2019). Energy expenditure computation of a single bursting neuron. Cogn. Neurodyn. 13, 75–87. doi: 10.1007/s11571-018-9503-3.

